# Experimental verification of the error minimization theory using non-standard genetic codes constructed in vitro

**DOI:** 10.64898/2026.02.24.707864

**Authors:** Ryota Miyachi, Norikazu Ichihashi

## Abstract

All living systems use an almost identical genetic code, the standard genetic code, in which 20 amino acids are assigned to 61 codons non-randomly. According to the error minimization theory, amino acids are arranged to minimize the mutational effect on protein function, while experimental verification remains limited. In this study, we constructed 10 non-standard genetic codes in vitro by reassigning three amino acids (Ala, Ser, and Leu) in vacant codons of the minimal genetic code, which consists of 21 tRNAs. Most of these non-standard genetic codes have a higher cost of amino acid replacement than the standard genetic code, calculated based on three amino acid properties: polar requirement (PR), molecular volume (MV), and hydropathy index (HI). The protein function of three reporter genes expressed using these non-standard genetic codes decreased similarly when random mutations were introduced into the genes, implying that the effect of mutations was similar across all the non-standard genetic codes tested here. This result provides direct experimental evidence that mutational robustness does not significantly change in individual reporter protein activity when the genetic code is altered within the range of mutational cost tested in this study (Cost_PR_: 5.29 – 5.77, Cost_MV_: 1848 – 2348, and Cost_HI_: 3.27 – 5.10), which covers approximately 18.4% (PR), 37.6% (MV), and 50.8% (HI) of possible cost range achievable among one million randomly-generated genetic codes.

## Introduction

The genetic code is the set of rules for the translation of nucleotide sequences in mRNA into amino acid sequences in proteins. All known living organisms use a nearly identical genetic code, referred to as the standard genetic code (SGC), representing a universal mechanism shared by almost all extant living systems on Earth (Crick, 1968; Koonin and Novozhilov, 2017; Woese, 1965; Woese et al., 1966). In SGC, 20 amino acids and stop codons are nonrandomly assigned to 64 codons; amino acids with similar physicochemical properties are assigned to neighboring codons (see Fig. 1A, SGC)(Koonin and Novozhilov, 2017, 2009). The origin and reason for this non-random codon organization remain a major mystery in biology(Crick, 1968; Di Giulio, 2005; Koonin and Novozhilov, 2017, 2009).

**Figure 1.**
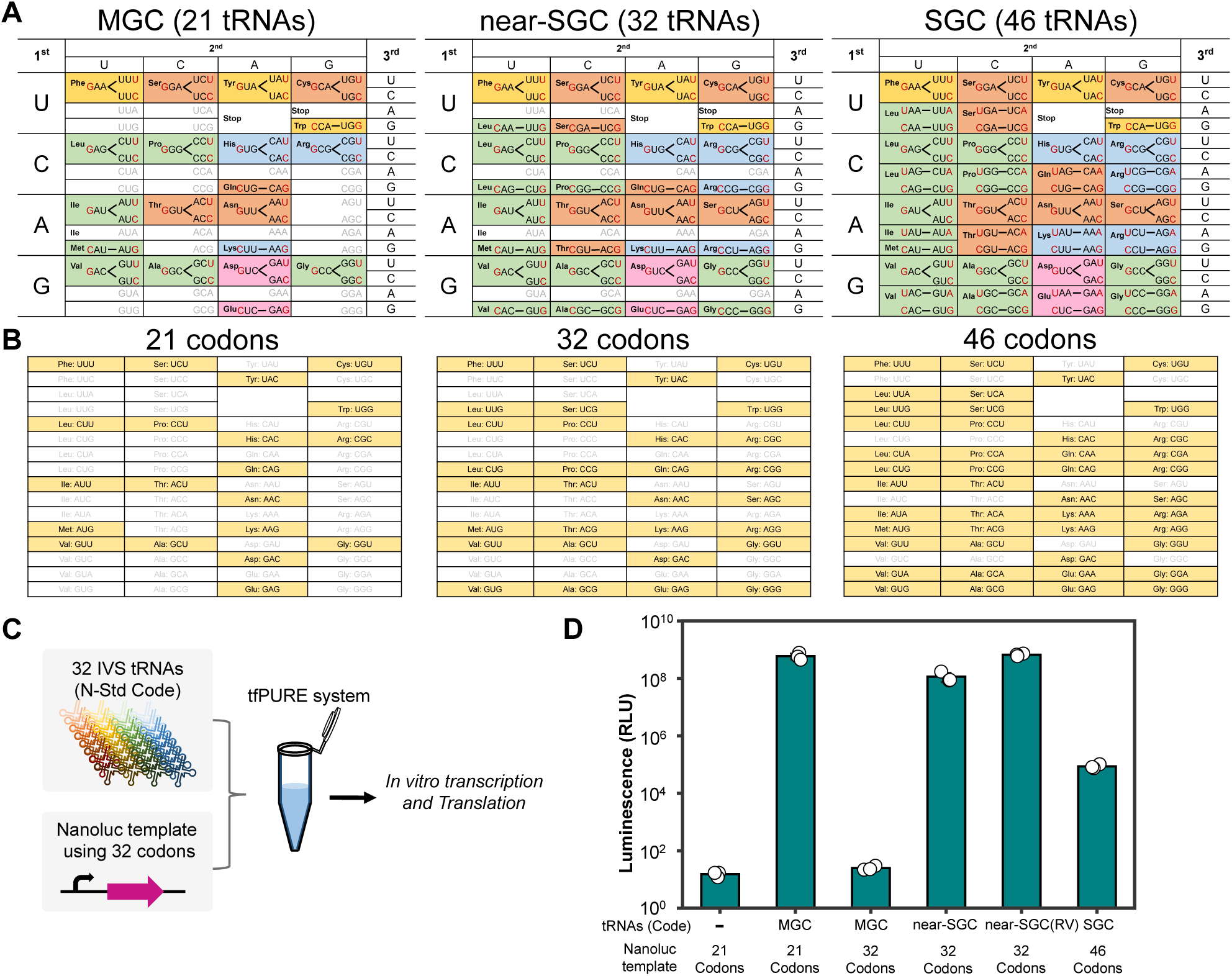
In vitro construction of minimal, near-standard, and standard genetic codes. **(A)** Anticodons of tRNAs and their corresponding codon assignments in the minimal genetic code (MGC), the near-standard genetic code (near-SGC), and the standard genetic code (SGC). Each codon is colored according to the physicochemical properties of the assigned amino acid: hydrophobic (green), aromatic (yellow), polar uncharged (orange), basic (blue), and acidic (pink). In each box, the anticodons of tRNAs (left) and the corresponding codons (right) are shown. (B) Codon sets used for reporter genes. The 21 codons contain only codons that are usable for MGC. Reporter genes composed of these codons were used in D and the subsequent experiments using non-standard genetic codes. The 32 and 46 codons contain codons that are usable for near-SGC and SGC, respectively. Reporter genes composed of these codons were used in D. (C) Schematic of the translation assay. Reporter genes (NanoLuc, 1 nM) consisting of the 21, 32, or 46-codon were translated in a customized reconstituted translation system lacking endogenous tRNAs (tRNA-free PURE system; tfPURE) supplemented with in vitro-synthesized tRNAs corresponding to MGC, near-SGC, or SGC (IPEN tRNA at 100 ng/µL; all other tRNAs at 12 ng/µL), and T7 RNA polymerase (0.42 U/µL) at 30 °C for 16 h. (D) NanoLuc activity after incubation. In near-SGC (RV), two tRNAs (tRNA^Val^ and tRNA^Arg^) were increased to 100 ng/µL. Each dot represents an independent experiment (n = 3). Bars indicate mean values, and error bars represent standard deviations. Statistical comparisons in (D) were performed using one-way ANOVA followed by Tukey’s post hoc test on NanoLuc activity; major comparisons are summarized in Table S8.

One explanation for this nonrandomness in SGC is mutational robustness. According to the error minimization theory, amino acid arrangement in SGC was selected to minimize the functional impact of amino acid substitutions caused by single-nucleotide mutations or translational errors(Freeland et al., 2003; Freeland and Hurst, 1998; Haig and Hurst, 1991). Several theoretical studies have estimated the possible cost of amino acid replacement (referred to as mutational cost in this study) based on the physicochemical properties of amino acids. These analyses consistently demonstrated that SGC exhibits an exceptionally low mutational cost compared with randomly generated genetic codes (Buhrman et al., 2013; Freeland and Hurst, 1998; Gilis et al., 2001; Goodarzi et al., 2004; Haig and Hurst, 1991; Omachi et al., 2023), although some genetic codes still have a lower cost than SGC(Błażej et al., 2018; Novozhilov et al., 2007; Wnȩtrzak et al., 2018). An alternative explanation for the non-randomness of SGC is a by-product of genetic code expansion(Massey, 2016, 2008). According to this explanation, the arrangement of SGC does not confer a selective advantage over other genetic codes. Despite these many theoretical studies, experimental verification is still limited.

A fundamental obstacle to the experimental verification of the role of genetic code arrangement in mutational robustness is the lack of variation in genetic codes in nature; all extant organisms use a nearly identical genetic code (Massey, 2015; Shulgina and Eddy, 2021). Recently, *E. coli* with designed genetic codes have been developed by removing specific stop codons and rarely used sense codons from the genome (Fredens et al., 2019; Lajoie et al., 2013; Nyerges et al., 2023; Robertson et al., 2025, 2021; Zürcher et al., 2022) and reassigning them to non-canonical or alternative amino acids through the introduction of orthogonal tRNA-aminoacyl-tRNA synthetase (aaRS) pairs (Costello et al., 2024; Shandell et al., 2021). However, these approaches based on living cells require extensive genome engineering, and thus, the number of codons that can be reassigned remains limited.

An alternative approach for genetic code alteration is in vitro reconstitution(Hibi et al., 2020; Miyachi et al., 2025; Shimizu et al., 2001). In this approach, the components required for translation, such as tRNAs and aaRSs, are individually prepared and supplied, allowing simultaneous reassignment of multiple codons. A representative platform for such reconstitution is the PURE system (Shimizu et al., 2001), a reconstituted cell-free translation system composed of purified translation components, including ribosomes, translation factors, aaRSs, amino acids, and energy-regeneration components. In particular, a tRNA-free PURE system (Miyachi et al., 2022), in which endogenous tRNA activity is minimized and defined tRNA sets are supplied externally, enables genetic codes to be reconstructed by controlling the supplied tRNAs. To date, several genetic codes have been reconstituted in vitro, including minimal genetic codes composed of 21 tRNAs that mimic or are derived from SGC, and those incorporating non-canonical amino acids(Forster et al., 2003; Iwane et al., 2016; Jones et al., 2025; McFeely et al., 2023). Translation of peptides and proteins using these reconstituted genetic codes has also been demonstrated (Calles et al., 2019; Fujino et al., 2020; Hibi et al., 2020; Iwane et al., 2016; Miyachi et al., 2025). However, reconstitution of elaborately arranged genetic codes and protein translation by them remains challenging.

In this study, to assess the effect of genetic code arrangement on mutational robustness, we constructed non-standard genetic codes in vitro and compared the effect of mutations on the function of translated proteins. We first established a minimal genetic code, composed of 21 tRNAs with vacant codons, which allows multiple alternative codon assignments to be introduced under otherwise comparable translation conditions. By introducing tRNA variants into these vacant codons, we then constructed 10 non-standard genetic codes with distinct costs of amino acid replacement, along with a near-standard genetic code. Using these genetic codes, we translated reporter gene libraries carrying random mutations and quantified the resulting protein activities. Random mutations decreased reporter protein function at similar levels across all genetic codes examined, implying that alterations of the genetic code within the ranges explored in this study have no significant effect on mutational robustness of individual protein activity.

## Results

### Construction of minimal and standard genetic codes

Before constructing non-standard genetic codes, we first constructed a minimal genetic code (MGC) using 21 tRNAs (20 for 20 amino acids and one for formyl-Met) according to a previously reported method (Fujino et al., 2020; Hibi et al., 2020)(Fig. 1A, left). In this MGC, we used tRNAs with G at the first position of the anticodon for 15 amino acids (Phe, Ser, Tyr, Cys, Leu, Pro, His, Arg, Ile, Thr, Asn, Val, Ala, Asp, and Gly), which can decode codons with C or U at the third position through wobble base pairing (Hibi et al., 2020). We also used tRNAs with C at the first position of the anticodon for 5 amino acids (Trp, Gln, Met, Lys, and Glu), which can decode codons with G at the third position. The expected correspondences between anticodons and the codons they decode are represented as lines in Fig. 1A (left three letters in each box indicate the anticodon of the tRNA, and the right ones indicate the codon).

To ensure fair comparisons, we also constructed genetic codes close to the SGC using the same method (i.e., using unmodified tRNAs). Previously, a genetic code approximating the SGC was constructed in vitro using 32 tRNAs for peptide synthesis (Iwane et al., 2016). Following this method, we introduced 11 additional tRNAs (Leu_CAA, Ser_CGA, Leu_CAG, Pro_CGG, Arg_CCG, Ser_GCU, Thr_CGU, Arg_CCU, Val_CAC, Ala_CGC, and Gly_CCC) into the MGC (Fig. 1A, middle). We refer to this as the near-standard genetic code (near-SGC). Although the near-SGC mimics the SGC in most of the codons, it does not cover codons with A at the third nucleotide position (NNA). We therefore further introduced 14 tRNAs with UNN anticodons that are expected to decode NNA codons to construct SGC consisting of a 46-tRNA set (Fig. 1A, right).

We next examined the translation efficiency of the reconstructed MGC, near-SGC, and SGC. Translation was performed in a custom-made tRNA-free PURE system (tfPURE) (Miyachi et al., 2025), supplemented with 21 tRNAs for MGC, 32 tRNAs for near-SGC, or 46 tRNAs for SGC. As a reporter protein, we used the NanoLuc luciferase, originally derived from deep-sea shrimp (Hall et al., 2012). For MGC, the NanoLuc sequence was designed to use only the 21 corresponding codons (Fig. 1B, 21codons). For near-SGC and SGC, the NanoLuc coding sequences were designed so that the codons available in each genetic code were used with minimal differences in codon counts, while preserving the amino acid sequence (Fig. 1B, 32 codons and 46 codons). After incubation at 30 °C for 16 hours, luminescence was measured as an indicator of translation efficiency. Translation using near-SGC exhibited lower luminescence (1.2×10^8^; Fig. 1D, near-SGC) than that using MGC (6.0×10^8^; MGC, 21 codons). To improve translation efficiency with near-SGC, we focused on two tRNA concentrations (tRNA^Val^_CAC_ and tRNA^Arg^_CCU_), which were suggested to have low activities in a previous study (Iwane et al., 2016). Upon increasing the concentrations of these two tRNAs, the translation increased up to 100 ng/μL (Fig. S1). At 100 ng/μL for each tRNA, the translation efficiency of near-SGC was increased to a level comparable to that of MGC (6.6×10^8^; Fig. 1D, near-Std (RV)). The increased concentrations of these two tRNAs were used in all the subsequent experiments. The translation level of SGC, which uses 46 codons with 46 tRNAs, was approximately 100-fold lower (8.6×10^4^; Fig. 1D), indicating that the additional tRNAs corresponding to NNA codons may not function as efficiently as other tRNAs. This inefficiency is probably due to the lack of chemical modifications in tRNAs used here, as the anticodon of the native tRNAs that decode NNA are frequently modified in the cell (El Yacoubi et al., 2012; Masuda and Hou, 2024). Such chemical modifications may be required for efficient translation of the NNA codon (Cui et al., 2015; Madore et al., 1999; Tamura et al., 1992).

We also examined whether the lower translation efficiency of the 46-codon NanoLuc template could be explained by sequence-dependent effects, such as codon context or mRNA structure. When the 21-, 32-, and 46-codon NanoLuc templates were translated using native *E. coli* tRNAs in the tfPURE system (Fig. S2), the 46-codon template showed lower activity than the 21- and 32-codon templates; however, this difference was within approximately two-fold. Accordingly, we decided to use only the 32 codons used in near-SGC (i.e., excluding NNA codons) in the subsequent construction of non-standard genetic codes.

### Test of the availability of each vacant codon for Ala, Ser, and Leu

We next aimed to construct multiple non-standard genetic codes (nonSGCs) by introducing tRNAs with altered anticodon sequences into the vacant codons of MGC, excluding NNA codons. Currently, anticodon sequences of tRNAs can be altered only for Ala, Ser, and Leu while maintaining recognition by their respective aminoacyl-tRNA synthetases (aaRSs) because the aaRSs for these amino acids do not recognize the anticodon sequence (Asahara et al., 1993; Francklyn and Schimmel, 1989; Sampson and Saks, 1993). Therefore, we decided to design nonSGCs by reassigning Ala, Ser, and Leu to 10 vacant codons (UUG, UCG, CUG, CCG, CGG, ACG, AGC, GUG, GCG, and GGG) in MGC. Note that one vacant codon (AGG) was assigned to the original amino acid (Arg) because this codon is already distant from the codon used for Arg (CGC) in MGC and does not need to be reassigned.

We tested whether each of the 10 vacant codons is usable for Ala, Leu, and Ser when the corresponding tRNA variant with an altered anticodon is introduced (Fig. 2A). To test this, we constructed a total of 25 genetic codes (nine for Ala, eight for Ser, and eight for Leu) by introducing one tRNA variant with an altered anticodon into MGC (referred to as 21 + 1) and also designed 25 corresponding NanoLuc template DNAs, in which some codons for Ala, Ser, and Leu (two for Ala, three for Ser, and four for Leu) were replaced with the corresponding vacant codons from the 21-codon set shown in Fig. 1B. For example, when assigning Ala to the UUG codon (Fig. 2A, bottom), two GCU codons used for Ala in the template DNA were changed to UUG codons.

**Figure 2.**
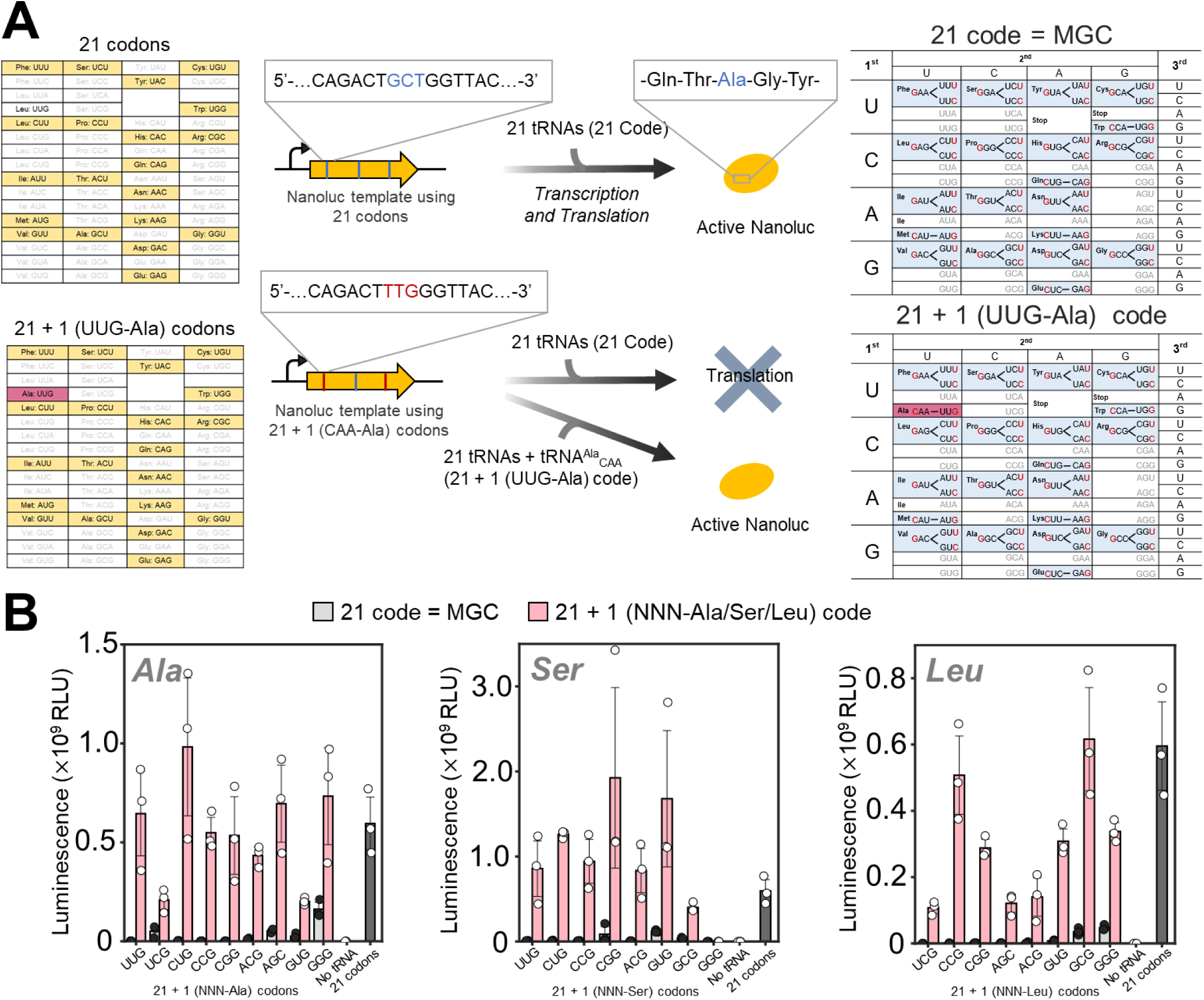
Reassignment experiments to test the availability of 10 vacant codons for Ala, Ser, and Leu. (A) Schematic illustration of reassignment experiments. Translation with the original MGC and NanoLuc template is shown at the top for comparison. An example of Ala reassignment to the UUG codon is shown at the bottom. In this example, three Ala codons in the NanoLuc sequence were replaced with one type of vacant codon (e.g., UUG), generating a 21 + 1 (UUG-Ala) codon set. Similar reassignment experiments were performed for three amino acids (Ala, Ser, and Leu) and nine vacant codons. Specifically, two Ala codons (Ala16 and Ala120), three Ser codons (Ser31, Ser49, and Ser150), or four Leu codons (Leu32, Leu67, Leu144, and Leu170) were replaced. (B) NanoLuc translation results for each codon reassignment experiment. Translation reactions were performed in tfPURE supplemented with a 21-tRNA mixture (600 ng/µL), one tRNA variant (12 ng/µL each), and each NanoLuc template (1 nM) that contains 2 – 4 of a corresponding codon to be tested (21 + 1 NNN-Ala/Ser/Leu codons). Reactions were incubated at 30 °C for 16 h, after which NanoLuc activity was measured. As a control, translation reactions lacking the additional tRNA variant were conducted (21 code, gray bars) and compared to the data with the additional tRNA (21 +1 code, pink bars). Additional controls included translation without any tRNA (no tRNA) and translation using MGC with NanoLuc templates encoded by the original 21 codons (21 codons), both shown for comparison. Each dot represents three technical replicates, and error bars represent standard deviations. For each template, NanoLuc activity in the 21-code and corresponding 21+1-code conditions was compared using Welch’s t-test on luminescence. Statistical results are summarized in Table S9.

Translation reactions were performed in tfPURE supplemented with 21 tRNAs alone (21 code, equivalent to MGC) or 21 tRNAs plus one additional tRNA variant for Ala, Ser, and Leu (21 + 1 code) with the corresponding NanoLuc template (1 nM) at 30 °C for 16 h. NanoLuc activity was low when using only 21 tRNAs (Fig. 2B, gray bars, 21 code), but increased when the corresponding tRNA variant was supplied (pink bars, 21 + 1 code) for most of the tested codons. An exception was observed when Ser was assigned to the GGG codon, in which NanoLuc activity did not increase with 21 + 1 code for reasons that remain unclear. These results demonstrate that the nine tested vacant codons (i.e., excluding GGG) can be reassigned to the three amino acids, allowing the construction of up to 3^9^ = 19,683 types of nonSGCs, or approximately 20,000 codes. For GGG, we retained the original glycine assignment in all subsequent experiments. We further optimized the concentration of each additional tRNA variant, particularly for those that exhibited lower translation efficiency (Fig. S3). Based on these results, the concentrations of the following tRNAs were increased to the indicated concentrations in subsequent experiments: tRNA^Ala^_CAA_ (40 ng/µL), tRNA^Ala^_CGA_ (60 ng/µL), tRNA^Ala^_CGU_ (60 ng/µL), tRNA^Leu^_CGU_ (80 ng/µL), tRNA^Leu^_CGA_ (80 ng/µL), tRNA^Leu^_GCU_ (80 ng/µL).

### Design of non-standard genetic codes

We next designed nonSGCs. By assigning Ala, Ser, or Leu to the nine vacant codons (gray boxes in Fig. 3A), a total of 19,683 (= 3^9^) possible genetic codes can be generated. The objective of this study is to evaluate how the arrangement of the genetic code influences the impact of point mutations on protein function. To guide the selection of genetic codes to construct, we defined three types of mutational cost for each genetic code based on three physicochemical properties of amino acids, polar requirement (PR), molecular volume (MV), and hydropathy index (HI), according to previous studies (Freeland and Hurst, 1998; Haig and Hurst, 1991). PR reflects polarity-related interactions of amino acids and has been used as a classical measure of amino acid similarity in error minimization analyses. MV represents side-chain size and steric volume, which could affect protein packing and structural stability, whereas HI reflects hydrophobicity, which could be closely related to protein folding or hydrophobic core formation. The mutational costs, three values (Cost_PR_, Cost_MV_, and Cost_HI_) associated with each genetic code, represent the expected average negative effect of a single nucleotide substitution in a target gene on the protein function, estimated from the change in each physicochemical property of the substituted amino acid. To make the design of non-SGCs more explicit, we show one representative non-SGC together with the near-SGC in Fig. 3B. This comparison illustrates how assignment of Ala, Ser, or Leu to the vacant codon boxes changes the three mutational cost metrics, Cost_tPR_, Cost_MV_, and Cost_HI_.

**Figure 3.**
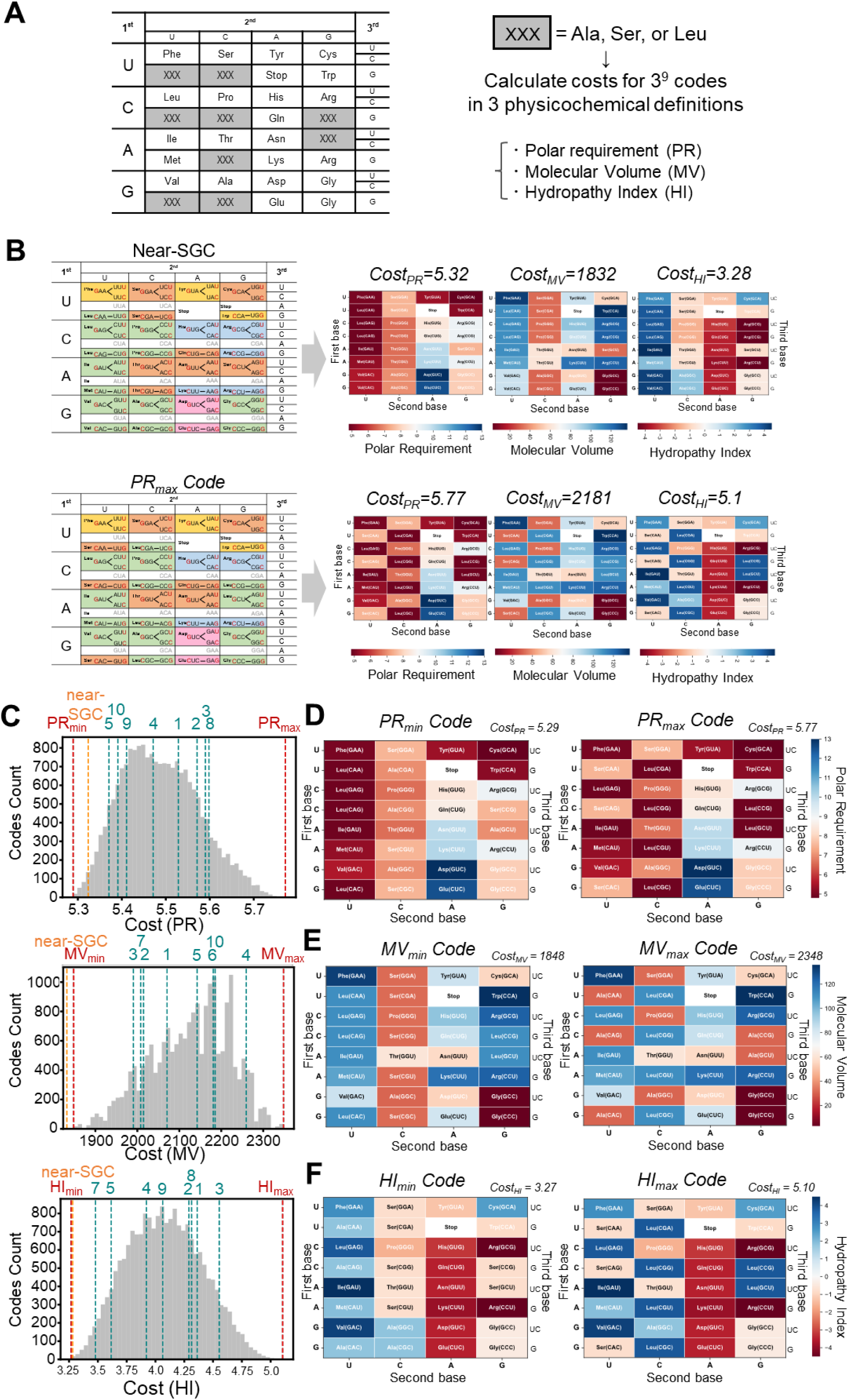
Distribution of mutational costs of reassigned genetic codes. (A) Calculation method of mutational costs for each genetic code based on three physicochemical properties of amino acids. The average change in each of the three physicochemical properties of amino acids upon single-nucleotide substitutions from the 21 codons was calculated (see Methods for details). In the reassigned genetic codes analyzed here, one of three amino acids (Ala, Ser, or Leu) was assigned to each of the nine vacant codons shown in gray, and the costs were calculated for all possible reassignment combinations. (B) Representative comparison of the near-SGC and PR_max_ code. Codon assignment schemes are shown on the left, and heatmap representations of the assigned amino acid values for polar requirement, molecular volume, and hydropathy index are shown on the right. The corresponding Cost_PR_, Cost_MV_, and Cost_HI_ values are indicated above each heatmap. (C) Distributions of mutational costs for each physicochemical property of amino acids. Dashed lines indicate the cost values of 10 genetic codes selected for experimental construction. Red dashed lines indicate the minimum and maximum cost values for each cost definition, and orange lines indicate the cost values of near-SGC. (D, E, F) Genetic codes exhibiting the minimum and maximum mutational costs based on PR (D), MV (E), and HI (F). The physicochemical values of amino acids assigned to each codon are shown as heatmaps.

The three types of mutation costs based on PR, MV, and HI were calculated for each of the 19,683 (= 3^9^) possible genetic codes as follows. First, we considered a virtual protein-encoding DNA sequence using the 21 codons shown in Fig. 1B (left) at equal frequencies. We then enumerated the set of codons reachable from each codon via single-nucleotide substitutions and calculated the average change in each physicochemical property of amino acids for each genetic code. In this calculation, mutation biases, including the relative frequencies of transitions and transversions, were incorporated as weighting factors. When mutations produced NNA codons, which are vacant in the genetic codes used here, their substitutions were neglected. The resulting average physicochemical change was defined as the mutational cost for the target genetic code. According to this calculation, a smaller mutational cost indicates that single-nucleotide substitutions lead to smaller changes in the corresponding amino acid properties and thus expectedly smaller changes in protein function, reflecting higher robustness against mutations.

The distributions of mutational costs calculated for each physicochemical property (Cost_PR_, Cost_MV_, and Cost_HI_) are shown in Fig. 3C. The near-SGC (orange dashed lines) was located in the low-cost region of the distributions for all three properties, consistent with previous theoretical studies (Freeland and Hurst, 1998; Haig and Hurst, 1991). We next selected genetic codes for construction and experimental evaluation. First, we selected the genetic codes exhibiting the maximum and minimum costs for each physicochemical property (Fig. 3B, red dashed lines; PR_min/max_, MV_min/max_, and HI_min/max_). These selected genetic codes are shown in Figs. 3D – 3 F, colored according to the physicochemical values of the assigned amino acids. The genetic codes of the minimum costs (PR_min_, MV_min_, and HI_min_) exhibit relatively ordered arrangements of the physicochemical values, whereas those with maximum costs (PR_max_, MV_max_, and HI_max_) exhibit relatively disordered arrangements. Because the PR_max_ and HI_max_ codes were identical, these five extreme genetic codes were selected for construction. Second, we selected five additional genetic codes representing various points across the cost distributions for all three properties (green dashed lines). These 10 nonSGCs (shown in Figs. S4–S6) were constructed by introducing the corresponding additional tRNA sets into MGC and used in subsequent experiments.

### Preparation and analysis of random mutation libraries of reporter genes

To compare functional robustness against mutations across different genetic codes, we prepared randomly mutagenized DNA libraries of three reporter genes, β-galactosidase (GAL), firefly luciferase (Luc), and mStayGold (Ando et al., 2023) (mSG), from the original sequences composed of only the 21 codons shown in Fig. 1B (left). DNA libraries were prepared through a three-step PCR procedure (Fig. S7A). In the first PCR, random mutations were introduced by error-prone PCR using Taq DNA polymerase in the presence of Mn^2+^. The mutation rate was tuned by varying the Mn^2+^ concentration across five levels: 10, 50, 100, 250, and 350 µM. For comparison, we also performed standard PCR using a high-fidelity polymerase (KOD Plus Neo) to prepare a low-mutation library. In the second PCR, a HiBiT tag was attached to the C-terminus using the high-fidelity polymerase for quantitation of translation levels. In the third PCR, Illumina sequencing adapters and unique barcode sequences were added using the high-fidelity polymerase. The products of the second PCR were used directly for translation experiments, whereas the products of the third PCR were subjected to Illumina sequencing to quantify mutation rates and characterize the resulting libraries.

Mutation frequencies at each position were calculated from the sequencing data obtained. The simple average of these values across all positions was defined as the mean per-base mutation rate. When using Taq DNA polymerase, the error rates increased monotonically with increasing Mn^2+^ concentration, ranging from 1.2 × 10^−3^ to 6.9 × 10^−3^ (Fig. S7B). In contrast, when using the high-fidelity polymerase, the error rates were much lower (ranging from 3.0 × 10^−4^ to 4.0 × 10^−4^). The mutations observed with the high-fidelity polymerase are thought to have been caused by sequencing errors because the rate is consistent with the reported sequencing error rate (Schirmer et al., 2016) and also the primer-binding regions, which did not undergo PCR amplification, exhibited a similar mutation rate (approximately 5.0 × 10^−4^), implying that the actual mutation rate obtained using the high-fidelity polymerase can be lower.

We further analyzed the patterns of nucleotide substitutions observed in the libraries. With Taq polymerase, transitions (A↔G or C↔T) accounted for 61 – 72% of all substitutions, and this bias was consistently observed across all Mn^2+^ concentrations tested (Fig. S7C). This pattern is consistent with previously reported mutation biases associated with this polymerase (Lin-Goerke et al., 1997). Visualization of the positional distribution of mutation rates further confirmed that error-prone PCR introduced mutations throughout the entire gene sequence (Fig. S7D).

### Translation of random mutation libraries using near-SGC

To examine the relationship between mutation rates and translated protein function, we performed translation reactions using the near-SGC (RV), containing higher concentrations of two tRNAs (tRNA^Val^_CAC_ and tRNA^Arg^_CCU_) as shown in Fig. 1D, and DNA libraries with different mutation rates (Fig. 4A). When using GAL library, we observed a decrease in mean protein activity as the mutation rate increased (Fig. 4B), indicating that a sufficient number of mutations were introduced to evaluate the mutational effect on protein function. Similar trends were observed for Luc and mSG (Figs. 4C and 4D), although the magnitude of the decrease varied among proteins. For example, protein activities decreased to 33%, 2.4%, and 79% for GAL, Luc, and mSG, respectively, at a mutation rate of approximately 0.002 per base, indicating that sensitivity to mutations differs substantially among proteins, consistent with a previous study (Lind et al., 2017).

**Figure 4.**
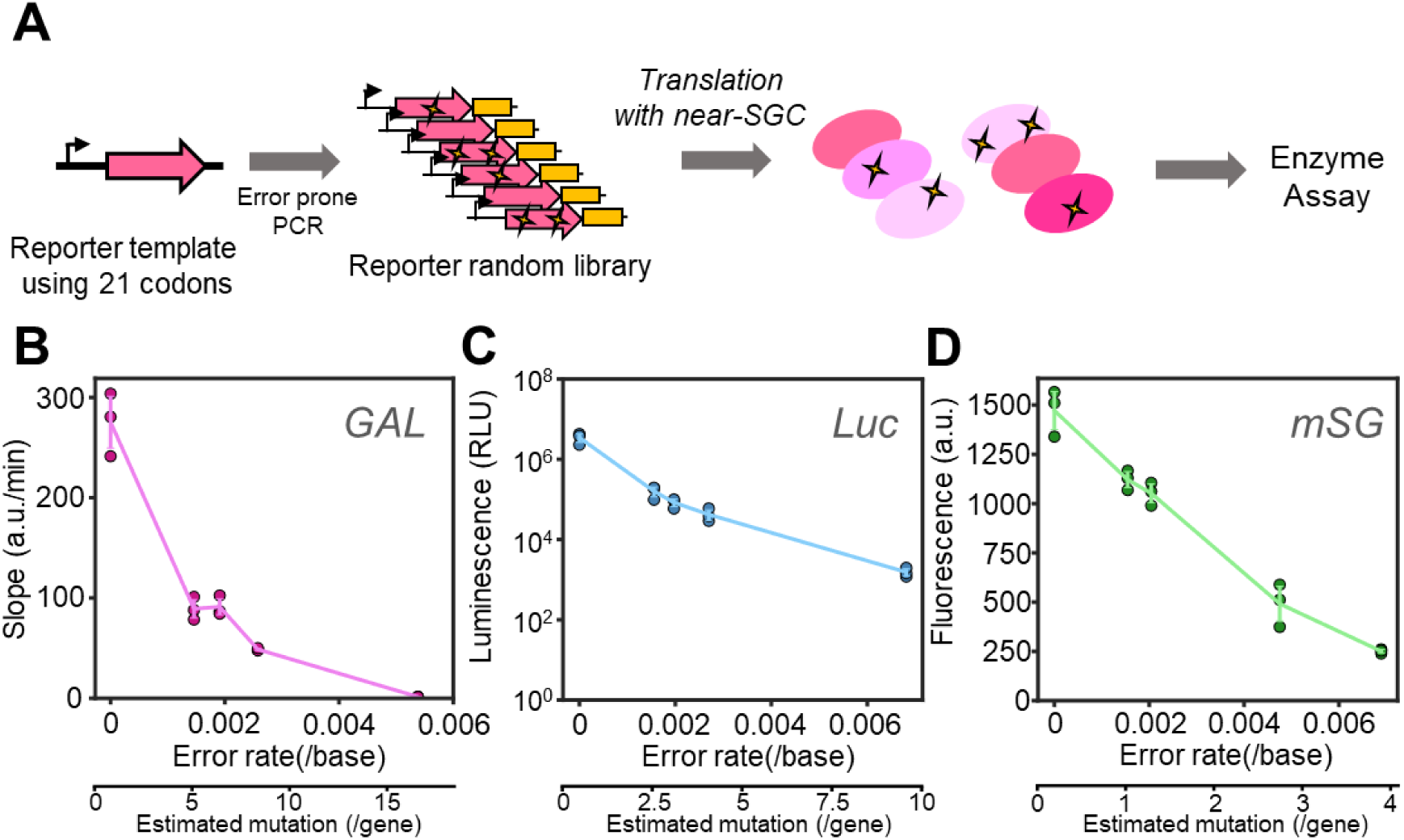
Translation of random mutagenesis libraries with near-SGC. (A) Schematic overview of the protein activity assay using a random mutation library. Reporter genes composed of the 21 codons were subjected to random mutagenesis by error-prone PCR at different Mn^2+^ concentrations to generate DNA libraries, as shown in Fig. S7. These libraries (5 nM) were translated using near-SGC, consisting of a 32-tRNA mixture (tRNA^IPEN^, tRNA^Val^_CAC_, and tRNA^Arg^_CCU_ at 100 ng/µL; all other tRNAs at 12 ng/µL) in tfPURE, including T7 RNA polymerase (1.7 U/µL) at 30 °C for 16 h, and each protein activity was measured. (B, C, D) Dependence of β-galactosidase (GAL) (B), firefly luciferase (Luc) (C), and mStayGold (mSG) (D) activity on mutation rate. Note that the vertical axis of panel C (Luc) is on a log scale. Each dot represents the results of three technical replicates, and error bars represent standard deviations. The lower x-axis indicates the estimated number of mutations per gene, calculated by multiplying the mutation rate per base by the coding sequence length of each reporter gene. Spearman’s rank correlation coefficients were ρ = −0.90 for GAL, ρ = −1.00 for Luc, and ρ = −1.00 for mSG.

### Translation of random mutation libraries using 10 non-SGCs

We next evaluated the effects of random mutations on protein activity using 10 non-SGCs constructed above. For this experiment, two random mutation libraries were used: a low-mutation library prepared using the high-fidelity polymerase and a high-mutation library prepared using Taq DNA polymerase at a Mn^2+^ concentration that yields mutation rates of 0.002 – 0.005 per base (0.0026 for GAL, 0.0027 for Luc, and 0.0048 for mSG, corresponding to approximately 8.0, 4.5, and 3.3 mutations per gene). We also plotted position-wise non-reference rates along the analyzed regions of each reporter gene, confirming that mutations were broadly distributed across the amplicons (Fig. S8). These libraries were transcribed and translated using 10 non-SGCs selected above, as well as the near-SGC (RV), and the resulting protein activities were measured (Fig. 5A).

**Figure 5.**
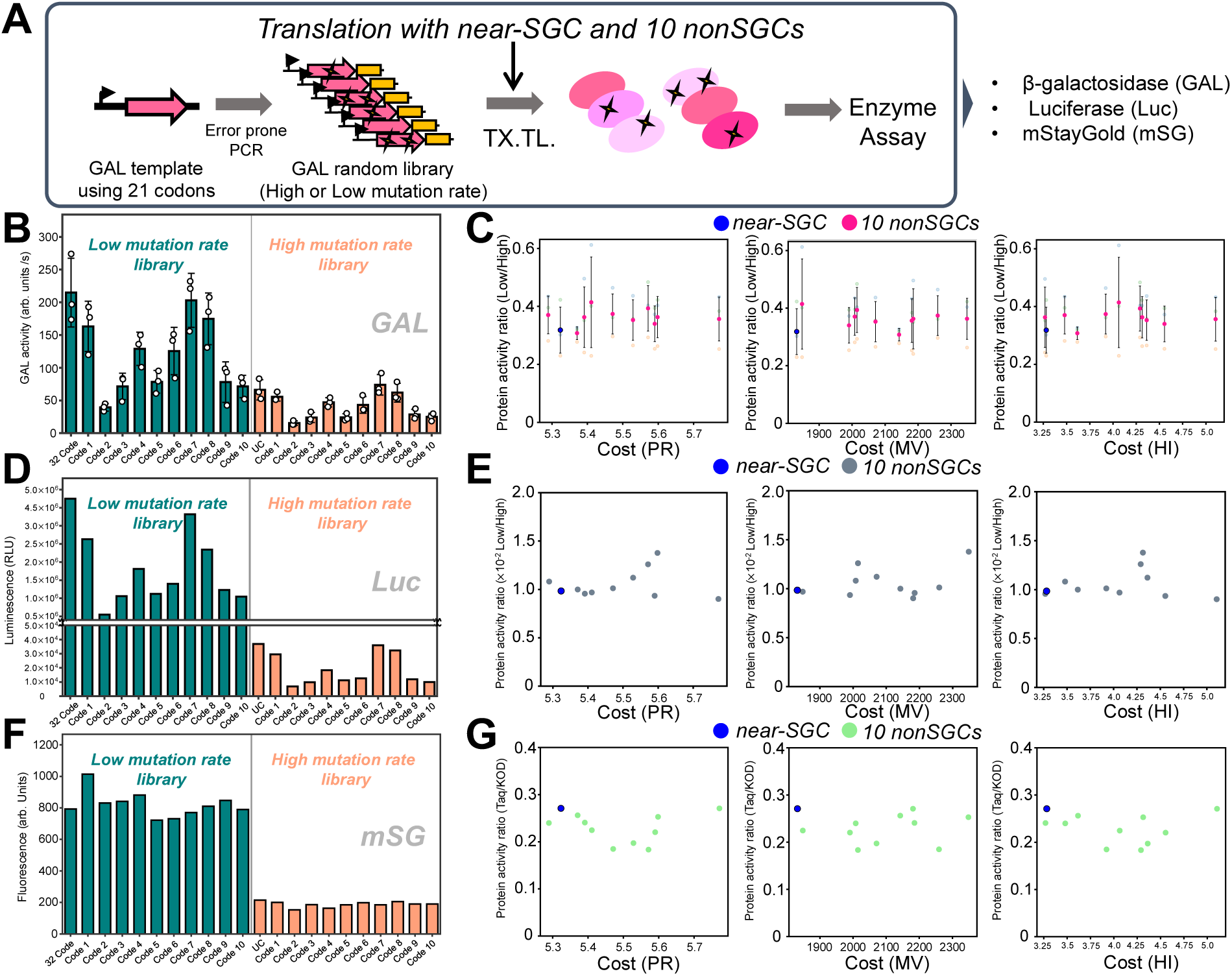
Translation of mutagenized DNA libraries with non-SGCs. **(A)** Schematic of the experiment for comparing protein activities translated with different genetic codes. Random libraries prepared at low and high mutation rates were translated using either the 10 non-SGCs or the near-SGC (RV). Translation conditions were identical to those described in Fig. 4. (B, D, F) Protein activities of products translated with each genetic code using low- and high-mutation DNA libraries. Activities are shown for β-galactosidase (GAL; B, mutation rate = 2.6 × 10^−3^ per base), firefly luciferase (Luc; D, mutation rate = 2.7 × 10^−3^ per base), and mStayGold (mSG; F, mutation rate = 4.8 × 10^−3^ per base). Quantification of protein synthesis levels for GAL is shown in Fig. S9. (C, E, G) Ratios of protein activity of high-mutation libraries to those of low-mutation libraries, plotted against the corresponding theoretical mutational costs. Data are shown for GAL (C), Luc (E), and mSG (G). Mean values of three technical replicates are shown with standard deviations for GAL. For GAL activity in (B), two-way ANOVA was performed using genetic code and mutation level as factors. Significant main effects of genetic code and mutation level were detected (both p < 0.0001), whereas their interaction was not significant. For (C), (E), and (G), Spearman’s rank correlation analysis was performed between each mutational cost metric and the high-/low-mutation activity ratio. Statistical details are summarized in Table S10.

We first examined GAL activities. For the low-mutation library, mean protein activity varied by up to 5.4-fold among different genetic codes (Fig. 5B left, green bars). For the high-mutation library, GAL activity decreased overall, while the relative differences in activity among genetic codes observed in the low-mutation library were broadly retained (Fig. 5B right, orange bars). Quantification of protein synthesis levels using the HiBiT assay revealed similar protein synthesis levels both among the different genetic codes and between the low- and high-mutation rates (Fig. S9), indicating that the variation in protein activity among genetic codes and between low- and high-mutation rates was not attributable to differences in translational levels. The variation in protein activity among genetic codes with the low-mutation library was possibly caused by differences in translational error frequencies among genetic codes. Such differences are not surprising because different tRNA sets were used for each genetic code, and each tRNA is expected to have distinct translational accuracies. However, the purpose of this study is to evaluate how the effect of mutations on protein function varies depending on the genetic code; this effect can be extracted even in the presence of different translational error rates by normalizing the protein activity of the mutated library by that of the less-mutated library.

To estimate the average activity reduction associated with increased mutational burden under each genetic code, we calculated the ratio of activity obtained from the high-mutation library to that from the corresponding low-mutation library and plotted this ratio against each of the three mutational costs (Fig. 5C). Data before this normalization are shown in Figs. S10–S12. According to the error minimization theory, genetic codes with higher mutational costs tend to reduce the protein activity more upon mutagenesis; thus, a negative correlation between mutational cost and protein activity would be expected. Contrary to this expectation, the effect of mutations on GAL activity remained nearly constant across the ranges of all three mutational costs examined (Fig. 5C).

We performed the same experiment for Luc (Fig. 5D) and mSG (Fig. 5F), and the results were similar to those obtained by GAL. For both proteins, activity with the low-mutation library varied among genetic codes, and decreased with the high-mutation library, although the variation among genetic codes was smaller for mSG. The activity ratios of the high- to low-mutation libraries showed no clear dependence on any of the three mutational costs (Figs. 5E and 5G). Taken together, these results indicate that the mutational robustness of individual reporter protein function did not substantially differ among the genetic codes that exhibit the mutational cost ranging from 5.29 to 5.77 for PR, 1848 to 2348 for MV, and 3.27 to 5.10 for HI.

## Discussion

In this study, we constructed 10 non-SGCs by assigning Ala, Ser, and Leu to vacant codons in MGC. These non-SGCs represent mutational costs ranging from 5.29 to 5.77 for Cost_PR_, 1848 to 2348 for Cost_MV_, and 3.27 to 5.10 for Cost_HI_, most of which are higher than those of SGC. The error minimization theory predicts that the deleterious effect of mutations on protein function becomes more severe when using genetic codes with higher mutational cost (Freeland et al., 2003; Freeland and Hurst, 1998; Haig and Hurst, 1991; Massey, 2008). In contrast to this prediction, translation experiments using the non-SGCs and mutagenized libraries revealed that the effects of mutations on protein functions were similar across all genetic codes and reporter genes examined here, suggesting that the mutational robustness of protein activity remained largely unchanged within at least the ranges of mutational cost tested in this study. It should be noted that this conclusion is limited to the activity of individual reporter proteins translated in a reconstituted in vitro system. Therefore, whether similar trends would be observed at the level of cellular fitness or long-term evolution remains an open question. Moreover, our results do not exclude other possible roles of SGC organization. The SGC may have been shaped by multiple factors, including robustness to translational errors, historical constraints during genetic code expansion, biosynthetic or coevolutionary relationships among amino acids, stereochemical interactions, and effects on protein evolvability (Katoh and Suga, 2023; Koonin and Novozhilov, 2017, 2009; Novozhilov et al., 2007; Wong, 2005).

Although protein amounts quantified by the HiBiT tag were comparable among genetic codes, GAL activities differed substantially. This indicates that the activity differences among genetic codes were not primarily attributable to differences in the amount of C-terminally completed translation products. The HiBiT assay does not provide information on the fraction of catalytically active protein, including sequence fidelity or folding state, and therefore cannot distinguish among these possibilities. Detailed characterization of translated products by mass spectrometry would provide further mechanistic insight into how individual non-SGCs affect protein quality. However, the primary objective of the present study was to compare mutation-dependent activity loss across genetic codes. Therefore, we evaluated this effect by normalizing the activity of the high-mutation library to that of the corresponding low-mutation library within each genetic code.

A further limitation of this study is that the reporter activities were measured at the level of pooled random mutation libraries. Therefore, the high-/low-mutation activity ratio used in this study should be interpreted as the relative reduction in average activity caused by increasing the mutational burden in a heterogeneous mutation pool, rather than as the effect of identical variants before and after additional mutations. This library-averaged approach was chosen because the mutational costs considered here are also defined as average expected physicochemical effects over many possible single-nucleotide substitutions. In addition, because the non-SGCs constructed in this study were generated by reassigning only Ala, Ser, and Leu, the detectable effects may depend on how frequently mutations involving these amino acids occur in each reporter gene and whether the affected positions are functionally important. If genetic code-dependent effects are restricted to a small subset of deleterious variants, such effects may be masked in pooled activity measurements. Future studies using defined variants or high-throughput genotype–phenotype mapping assays will be required to determine the mutation-specific and position-specific mechanisms underlying genetic code-dependent effects on protein function (Rozhoňová et al., 2024).

It should be noted that the range of mutational costs experimentally tested in this study is limited because we could only reassign three amino acids (Ala, Ser, and Leu) to the vacant codons in MGC. If all 20 amino acids could be reassigned to the nine vacant codons, the possible range of mutational costs would be 5.29 – 8.49 for Cost_PR_, 1826 – 2805 for Cost_MV_, and 3.21 – 6.65 for Cost_HI_ (Fig. S13). The genetic codes constructed in this study cover approximately 15.1%, 51.1%, and 53.3% of these possible cost ranges, for PR, MV, and HI, respectively. Furthermore, if we could assign any amino acids to any codon boxes with a degenerate pattern similar to that of SGC, as assumed in previous theoretical studies (Freeland and Hurst, 1998; Haig and Hurst, 1991), the cost ranges were 2.67 – 4.79 for Cost_PR_, 1952 – 3209 for Cost_MV_, and 4.69 – 12.44 for Cost_HI_ for one million randomly sampled genetic codes. The non-SGCs constructed in this study covered 18.4%, 37.6%, and 50.8% of these possible cost ranges for PR, MV, and HI, respectively (Fig. S14). We consider that the cost ranges examined in this study represent substantial fractions, especially for MV and HI. Although the near-SGC did not necessarily exhibit the lowest cost for each individual physicochemical metric, this does not mean that it is unfavorable in the multidimensional cost space. Because the SGC may reflect a balance among multiple physicochemical constraints rather than optimization of a single property, we also calculated integrated cost indices by combining Cost_PR_, Cost_MV_, and Cost_HI_ after min–max normalization or z-score normalization. In both integrated indices, the near-SGC showed the lowest overall cost when compared with all 19,683 candidate non-SGCs (Fig. S15), indicating that no candidate non-SGC exhibited a lower combined cost than the near-SGC when the three physicochemical properties were considered jointly. However, we also think that the finding of this study — mutational robustness does not detectably depend on the genetic codes — might change if nonSGCs with broader cost ranges could be constructed.

This limitation in the achievable mutational cost ranges in this study arises from the fact that only three amino acids (Ala, Ser, and Leu) can be reassigned to different codons. The other amino acids cannot be reassigned to different codons because their corresponding tRNAs require the cognate anticodon sequence for recognition by the corresponding aaRS. To construct genetic codes with a broader range of mutational costs, it will be necessary to identify new aaRSs that do not recognize anticodon sequence but still catalyze aminoacylation for the other 17 amino acids. To date, such aaRSs have not been clearly reported, but some aaRSs, such as a mitochondrial aaRS with altered anticodon recognition (Su et al., 2011) and aaRS mutants that shuffle the anticodon recognition domain (Brevet et al., 2003), may be suitable for this purpose. Although numerous engineered orthogonal aaRS–tRNA pairs have been developed to incorporate non-canonical amino acids into proteins, these systems are typically designed to target stop or limited sense codons(Costello et al., 2024; Robertson et al., 2021; Shandell et al., 2021; Zeng et al., 2014). Alternatively, ribozyme-based aminoacylation systems such as flexizymes (Goto and Suga, 2021; Jones et al., 2025; Murakami et al., 2006; Terasaka et al., 2014) offer a more flexible route to assigning diverse amino acids to vacant codons, although protein synthesis using ribozyme-based aminoacylation is still a challenge (Chen et al., 2021).

The results of this study suggest that altering the arrangement of the genetic code does not necessarily lead to a drastic loss of functional robustness against mutations. This finding indicates that genetic codes can be redesigned for specific purposes without severe functional penalties, at least in vitro. This insight paves the way for utilizing engineered genetic codes (Huang et al., 2024; Pines et al., 2017; Zürcher et al., 2022) for broader applications in protein engineering and in vitro synthetic biology, such as genetic code expansion using non-canonical amino acids (Iwane et al., 2016; Jones et al., 2025; Katoh and Suga, 2022; McFeely et al., 2023; Passioura et al., 2018), and the design of artificial translation systems orthogonal to natural translation systems for biocontainment (Calles et al., 2019; Fujino et al., 2024, 2020; Nyerges et al., 2023; Robertson et al., 2021). Accelerated protein evolution may also be achievable using non-standard genetic codes (Pines et al., 2017; Zürcher et al., 2022) because the structure of the genetic code determines the amino acids accessible through mutation. While some theoretical studies suggest that alternative genetic codes could enhance evolvability (Pines et al., 2017), others have argued that the standard genetic code itself possesses advantageous evolvability features (Firnberg and Ostermeier, 2013; Rozhoňová et al., 2024). The knowledge obtained in this study enhances the usefulness of genetic code engineering in vitro for broader applications.

## Method

### DNA preparation

All DNA templates encoding reporter proteins were obtained through a fragment synthesis service provided by Twist Bioscience. The NanoLuc reporter genes used for translation under MGC, near-SGC, or SGC were designed to utilize 21, 32, or 46 codons (Fig. 1B). For validation of translation by each anticodon variant tRNA in Fig. 2, NanoLuc reporter genes were designed based on the 21 codon set, in which two alanine, three serine, or four leucine codons were replaced by target codons, producing a total of 25 distinct NanoLuc templates. For assays involving non-SGCs, DNA sequences encoding β-galactosidase (GAL), firefly luciferase (Luc), and mStayGold (mSG) were designed using the 21 codon set. The sequences of all DNA templates used in this study are provided in Supplementary Table S1.

To prepare templates for translation reactions, PCR amplification was performed using primers 1 and 2, which were common to all constructs. For PCR under low-mutation conditions, KOD Plus Neo DNA polymerase (Toyobo) was used. Error-prone PCR conditions are described in a separate section below. To prepare templates with a C-terminal HiBiT tag for protein quantification (Fig. S7, 2^nd^ PCR), a HiBiT tag fragment was first amplified using primers 3 and 4. This fragment was then fused to the coding region of each reporter gene by overlap extension PCR; PCR products encoding GAL, Luc, or mSG were amplified using primers 1 and 5 (GAL), 6 (Luc), or 7 (mSG), respectively, using either KOD Plus Neo for the low-mutation library or Taq DNA polymerase for the higher-mutation libraries, and subsequently fused with the HiBiT fragment using primers 1 and 8. The resulting DNA fragments were purified using the FastGene Gel/PCR Extraction Kit (Nippon Genetics), and DNA concentrations were determined from A260 measurements using a NanoDrop spectrophotometer (Thermo Fisher Scientific). The sequences of all primers used in this study are listed in Supplementary Table S2.

### Error-prone PCR

To introduce random mutations into protein-coding DNA, error-prone PCR was performed as described schematically in Fig. S7 (1^st^ PCR). PCR reactions were carried out using Taq DNA polymerase (New England Biolabs) in the manufacturer-recommended Standard Taq Reaction Buffer supplemented with additional MgCl_2_ and MnCl_2_. The final concentration of MgCl_2_ was adjusted to 3.0 mM. To modulate the mutation rates, MnCl_2_ was added at final concentrations of 10, 50, 100, 250, or 350 µM. PCR amplification was performed for 30 cycles, and the resulting products were used as random DNA libraries for subsequent experiments.

### tRNA preparation

Among the 21 tRNAs used in MGC, those corresponding to our previous work (Miyachi et al., 2025, 2022) were used without modification, whereas the remaining 11 tRNAs for near-SGC and 14 for SGC were newly prepared by replacing their anticodon sequences. For these tRNAs, plasmid templates encoding the original sequences of the corresponding 21-tRNAs set were used as templates, and only the anticodon region was replaced by site-directed mutagenesis using inverse PCR. Anticodon variants of tRNA^Ala^, tRNA^Ser^, and tRNA^Leu^ were prepared in the same manner, with substitutions restricted to the anticodon region. The primer sets used for inverse PCR and the sequences of all tRNAs are listed in Tables S2 and S3.

Plasmids constructed by inverse PCR were used as templates for tRNA transcription. To prepare transcription templates, PCR amplification was performed using a forward primer containing a T7 promoter sequence and a reverse primer designed for run-off transcription carrying a 2’-O-methyl modification (Tables S2 and S3). PCR reactions were carried out using KOD Plus Neo DNA polymerase (Toyobo) according to the manufacturer’s instructions. Following amplification, PCR products were treated with DpnI at 37 °C for 2 h to remove template plasmids. The resulting DNA fragments were purified, and DNA concentrations were determined as described above.

Of the 46 tRNAs used in this study, the majority were prepared by in vitro transcription using T7 RNA polymerase. To enable efficient transcription, tRNAs whose native sequences do not begin with guanosine at the 5’-end were redesigned as 5’-G variants unless otherwise noted. Consequently, most tRNAs were synthesized by in vitro transcription regardless of their native 5’-terminal nucleotide. Exceptions were tRNA^Asn^_GUU_, tRNA^Gln^, tRNA^fMet^, tRNA^Ile^_GAU_, tRNA^Trp^, and tRNA^Pro^_GGG_, which were obtained by chemical synthesis (Agilent), as our previous report (Miyachi et al., 2022).

*In vitro* transcription reactions were performed in a mixture containing 40 mM Tris–HCl (pH 8.0), 10 mM MgCl_2_, 2 mM spermidine, 5 mM DTT, 1 U/µL T7 RNA polymerase (Takara Bio), 2 mM each NTP (ATP, GTP, CTP, and UTP), 3 mM GMP, 5 ng/µL template DNA, 2 U/µL inorganic pyrophosphatase (New England Biolabs), and 0.8 U/µL RNasin (Promega). Reactions were incubated at 37 °C for 12 h, and RNA products were purified using the PureLink RNA Mini Kit (Invitrogen). Purified tRNAs were dissolved in water and stored at −80 °C until use. RNA concentrations were determined by absorbance at 260 nm.

### Calculation of mutational cost

The mutational costs for each genetic code, *Cost*(*C*), were calculated based on previously proposed formulations of genetic code optimality(Freeland and Hurst, 1998; Haig and Hurst, 1991). In this calculation, the cost reflects the expected change in amino acid property caused by single-nucleotide substitutions from the initial 21-codon sets in this study (Fig. 1B). Details of calculations are as follows. We denote each codon in the 21-codon set as *β*, and the amino acid encoded by *β* as *AA*(*β*). For each *β*, we identified the set of codons reachable by a single-nucleotide substitution, denoted as *N*(*β*). Each element of this set is represented as a pair (*β*′, *w*_*β*→*β*_*_′_*), where *β′* is a neighboring codon box and *w*_*β*→*β*_*_′_* is a weight reflecting the relative likelihood of the corresponding nucleotide substitution. These weights account for differences in substitution probabilities depending on nucleotide position and whether the substitution is a transition or transversion. The substitution weights were adopted from the previous reports(Freeland and Hurst, 1998; Omachi et al., 2023) and are summarized in Table S4.

For each possible substitution from *β* to *β′*, we calculated the physicochemical distance between the corresponding amino acids, Δ(*AA*(*β*), *AA*(*β′*)). This distance was evaluated using three physicochemical properties: polar requirement (PR), molecular volume (MV), and hydropathy index (HI). For each property, the distance was defined as the squared difference between the property values of the two amino acids. For example, for PR, the distance function is defined as:

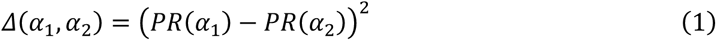

The expected cost contribution of each substitution was calculated by multiplying the amino acid distance by the corresponding substitution weight. The overall mutational cost was then obtained by summing these weighted distances over all single-nucleotide substitutions from all codon boxes in the 21-codon set and dividing by the total number of substitutions considered. Formally, the cost of the genetic code *C* was defined as:

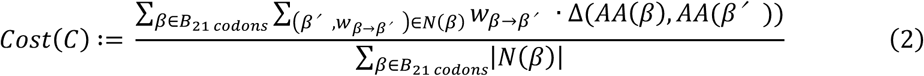

Here, *B*_21 *codons*_ denotes the set of codon boxes fixed in the 21-box code. The denominator represents the total number of possible single-nucleotide substitutions considered, ensuring that the cost corresponds to the average expected physicochemical impact per mutation. Physicochemical property values for each amino acid were taken from the datasets adopted by Haig and Hurst (Haig and Hurst, 1991), and are listed in Table S5.

For integrated cost analysis, Cost_PR_, Cost_MV_, and Cost_HI_ were combined after normalization because the three metrics have different numerical scales. For min–max normalization, each metric *m* ∈ {*PR*, *MV*, *HI*} cost *Cost*_*mnorm*_(*C*) was normalized across the set *G* of candidate non-SGCs as

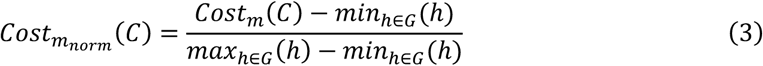

The integrated min–max cost was defined as the equal-weight mean of the three normalized costs. For z-score normalization, each cost was transformed as

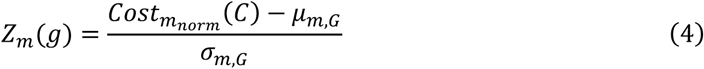

where *μ*_*m*,*G*_ and *σ*_*m*,*G*_are the mean and standard deviation across candidate non-SGCs. The integrated z-score cost was defined as the equal-weight mean of the three z-scores. The near-SGC was evaluated using the same normalization parameters calculated from the candidate non-SGC distribution.

### Sequence analysis of random libraries

Random mutagenesis libraries (2^nd^ PCR products; Fig. S7) were used as templates for amplicon preparation for sequencing (3^rd^ PCR). Using 40 pM of each library as input, an approximately 500-bp region spanning the coding sequence of each reporter gene was amplified by PCR using KOD Plus Neo DNA polymerase (Toyobo). Primers used for this PCR contained both sample-identifying barcodes and Illumina adapter sequences. For GAL, primers 107–111 were used as forward primers with primer 112 as a common reverse primer. For Luc, primers 113–117 were used as forward primers with primer 118 as a common reverse primer. For mSG, primers 119–123 were used as forward primers with primer 124 as a common reverse primer. PCR products were purified using the FastGene Gel/PCR Extraction Kit (Nippon Genetics). DNA concentrations were determined by absorbance at 260 nm and subjected to Illumina MiSeq sequencing.

Paired-end FASTQ files generated by the Illumina platform were used for all analyses. Libraries were prepared from three reporter genes across five error-prone PCR conditions, with each sample corresponding to an amplicon of approximately 500 bp. Sample-identifying barcode sequences were positioned at the 5′ end of read 1 (R1) of the pair-end sequencing, and the R1 reads covering approximately 300 bp from the 5′ side of each amplicon were used for non-reference-rate analysis. Demultiplexing was performed using cutadapt, requiring a perfect full-length match to the barcode sequence; only read pairs in which the R1 barcode matched exactly were retained, and all other read pairs were excluded from downstream analyses. Demultiplexed FASTQ files were generated for each sample. Paired-end reads were aligned to the corresponding amplicon reference sequence (generated for each reporter gene and condition) using BWA-MEM. Alignments were processed with SAMtools to generate sorted and indexed BAM files. Position-wise base information was extracted using SAMtools mpileup, applying filters of mapping quality MAPQ ≥ 20 (−q 20) and base quality Phred Q ≥ 30 (−Q 30). Insertions and deletions were excluded from all analyses.

For each nucleotide position *i*, the mutation count *n*_*i*_ was defined as the total number of bases in the pileup that differed from the reference base, and the total coverage at that position was denoted as *d*_*i*_. The position-wise error rate was calculated as:

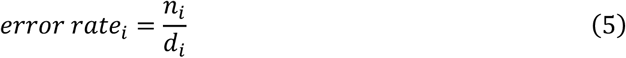

The representative error rate for a given sample (i.e., a specific PCR condition) was defined as the simple average across all positions in the analyzed region (length *L*):

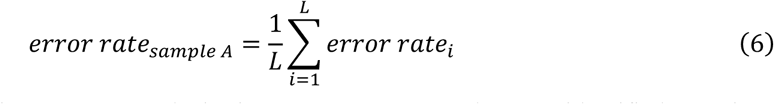

To characterize the mutation spectrum, substitution types were counted among identified mutations by tallying the frequency of each reference-to-alternate base change (e.g., A>C). In addition, for each reporter gene, primer-binding regions (19 bp) were defined based on the known primer sequences. Mutation rates and substitution spectra were calculated separately for primer-binding regions and non-primer regions. Primary comparisons were performed using profiles from non-primer regions, whereas primer-region profiles were used to assess the background mutation rate under low-mutation conditions (using KOD Plus Neo).

### tRNA-free PURE system (tfPURE) preparation

All components of the laboratory-made PURE system were prepared following previously established procedures. Briefly, individual factors were expressed with histidine tags and purified by affinity chromatography, followed by gel-filtration chromatography, as described previously (Miyachi et al., 2025). Additionally, EF-Tu was subjected to two successive rounds of affinity purification in a stringent buffer to eliminate contaminating tRNAs, as described previously (Miyachi et al., 2025, 2022). Ribosomes were purified by a butyl column and sucrose cushion method and further processed by ultrafiltration to remove residual tRNAs, using the same strategy as in our previous work (Miyachi et al., 2025, 2022). The complete composition and concentrations of tfPURE used in this study are provided in Table S6.

### Cell-free translation with reconstituted tRNAs

Cell-free translation reactions were performed using tfPURE supplemented with defined mixtures of in vitro-synthesized tRNAs and reporter gene DNA templates. The concentrations of DNA templates are specified in the corresponding figure legends, and the concentrations of tRNAs used for each genetic code are summarized in Table S7. Reaction mixtures were incubated at 30°C for 16 h.

Following incubation (or during incubation for mSG), reporter protein production was quantified by three different methods. For firefly luciferase (Luc) or NanoLuc, a 1 µL aliquot of the translation reaction was mixed with 30 µL of Luciferase Assay Reagent (Promega) or 50 µL of NanoLuc Assay Reagent (Promega), respectively, and luminescence was measured using a GloMax luminometer (Promega). For mSG, fluorescence signals were monitored continuously for up to 16 h using an Mx3005P real-time PCR system (Agilent Technologies) during the translation reaction. The fluorescence intensity at 0 h was subtracted from that at 16 h, and the resulting difference was used as a measure of mSG activity. For GAL (Matsuura et al., 2011), a 1 µL aliquot of the translation reaction was added to 9 µL of GAL reaction buffer (final concentrations: 50 mM HEPES–KOH, pH 7.4; 1 mM MgCl_2_; 5 mM DTT; 5 µM TokyoGreen-βGal (Sekisui Medical, Japan)). Fluorescence was recorded at 1-min intervals for 60 min using an Mx3005P instrument with FAM detection settings. GAL activity was determined from the slope of the linear region of the fluorescence time course.

## Supporting Information

Supporting Information includes supplemental figures (Figures S1 – S12) and tables (Tables S1 – S7).

## Author contribution

RM and NI planned the experiments and wrote the manuscript. RM performed the experiments.

## Acknowledgement

We thank Ms. Kayo Aoyama and Ayu Saito for their technical support.

