## Supplemental figures and tables for "Experimental verification of the error minimization theory using non-standard genetic codes constructed in vitro"

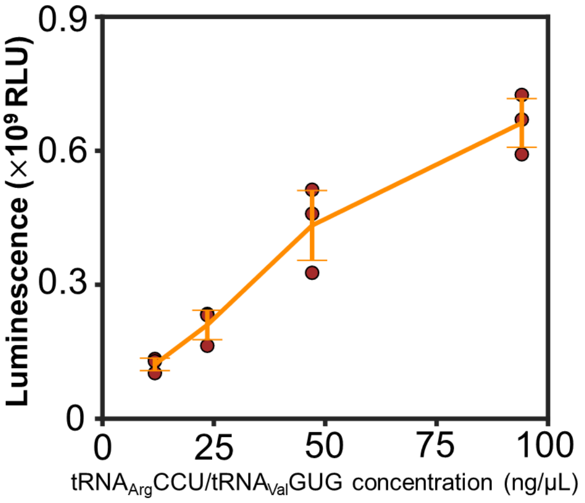


**Figure S1. Effect of tRNA$^{Val}_{CAC}$ and tria$^{rch}$ concentrations on NanoLuc translation using near-SGC.**

The concentrations of both tRNAs were increased simultaneously at equal ratios. Each dot represents the results of three technical replicates, and error bars represent standard deviations.

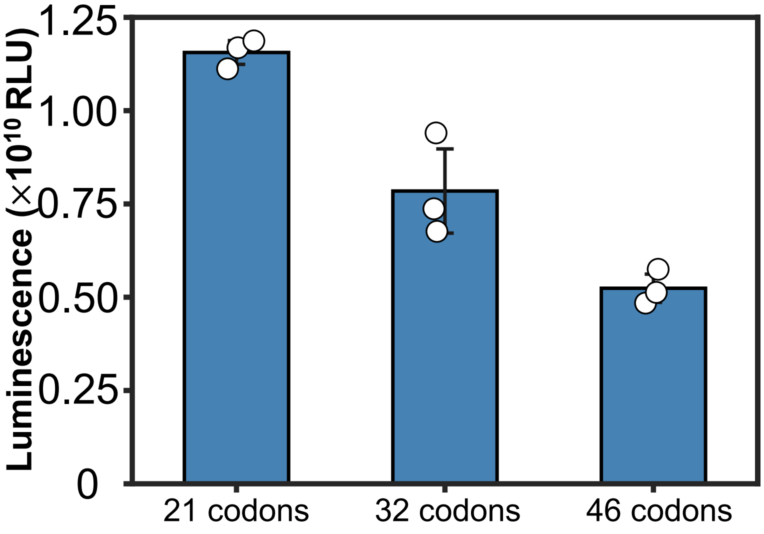


**Figure S2. Translation of each NanoLuc template using Native *E. coli* tRNAs.**

Translation of NanoLuc templates encoded with 21, 32, or 46 codons using native *E. coli* tRNAs (600 ng/µL) in the tfPURE system.

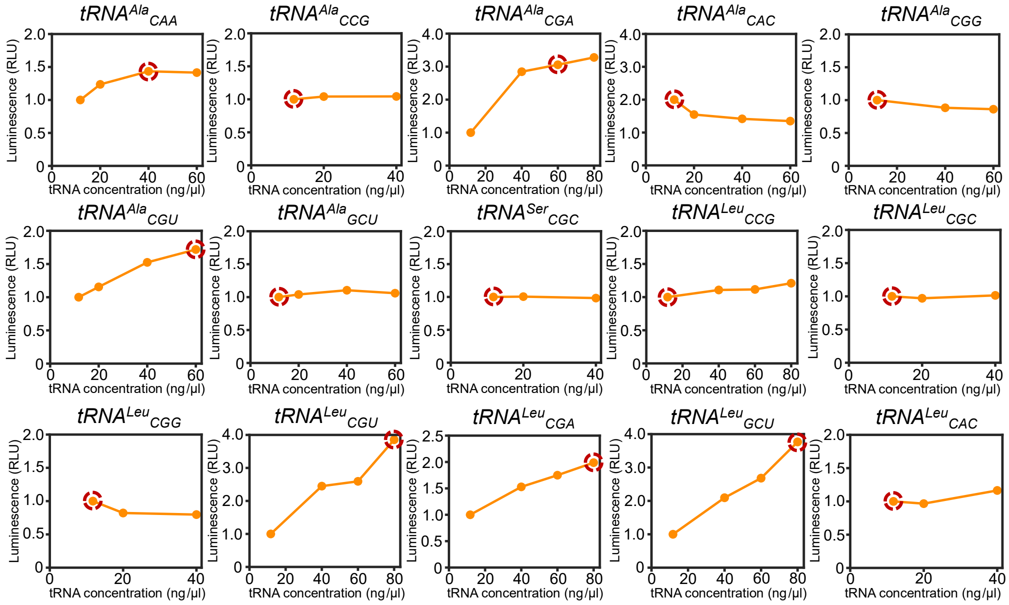


**Figure S3. Optimization of the concentration of each anticodon variant tRNA**

For 15 of the 25 anticodon variant tRNAs that exhibited relatively low translational activity, the effect of increasing tRNA concentration on translation efficiency was examined. Translation assays were performed under the same conditions as described in Fig. 2. The original concentration of each tRNA variant was 12 ng/µL. The concentrations selected for use in subsequent experiments are indicated by red circles.

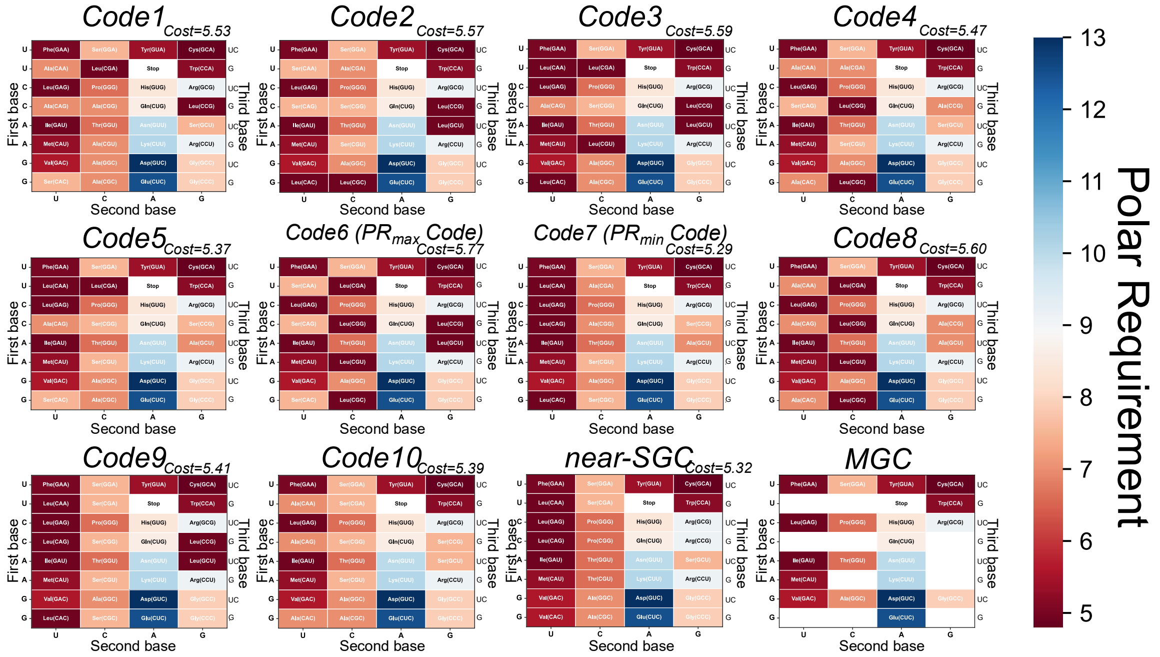


**Figure S4. Polar requirement values of amino acids assigned in the constructed non-SGCs**

The experimentally constructed non-SGCs and their corresponding mutational costs are shown. For each genetic code, the polar requirement values of amino acids assigned to individual codons are displayed as heatmaps. Near-SGC and MGC are shown for comparison.

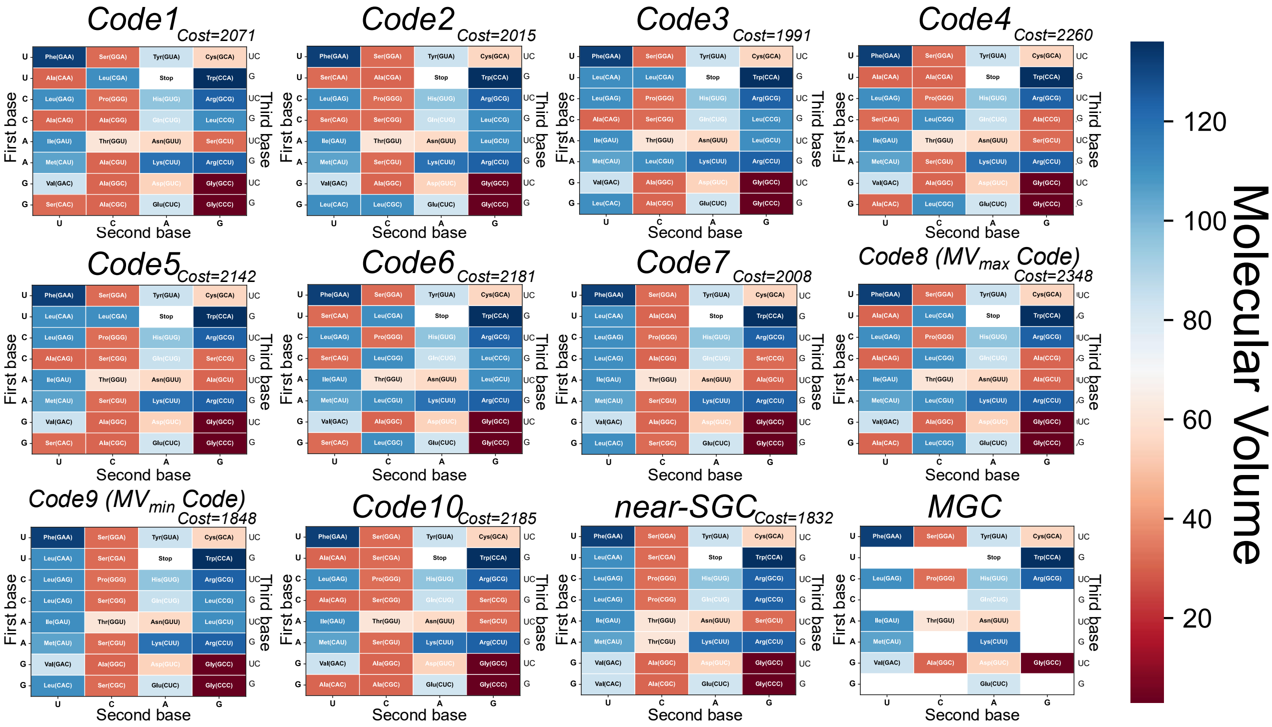

**Figure S5. Molecular volume values of amino acids assigned in the constructed non-SGCs**

The experimentally constructed non-SGCs and their corresponding mutational costs are shown. For each genetic code, the molecular volume values of amino acids assigned to individual codons are displayed as heatmaps. Near-SGC and MGC are shown for comparison.

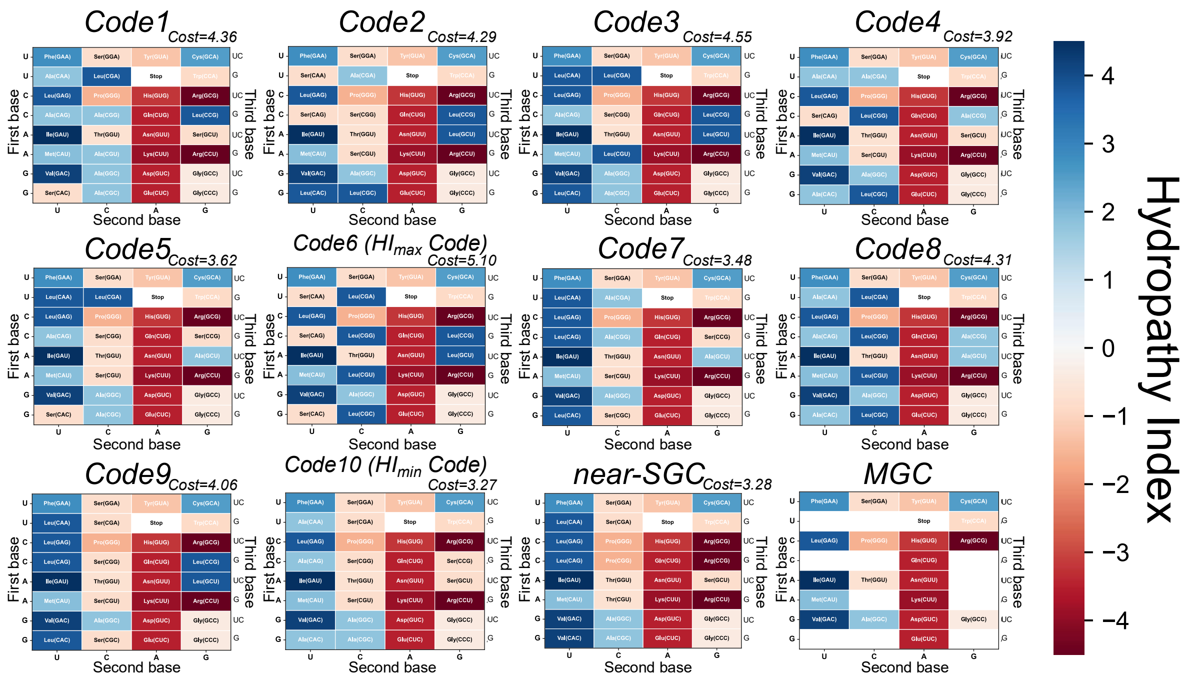

**Figure S6. Hydropathy index values of amino acids assigned in the constructed non-SGCs**

The experimentally constructed non-SGCs and their corresponding mutational costs are shown. For each genetic code, the hydropathy index values of amino acids assigned to individual codons are displayed as heatmaps. Near-SGC and MGC are shown for comparison.

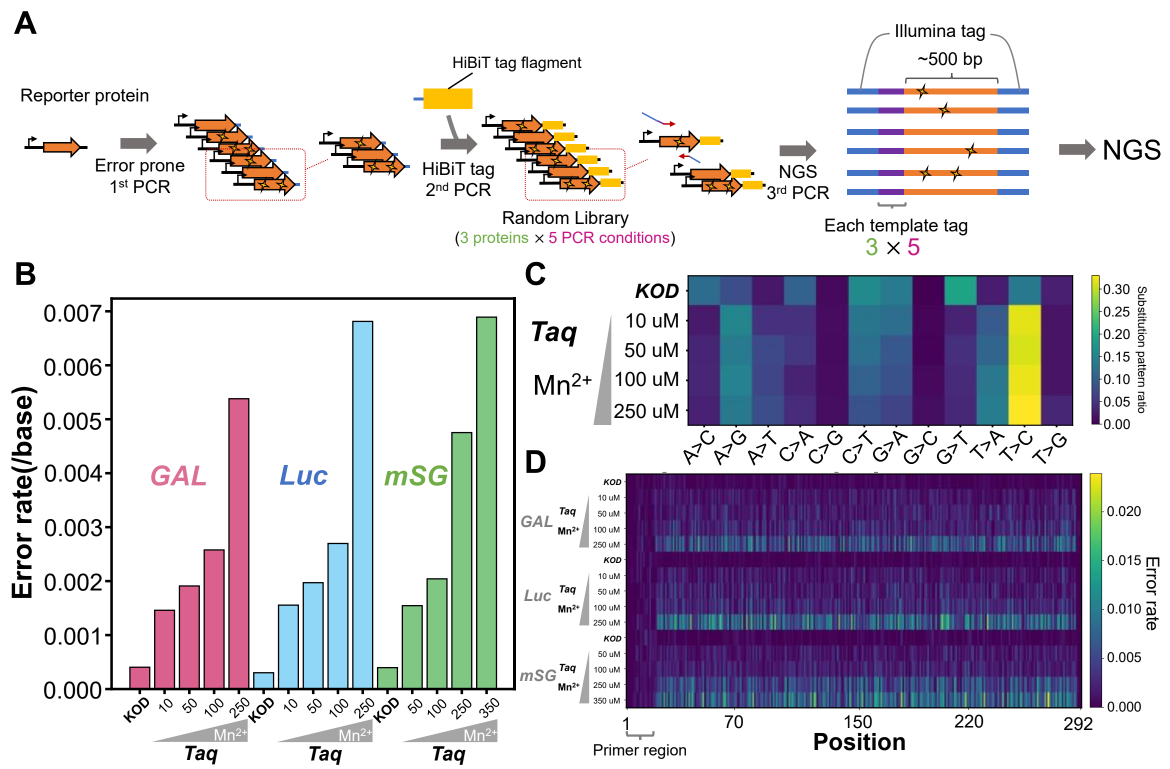


**Figure S7. Construction of random libraries by error-prone PCR and analysis of mutation patterns**

(A) Schematic of the method for preparing DNA libraries. Reporter gene templates composed of the 21 codons (Fig. 1B) were amplified by a first PCR using either a high-fidelity polymerase (KOD Plus Neo) or Taq DNA polymerase under error-prone conditions in the presence of $Mn^{2+}$. A second PCR was then performed using the high-fidelity polymerase to append a C-terminal HiBiT tag for protein quantification; these products were used for subsequent translation assays under each genetic code. For sequencing, a third PCR was performed using the high-fidelity polymerase to add Illumina adapters and unique barcode sequences identifying each reporter gene and PCR condition. (B) Mean per-base error rates under each 1$^{st}$ PCR condition. The mutation probability was calculated for each position of the template, and the simple average across all positions was used as the mean per-base error rate (see Methods). (C) Mutation spectra under each 1$^{st}$ PCR condition. The fractions of each substitution type were computed across all observed substitutions (e.g., “A>C” denotes substitution from A to C). (D) Position-wise distribution of error rate. The mutation probability at each nucleotide position is shown.

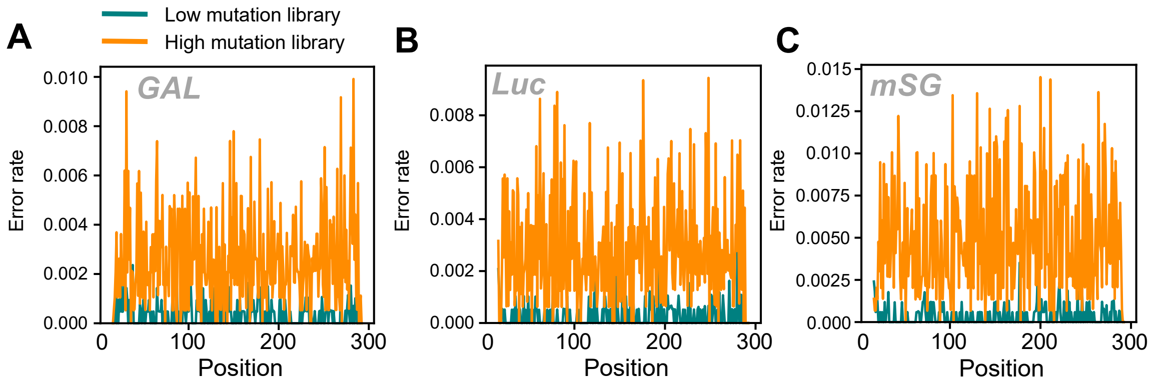


**Figure S8. Position-wise distribution of non-reference rates in random mutagenesis libraries.**

For each reporter gene, the non-reference rate at each position was calculated from amplicon sequencing data and plotted along the analyzed region (approximately 300 nt in the middle of each gene). Low- and high-mutation libraries are shown separately. These profiles show the positional distribution of mutations introduced by error-prone PCR.

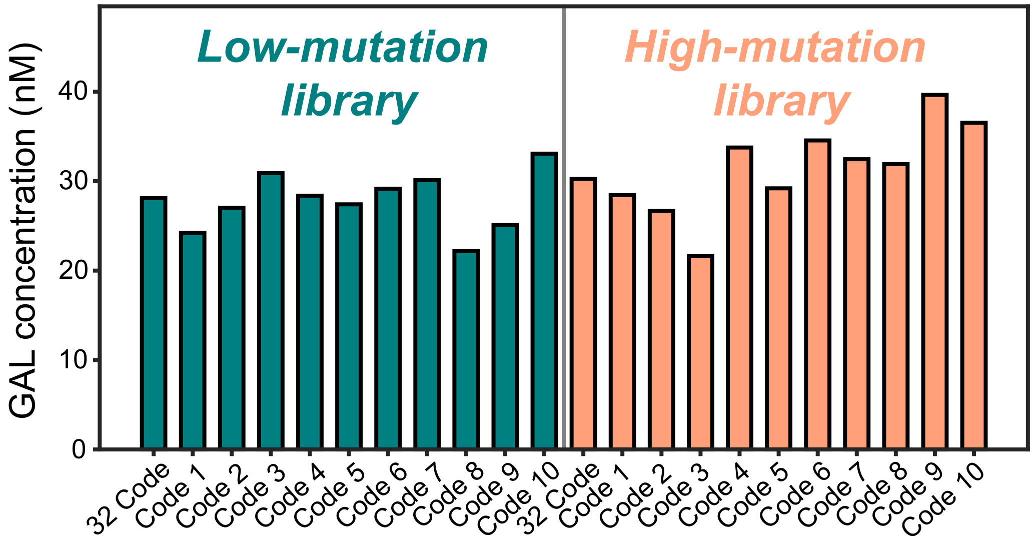


**Figure S9. Quantification of translated GAL protein concentrations across different genetic codes**

GAL protein synthesized using each low- or high-mutation library under each genetic code was quantified by the HiBiT assay. A HiBiT tag was fused to the C-terminus of the GAL gene. After translation, the HiBiT tag formed an active NanoLuc luciferase upon addition of LgBiT, and the resulting luminescence was measured. A standard curve generated using known concentrations of a HiBiT control protein was measured in parallel and used to quantify the amount of synthesized GAL protein.

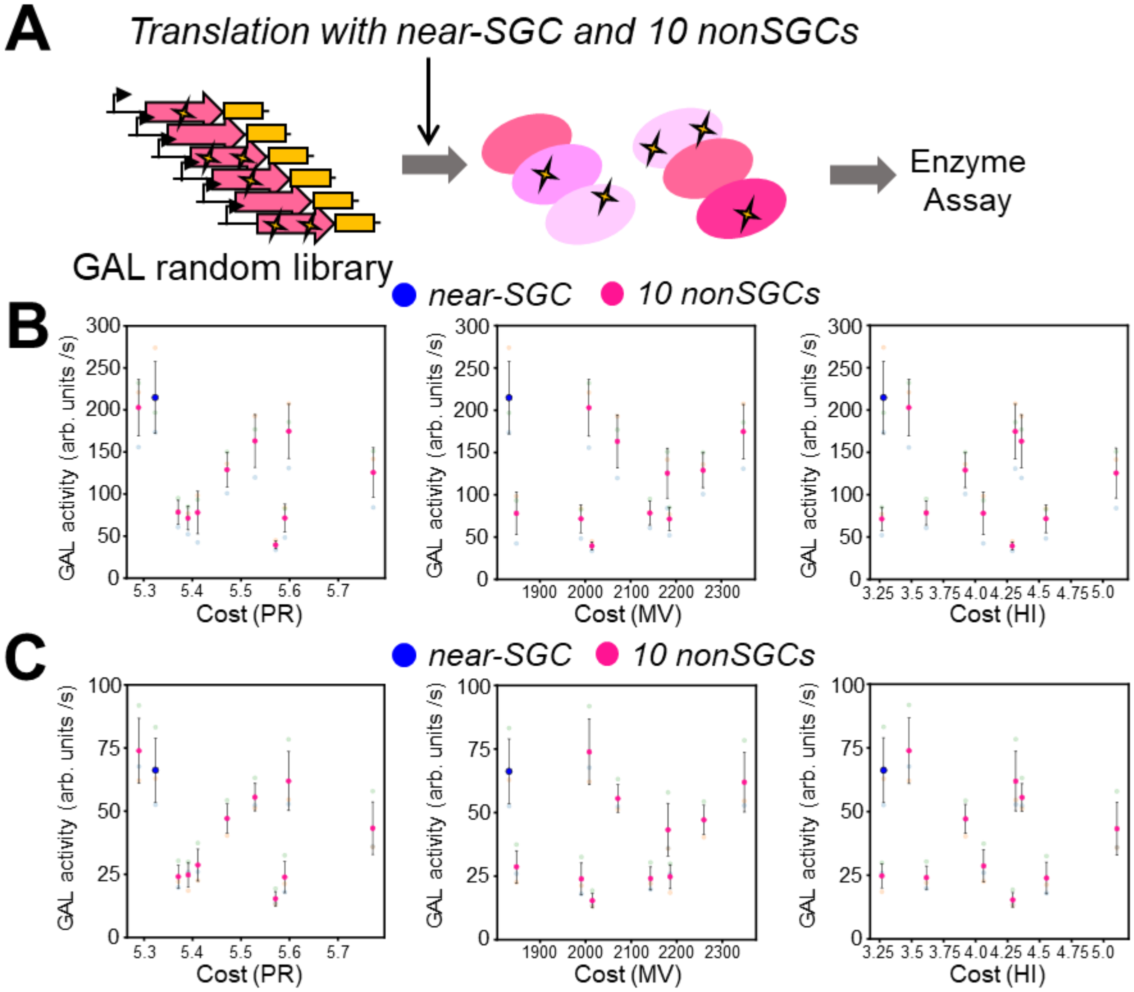


**Figure S10. Translation of GAL random library with non-SGCs**

(A) Schematic of the experimental procedure. Random DNA libraries prepared at low and high mutation rates were translated using 10 non-SGCs or near-SGC with the GAL random DNA library (5 nM). After incubation at 30°C for 16 h, GAL activity was quantified. (B, C) Distribution of GAL protein activity plotted against theoretical mutational cost for the low-mutation library (B) and the high-mutation library (C).

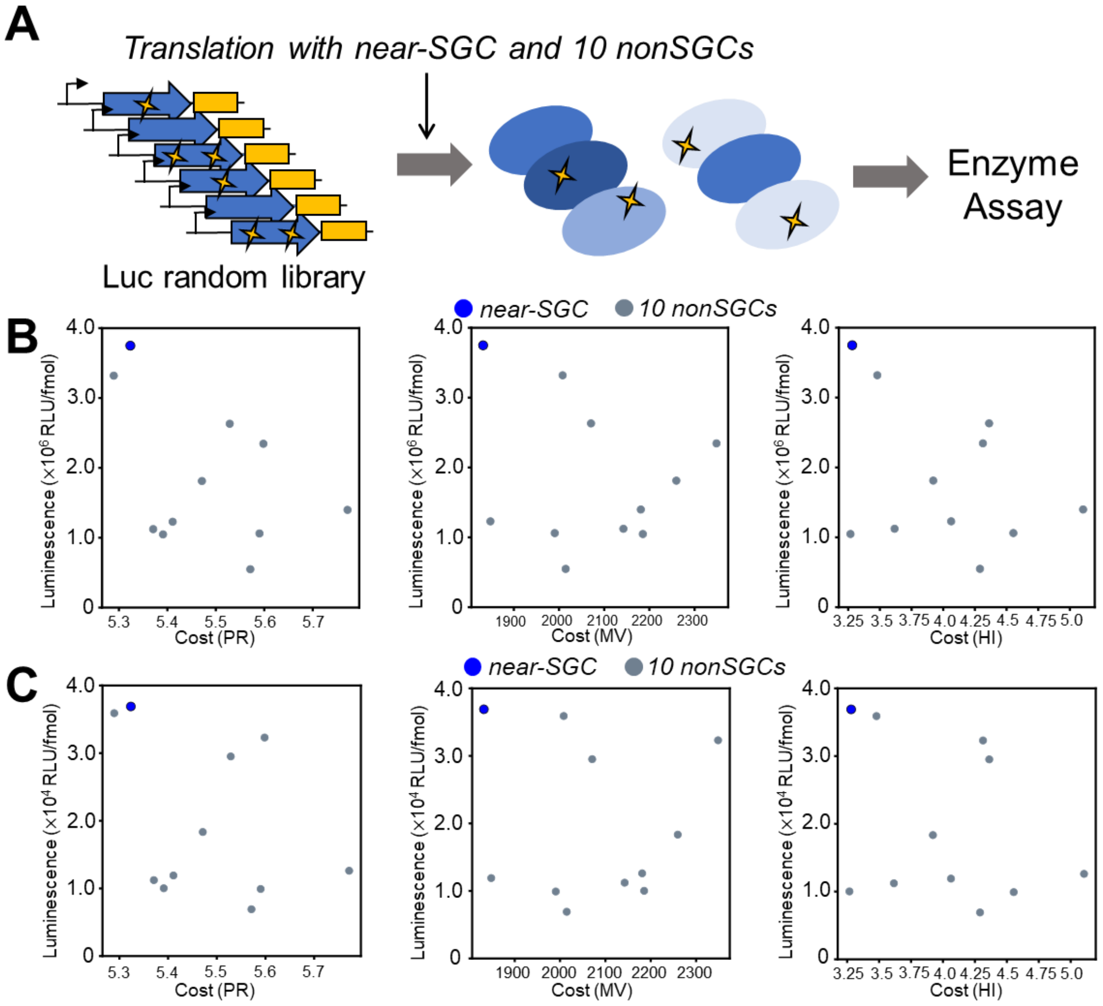


**Figure S11. Translation of Luc random library with non-SGCs**

(A) Schematic of the experimental procedure. Random DNA libraries prepared at low and high mutation rates were translated using 10 non-SGCs or near-SGC with the Luc random DNA library (5 nM). After incubation at 30°C for 16 h, luciferase activity was quantified. (B, C) Distribution of Luc protein activity plotted against theoretical mutational cost for the low-mutation library (B) and high-mutation library (C).

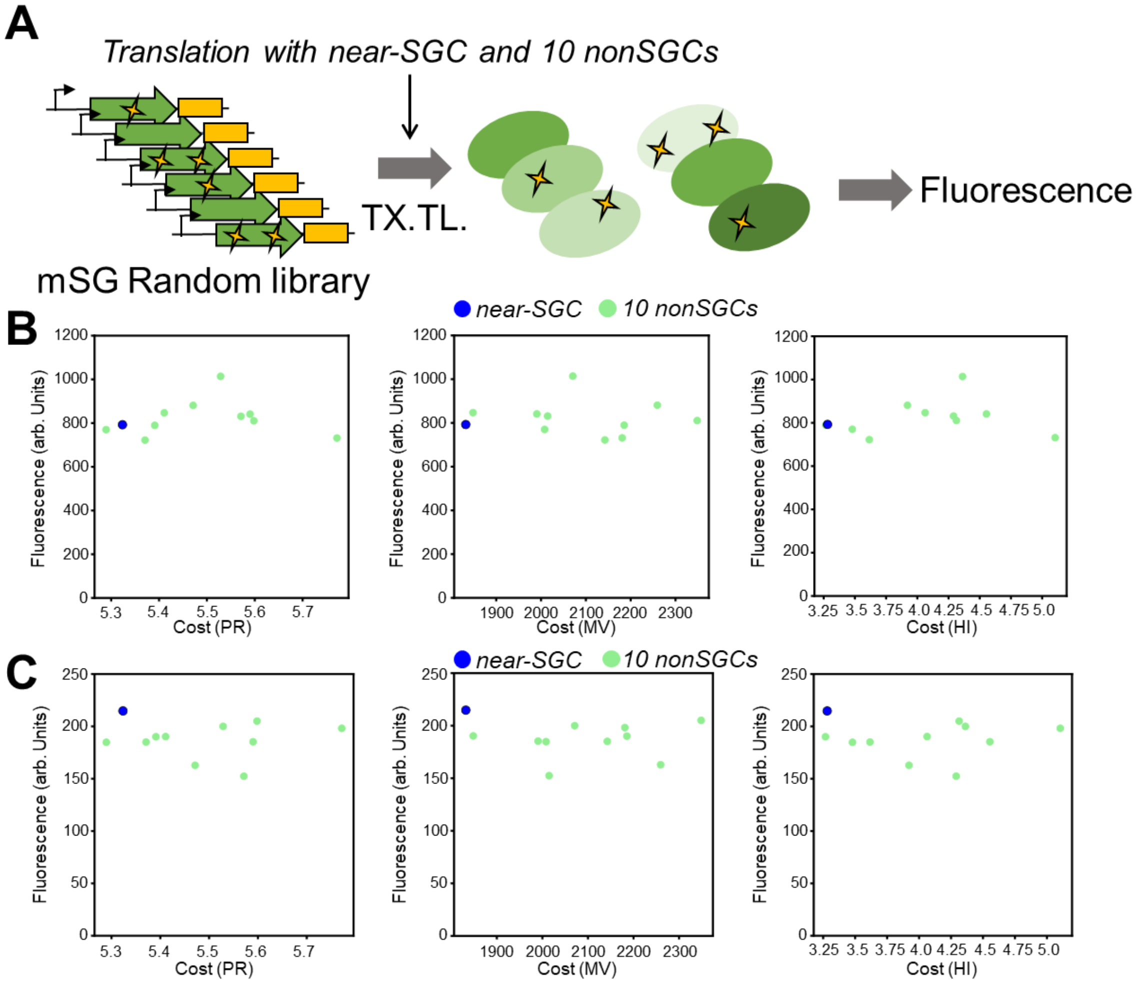


**Figure S12. Translation of mSG random library with non-SGCs**

(A) Schematic of the experimental procedure. Random DNA libraries prepared at low and high mutation rates were translated using 10 non-SGCs or near-SGC with the mSG random DNA library (5 nM). During incubation at 30°C for 16 h, mSG fluorescence was quantified. (B, C) Distribution of mSG protein activity plotted against theoretical mutational cost for the low-mutation library (B) and high-mutation library (C).

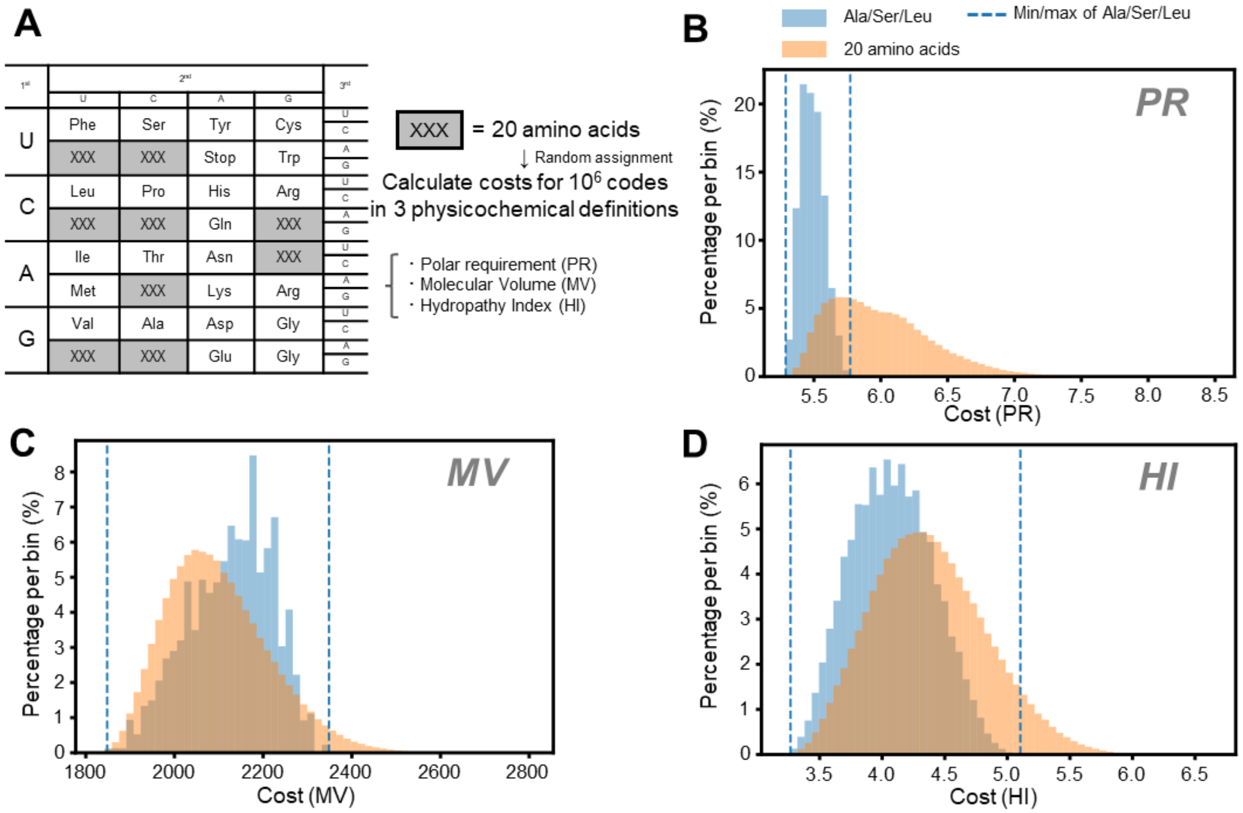


**Figure S13. Distributions of the mutational costs when assigning all 20 amino acids to the vacant codons (orange).**

(A) Calculation method of mutational costs for each genetic code based on three physicochemical properties of amino acids. For each genetic code, the average magnitude of change in amino acid physicochemical properties resulting from single-nucleotide substitutions from the 21 codons was calculated, taking mutation weighting into account (see Methods). Three physicochemical metrics were used: polar requirement (PR), molecular volume (MV), and hydropathy index (HI). In this analysis, all 20 amino acids were randomly assigned to each of the nine vacant codon boxes (gray), in contrast to the analysis in Fig. 3B, in which only three amino acids (Ala, Leu, and Ser) were assigned. Mutational costs were calculated for 1,000,000 randomly sampled genetic codes. (B, C, D) Distributions of mutational costs for each physicochemical property of amino acids when assigning 20 amino acids (orange). The distributions obtained when assigning only three amino acids (Ala, Ser, and Leu), identical to the data shown in Fig. 3B, are shown for comparison (blue). Dashed lines indicate the maximum and minimum values of the blue distribution, the cost ranges of the experimentally constructed non-SGCs in this study.

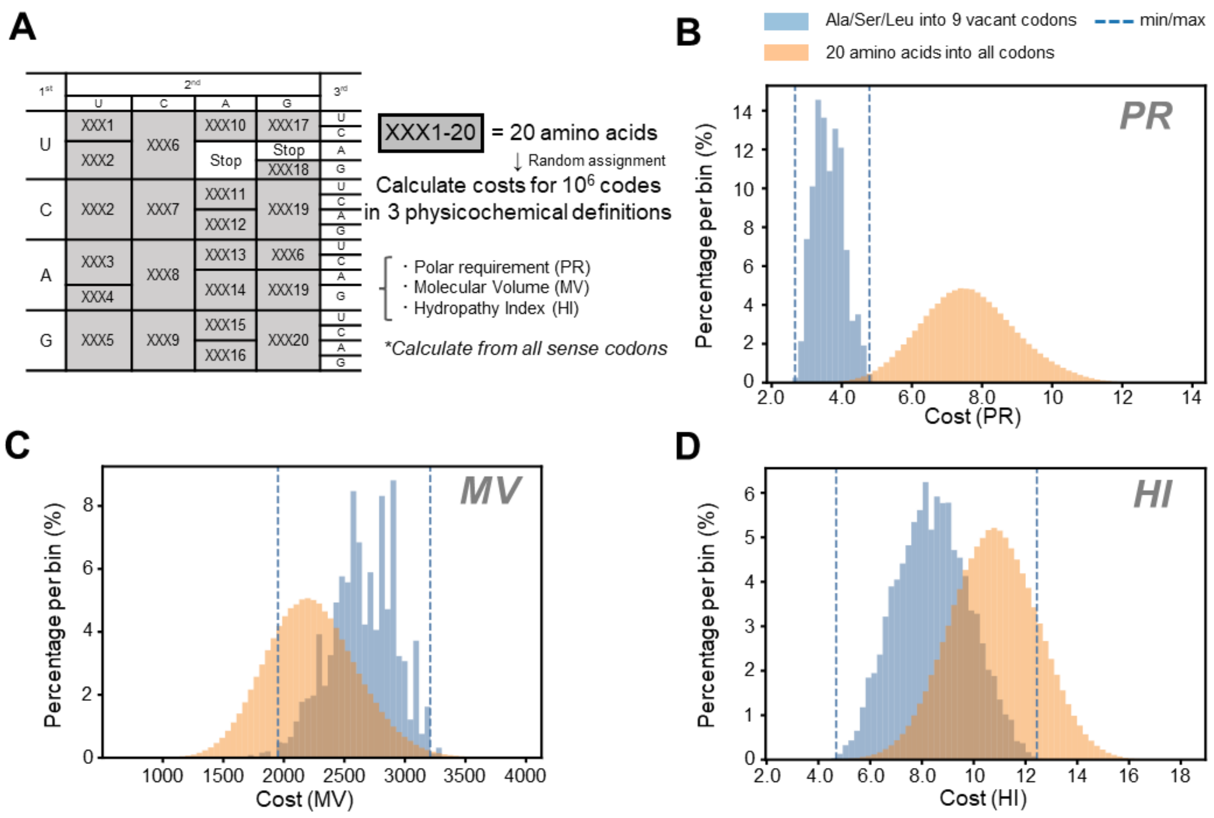


**Figure S14. Distributions of the mutational costs when assigning all 20 amino acids to all sense codons (orange).**

(A) Calculation method of mutational costs for each genetic code based on three physicochemical properties of amino acids. For each genetic code, the average magnitude of change in amino acid physicochemical properties resulting from single-nucleotide substitutions from all sense codons was calculated, taking mutation weighting into account (see Methods). Three physicochemical metrics were used: polar requirement (PR), molecular volume (MV), and hydropathy index (HI). In this analysis, all 20 amino acids were randomly assigned to each of the 20 codon boxes (gray) while preserving the degeneracy pattern of the standard genetic code. Mutational costs were calculated for 1,000,000 randomly sampled genetic codes. (B, C, D) Distributions of mutational costs for each physicochemical property of amino acids when assigning all 20 amino acids into all sense codons (orange). The distributions obtained when assigning only three amino acids (Ala, Ser, and Leu), identical to the data shown in Fig. 3B, are shown for comparison (blue). Dashed lines indicate the maximum and minimum values of the blue distribution, the cost ranges of the experimentally constructed non-SGCs in this study.

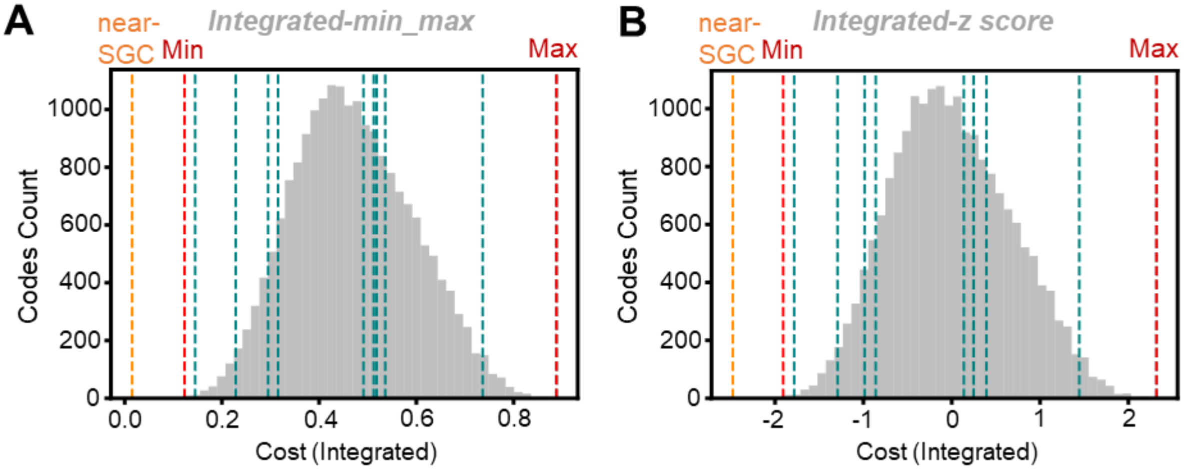


**Figure S15. Integrated mutational cost analysis combining PR, MV, and HI.**

(A) Distribution of integrated min–max costs among 19,683 candidate non-SGCs. For each cost metric, $Cost_{PR}$, $Cost_{MV}$, and $Cost_{HI}$ were min–max normalized across the candidate non-SGCs and averaged with equal weights. The orange dashed line indicates the near-SGC reference, and the red dashed line indicates the candidate non-SGC with the lowest and highest integrated cost. The green dashed lines indicate the cost values of 10 genetic codes selected for experimental construction. (B) Distribution of integrated z-score costs among candidate non-SGCs. For each metric, costs were z-score normalized using the mean and standard deviation of the candidate non-SGC distribution and averaged with equal weights.

### Table S1. DNA sequences used in this study

| Template | Sequence |
|---|---|
| Nanoluc_32C<br>*Nannoluc gene | GGCGATTAAGTTGGGTAACGCCAGGGTTTTCCCAGTCACGACGTTGTAaAACGACGGCCAGTGAATTCTAATACGACTCACTATAGGGGAATTGTGAGCGGATAACA<br>ATTCCCCTCTAGAAATAATTTTGTTTAACTTTAAGAAGGAGATATACATATGGTTTTTACTCTGGAGGACTTTGTGGGTGACTGGCGGCAGACGGCTGGGTACAACTT<br>GGACCAGGTTCTGGAGCAGGGTGGGGTGTCTTCGCTCTTTCAGAACCTGGGTGTTAGCGTGACTCCTATTCAGCGCATTGTTTTGTCTGGGGAGAACGGTCTGAAGA<br>TTGACATTCACGTGATTATTCCGTACGAGGGGCTCTCGGGTGACCAGATGGGGCAGATTGAGAAGATTTTTAAGGTTGTGTACCCTGTTGACGACCACCACTTTAAG<br>GTGATTCTGCACTACGGTACGTTGGTTATTGACGGGGTGACTCCGAACATGATTGACTACTTTGGTAGGCCTTACGAGGGGATTGCGGTTTTTGACGGTAAGAAGAT<br>TACGGTGACTGGGACGCTGTGGAACGGTAACAAGATTATTGACGAGCGGCTCATTAACCCGGACGGGAGCCTGTTGTTTCGCGTTACTATTAACGGTGTGACGGGGT<br>GGAGGCTGTGTGAGCGGATTCTCGCGTAAAAGCTTGGCGTAATCATGGTCATAGCTGTTTCCTGTGTGAAATTGTTATCCGCTCACAATTCCACACAACATACGAGC<br>CGG |
| Nanoluc_46C<br>*Nannoluc gene | GGCGATTAAGTTGGGTAACGCCAGGGTTTTCCCAGTCACGACGTTGTAAAACGACGGCCAGTGAATTCTAATACGACTCACTATAGGGGAATTGTGAGCGGATAAC<br>AATTCCCCTCTAGAAATAATTTTGTTTAACTTTAAGAAGGAGATATACATATGGTTTTTACTCTGGAGGACTTTGTGGGTGACTGGCGGCAGACGGCTGGGTACAAC<br>TTGGACCAAGTATTAGAACAGGGAGGGGTTTCTTCGCTCTTTCAAAACCTAGGTGTGAGCGTAACACCTATTCAGCGCATAGTTTATCAGGAGAGAACGGTCTGAA<br>GATTGACATACACGTGATTATACCGTACGAAGGGCTCTCGGGAGACCAAATGGGGCAGATTGAGAAAATATTTAAGGTAGTTTACCCAGTGGACGACCACCACTTT<br>AAAGTAATTCTACACTACGGTACGTTGGTTATAGACGGAGTGACTCCGAACATGATTGACTACTTTGGTAGGCCTTACGAAGGGATAGCAGTATTTGACGGAAAGA<br>AAATTACAGTTACTGGGACGCTGTGGAACGGTAACAAGATAATTGACGAGCGATTAATAAACCCAGACGGATCACTGTTGTTTAGAGTGACTATTAACGGTGTAAC<br>AGGGTGGCGACTATGTGAAAGAATACTCGCGTAAAAGCTTGGCGTAATCATGGTCATAGCTGTTTCCTGTGTGAAATTGTTATCCGCTCACAATTCCACACAACATA<br>CGAGCCGG |
| Nanoluc_21C<br>*Nannoluc gene | GGCGATTAAGTTGGGTAACGCCAGGGTTTTCCCAGTCACGACGTTGTAAAACGACGGCCAGTGAATTCTAATACGACTCACTATAGGGGAATTGTGAGCGGATAAC<br>AATTCCCCTCTAGAAATAATTTTGTTTAACTTTAAGAAGGAGATATACATATGGTTTTTACTCTGGAGGACTTTGTTGGTGACTGGCGGCAGACTGCTGGTTACAACC<br>TGGACCAGGTTCTGGAGCAGGGTGGTGTTTCTTCTCTGTTTCAGAACCTGGGTGTTTCTGTTACTCCTATTCAGCGGATTGTTCTGTCTGGTGAGAACGGTCTGAAGA<br>TTGACATTCACGTTATTATTCCTTACGAGGGTCTGTCTGGTGACCAGATGGGTCAGATTGAGAAGATTTTTAAGGTTGTTTACCCTGTTGACGACCACCACTTTAAGG<br>TTATTCTGCACTACGGTACTCTGGTTATTGACGGTGTTACTCCTAACATGATTGACTACTTTGGTCGGCCTTACGAGGGTATTGCTGTTTTTGACGGTAAGAAGATTA<br>CTGTTACTGGTACTCTGTGGAACGGTAACAAGATTATTGACGAGCGGCTGATTAACCCTGACGGTTCTCTGCTGTTTCGGGTTACTATTAACGGTGTTACTGGTTGGC<br>GGCTGTGTGAGCGGATTCTGGCTTAAAAGCTTGGCGTAATCATGGTCATAGCTGTTTCCTGTGTGAAATTGTTATCCGCTCACAATTCCACACAACATACGAGCCGG |
| Nanoluc_21+Ala_<br>CAA<br>*Nannoluc gene | GGCGATTAAGTTGGGTAACGCCAGGGTTTTCCCAGTCACGACGTTGTAAAACGACGGCCAGTGAATTCTAATACGACTCACTATAGGGGAATTGTGAGCGGATAAC<br>AATTCCCCTCTAGAAATAATTTTGTTTAACTTTAAGAAGGAGATATACATATGGTTTTTACTCTCGAGGACTTTGTTGGTGACTGGCGCCAGACTTTGGGTTACAACC<br>TCGACCAGGTTCTCGAGCAGGGTGGTGTTTCTTCTCTCTTTCAGAACCTCGGTGTTTCTGTTACTCCTATTCAGCGCATTGTTCTCTCTGGTGAGAACGGTCTCAAGAT<br>TGACATTCACGTTATTATTCCTTACGAGGGTCTCTCTGGTGACCAGATGGGTCAGATTGAGAAGATTTTTAAGGTTGTTTACCCTGTTGACGACCACCACTTTAAGGT<br>TATTCTCCACTACGGTACTCTCGTTATTGACGGTGTTACTCCTAACATGATTGACTACTTTGGTCGCCCTTACGAGGGTATTTTGGTTTTTGACGGTAAGAAGATTACT<br>GTTACTGGTACTCTCTGGAACGGTAACAAGATTATTGACGAGCGCCTCATTAACCCTGACGGTTCTCTCCTCTTTCGCGTTACTATTAACGGTGTTACTGGTTGGCGC<br>CTCTGTGAGCGCATTCTCGCTTAAAAGCTTGGCGTAATCATGGTCATAGCTGTTTCCTGTGTGAAATTGTTATCCGCTCACAATTCCACACAACATACGAGCCGG |
| Nanoluc_21+Ala_<br>CGA<br>*Nannoluc gene | GGCGATTAAGTTGGGTAACGCCAGGGTTTTCCCAGTCACGACGTTGTAAAACGACGGCCAGTGAATTCTAATACGACTCACTATAGGGGAATTGTGAGCGGATAAC<br>AATTCCCCTCTAGAAATAATTTTGTTTAACTTTAAGAAGGAGATATACATATGGTTTTTACTCTCGAGGACTTTGTTGGTGACTGGCGCCAGACTTCGGGTTACAACC<br>TCGACCAGGTTCTCGAGCAGGGTGGTGTTTCTTCTCTCTTTCAGAACCTCGGTGTTTCTGTTACTCCTATTCAGCGCATTGTTCTCTCTGGTGAGAACGGTCTCAAGAT<br>TGACATTCACGTTATTATTCCTTACGAGGGTCTCTCTGGTGACCAGATGGGTCAGATTGAGAAGATTTTTAAGGTTGTTTACCCTGTTGACGACCACCACTTTAAGGT<br>TATTCTCCACTACGGTACTCTCGTTATTGACGGTGTTACTCCTAACATGATTGACTACTTTGGTCGCCCTTACGAGGGTATTTCGGTTTTTGACGGTAAGAAGATTACT<br>GTTACTGGTACTCTCTGGAACGGTAACAAGATTATTGACGAGCGCCTCATTAACCCTGACGGTTCTCTCCTCTTTCGCGTTACTATTAACGGTGTTACTGGTTGGCGC<br>CTCTGTGAGCGCATTCTCGCTTAAAAGCTTGGCGTAATCATGGTCATAGCTGTTTCCTGTGTGAAATTGTTATCCGCTCACAATTCCACACAACATACGAGCCGG |
| Nanoluc_21+Ala_<br>CAG<br>*Nannoluc gene | GGCGATTAAGTTGGGTAACGCCAGGGTTTTCCCAGTCACGACGTTGTAAAACGACGGCCAGTGAATTCTAATACGACTCACTATAGGGGAATTGTGAGCGGATAAC<br>AATTCCCCTCTAGAAATAATTTTGTTTAACTTTAAGAAGGAGATATACATATGGTTTTTACTCTCGAGGACTTTGTTGGTGACTGGCGCCAGACTCTGGGTTACAACC<br>TCGACCAGGTTCTCGAGCAGGGTGGTGTTTCTTCTCTCTTTCAGAACCTCGGTGTTTCTGTTACTCCTATTCAGCGCATTGTTCTCTCTGGTGAGAACGGTCTCAAGAT<br>TGACATTCACGTTATTATTCCTTACGAGGGTCTCTCTGGTGACCAGATGGGTCAGATTGAGAAGATTTTTAAGGTTGTTTACCCTGTTGACGACCACCACTTTAAGGT<br>TATTCTCCACTACGGTACTCTCGTTATTGACGGTGTTACTCCTAACATGATTGACTACTTTGGTCGCCCTTACGAGGGTATTCTGGTTTTTGACGGTAAGAAGATTACT<br>GTTACTGGTACTCTCTGGAACGGTAACAAGATTATTGACGAGCGCCTCATTAACCCTGACGGTTCTCTCCTCTTTCGCGTTACTATTAACGGTGTTACTGGTTGGCGC<br>CTCTGTGAGCGCATTCTCGCTTAAAAGCTTGGCGTAATCATGGTCATAGCTGTTTCCTGTGTGAAATTGTTATCCGCTCACAATTCCACACAACATACGAGCCGG |
| Nanoluc_21+Ala_<br>CGG<br>*Nannoluc gene | GGCGATTAAGTTGGGTAACGCCAGGGTTTTCCCAGTCACGACGTTGTAAAACGACGGCCAGTGAATTCTAATACGACTCACTATAGGGGAATTGTGAGCGGATAAC<br>AATTCCCCTCTAGAAATAATTTTGTTTAACTTTAAGAAGGAGATATACATATGGTTTTTACTCTCGAGGACTTTGTTGGTGACTGGCGCCAGACTCCGGGTTACAACC<br>TCGACCAGGTTCTCGAGCAGGGTGGTGTTTCTTCTCTCTTTCAGAACCTCGGTGTTTCTGTTACTCCTATTCAGCGCATTGTTCTCTCTGGTGAGAACGGTCTCAAGAT<br>TGACATTCACGTTATTATTCCTTACGAGGGTCTCTCTGGTGACCAGATGGGTCAGATTGAGAAGATTTTTAAGGTTGTTTACCCTGTTGACGACCACCACTTTAAGGT<br>TATTCTCCACTACGGTACTCTCGTTATTGACGGTGTTACTCCTAACATGATTGACTACTTTGGTCGCCCTTACGAGGGTATTCCGGTTTTTGACGGTAAGAAGATTACT<br>GTTACTGGTACTCTCTGGAACGGTAACAAGATTATTGACGAGCGCCTCATTAACCCTGACGGTTCTCTCCTCTTTCGCGTTACTATTAACGGTGTTACTGGTTGGCGC<br>CTCTGTGAGCGCATTCTCGCTTAAAAGCTTGGCGTAATCATGGTCATAGCTGTTTCCTGTGTGAAATTGTTATCCGCTCACAATTCCACACAACATACGAGCCGG |
| Nanoluc_21+Ala_<br>CCG<br>*Nannoluc gene | GGCGATTAAGTTGGGTAACGCCAGGGTTTTCCCAGTCACGACGTTGTAAAACGACGGCCAGTGAATTCTAATACGACTCACTATAGGGGAATTGTGAGCGGATAAC<br>AATTCCCCTCTAGAAATAATTTTGTTTAACTTTAAGAAGGAGATATACATATGGTTTTTACTCTCGAGGACTTTGTTGGTGACTGGCGCCAGACTCGGGGTTACAACC<br>TCGACCAGGTTCTCGAGCAGGGTGGTGTTTCTTCTCTCTTTCAGAACCTCGGTGTTTCTGTTACTCCTATTCAGCGCATTGTTCTCTCTGGTGAGAACGGTCTCAAGAT<br>TGACATTCACGTTATTATTCCTTACGAGGGTCTCTCTGGTGACCAGATGGGTCAGATTGAGAAGATTTTTAAGGTTGTTTACCCTGTTGACGACCACCACTTTAAGGT<br>TATTCTCCACTACGGTACTCTCGTTATTGACGGTGTTACTCCTAACATGATTGACTACTTTGGTCGCCCTTACGAGGGTATTCGGGTTTTTGACGGTAAGAAGATTACT<br>GTTACTGGTACTCTCTGGAACGGTAACAAGATTATTGACGAGCGCCTCATTAACCCTGACGGTTCTCTCCTCTTTCGCGTTACTATTAACGGTGTTACTGGTTGGCGC<br>CTCTGTGAGCGCATTCTCGCTTAAAAGCTTGGCGTAATCATGGTCATAGCTGTTTCCTGTGTGAAATTGTTATCCGCTCACAATTCCACACAACATACGAGCCGG |
| Nanoluc_21+Ala_<br>CGU<br>*Nannoluc gene | GGCGATTAAGTTGGGTAACGCCAGGGTTTTCCCAGTCACGACGTTGTAAAACGACGGCCAGTGAATTCTAATACGACTCACTATAGGGGAATTGTGAGCGGATAAC<br>AATTCCCCTCTAGAAATAATTTTGTTTAACTTTAAGAAGGAGATATACATATGGTTTTTACTCTCGAGGACTTTGTTGGTGACTGGCGCCAGACTACGGGTTACAACC<br>TCGACCAGGTTCTCGAGCAGGGTGGTGTTTCTTCTCTCTTTCAGAACCTCGGTGTTTCTGTTACTCCTATTCAGCGCATTGTTCTCTCTGGTGAGAACGGTCTCAAGAT<br>TGACATTCACGTTATTATTCCTTACGAGGGTCTCTCTGGTGACCAGATGGGTCAGATTGAGAAGATTTTTAAGGTTGTTTACCCTGTTGACGACCACCACTTTAAGGT<br>TATTCTCCACTACGGTACTCTCGTTATTGACGGTGTTACTCCTAACATGATTGACTACTTTGGTCGCCCTTACGAGGGTATTACGGTTTTTGACGGTAAGAAGATTACT<br>GTTACTGGTACTCTCTGGAACGGTAACAAGATTATTGACGAGCGCCTCATTAACCCTGACGGTTCTCTCCTCTTTCGCGTTACTATTAACGGTGTTACTGGTTGGCGC<br>CTCTGTGAGCGCATTCTCGCTTAAAAGCTTGGCGTAATCATGGTCATAGCTGTTTCCTGTGTGAAATTGTTATCCGCTCACAATTCCACACAACATACGAGCCGG |
| Nanoluc_21+Ala_<br>GCU<br>*Nannoluc gene | GGCGATTAAGTTGGGTAACGCCAGGGTTTTCCCAGTCACGACGTTGTAAAACGACGGCCAGTGAATTCTAATACGACTCACTATAGGGGAATTGTGAGCGGATAAC<br>AATTCCCCTCTAGAAATAATTTTGTTTAACTTTAAGAAGGAGATATACATATGGTTTTTACTCTCGAGGACTTTGTTGGTGACTGGCGCCAGACTAGCGGTTACAACC<br>TCGACCAGGTTCTCGAGCAGGGTGGTGTTTCTTCTCTCTTTCAGAACCTCGGTGTTTCTGTTACTCCTATTCAGCGCATTGTTCTCTCTGGTGAGAACGGTCTCAAGAT<br>TGACATTCACGTTATTATTCCTTACGAGGGTCTCTCTGGTGACCAGATGGGTCAGATTGAGAAGATTTTTAAGGTTGTTTACCCTGTTGACGACCACCACTTTAAGGT<br>TATTCTCCACTACGGTACTCTCGTTATTGACGGTGTTACTCCTAACATGATTGACTACTTTGGTCGCCCTTACGAGGGTATTAGCGTTTTTGACGGTAAGAAGATTACT<br>GTTACTGGTACTCTCTGGAACGGTAACAAGATTATTGACGAGCGCCTCATTAACCCTGACGGTTCTCTCCTCTTTCGCGTTACTATTAACGGTGTTACTGGTTGGCGC<br>CTCTGTGAGCGCATTCTCGCTTAAAAGCTTGGCGTAATCATGGTCATAGCTGTTTCCTGTGTGAAATTGTTATCCGCTCACAATTCCACACAACATACGAGCCGG |
| Nanoluc_21+Ala_<br>CAC<br>*Nannoluc gene | GGCGATTAAGTTGGGTAACGCCAGGGTTTTCCCAGTCACGACGTTGTAAAACGACGGCCAGTGAATTCTAATACGACTCACTATAGGGGAATTGTGAGCGGATAAC<br>AATTCCCCTCTAGAAATAATTTTGTTTAACTTTAAGAAGGAGATATACATATGGTTTTTACTCTCGAGGACTTTGTTGGTGACTGGCGCCAGACTGTGGGTTACAACC<br>TCGACCAGGTTCTCGAGCAGGGTGGTGTTTCTTCTCTCTTTCAGAACCTCGGTGTTTCTGTTACTCCTATTCAGCGCATTGTTCTCTCTGGTGAGAACGGTCTCAAGAT<br>TGACATTCACGTTATTATTCCTTACGAGGGTCTCTCTGGTGACCAGATGGGTCAGATTGAGAAGATTTTTAAGGTTGTTTACCCTGTTGACGACCACCACTTTAAGGT<br>TATTCTCCACTACGGTACTCTCGTTATTGACGGTGTTACTCCTAACATGATTGACTACTTTGGTCGCCCTTACGAGGGTATTGTGGTTTTTGACGGTAAGAAGATTACT<br>GTTACTGGTACTCTCTGGAACGGTAACAAGATTATTGACGAGCGCCTCATTAACCCTGACGGTTCTCTCCTCTTTCGCGTTACTATTAACGGTGTTACTGGTTGGCGC<br>CTCTGTGAGCGCATTCTCGCTTAAAAGCTTGGCGTAATCATGGTCATAGCTGTTTCCTGTGTGAAATTGTTATCCGCTCACAATTCCACACAACATACGAGCCGG |
| Nanoluc_21+Ser_<br>CAA<br>*Nannoluc gene | GGCGATTAAGTTGGGTAACGCCAGGGTTTTCCCAGTCACGACGTTGTAAAACGACGGCCAGTGAATTCTAATACGACTCACTATAGGGGAATTGTGAGCGGATAAC<br>AATTCCCCTCTAGAAATAATTTTGTTTAACTTTAAGAAGGAGATATACATATGGTTTTTACTCTCGAGGACTTTGTTGGTGACTGGCGCCAGACTGCTGGTTACAACC<br>TCGACCAGGTTCTCGAGCAGGGTGGTGTTTCTTTGCTCTTTCAGAACCTCGGTGTTTCTGTTACTCCTATTCAGCGCATTGTTCTCTTGGGTGAGAACGGTCTCAAGAT<br>TGACATTCACGTTATTATTCCTTACGAGGGTCTCTCTGGTGACCAGATGGGTCAGATTGAGAAGATTTTTAAGGTTGTTTACCCTGTTGACGACCACCACTTTAAGGT<br>TATTCTCCACTACGGTACTCTCGTTATTGACGGTGTTACTCCTAACATGATTGACTACTTTGGTCGCCCTTACGAGGGTATTGCTGTTTTTGACGGTAAGAAGATTACT |

| | |
|---|---|
| | GTTACTGGTACTCTCTGGAACGGTAACAAGATTATTGACGAGCGCCTCATTAACCCTGACGGTTTGCTCCTCTTTCGCGTTACTATTAACGGTGTTACTGGTTGGCGC<br>CTCTGTGAGCGCATTCTCGCTTAAAAGCTTGGCGTAATCATGGTCATAGCTGTTTCCTGTGTGAAATTGTTATCCGCTCACAATTCCACACAACATACGAGCCGG |
| Nanoluc_21+Ser_CAG<br>*Nannoluc gene | GGCGATTAAGTTGGGTAACGCCAGGGTTTTCCCAGTCACGACGTTGTAAAACGACGGCCAGTGAATTCTAATACGACTCACTATAGGGGAATTGTGAGCGGATAAC<br>AATTCCCCTCTAGAAATAATTTTGTTTAACTTTAAGAAGGAGATATACATATGGTTTTTACTCTCGAGGACTTTGTTGGTGACTGGCGCCAGACTGCTGGTTACAACC<br>TCGACCAGGTTCTCGAGCAGGGTGGTGTTTCTCTGCTCTTTCAGAACCTCGGTGTTTCTGTTACTCCTATTCAGCGCATTGTTCTCCTGGGTGAGAACGGTCTCAAGAT<br>TGACATTCACGTTATTATTCCTTACGAGGGTCTCTCTGGTGACCAGATGGGTCAGATTGAGAAGATTTTTAAGGTTGTTTACCCTGTTGACGACCACCACTTTAAGGT<br>TATTCTCCACTACGGTACTCTCGTTATTGACGGTGTTACTCCTAACATGATTGACTACTTTGGTCGCCCTTACGAGGGTATTGCTGTTTTTGACGGTAAGAAGATTACT<br>GTTACTGGTACTCTCTGGAACGGTAACAAGATTATTGACGAGCGCCTCATTAACCCTGACGGTCTGCTCCTCTTTCGCGTTACTATTAACGGTGTTACTGGTTGGCGC<br>CTCTGTGAGCGCATTCTCGCTTAAAAGCTTGGCGTAATCATGGTCATAGCTGTTTCCTGTGTGAAATTGTTATCCGCTCACAATTCCACACAACATACGAGCCGG |
| Nanoluc_21+Ser_CGG<br>*Nannoluc gene | GGCGATTAAGTTGGGTAACGCCAGGGTTTTCCCAGTCACGACGTTGTAAAACGACGGCCAGTGAATTCTAATACGACTCACTATAGGGGAATTGTGAGCGGATAAC<br>AATTCCCCTCTAGAAATAATTTTGTTTAACTTTAAGAAGGAGATATACATATGGTTTTTACTCTCGAGGACTTTGTTGGTGACTGGCGCCAGACTGCTGGTTACAACC<br>TCGACCAGGTTCTCGAGCAGGGTGGTGTTTCTCCGCTCTTTCAGAACCTCGGTGTTTCTGTTACTCCTATTCAGCGCATTGTTCTCCCGGGTGAGAACGGTCTCAAGA<br>TTGACATTCACGTTATTATTCCTTACGAGGGTCTCTCTGGTGACCAGATGGGTCAGATTGAGAAGATTTTTAAGGTTGTTTACCCTGTTGACGACCACCACTTTAAGG<br>TTATTCTCCACTACGGTACTCTCGTTATTGACGGTGTTACTCCTAACATGATTGACTACTTTGGTCGCCCTTACGAGGGTATTGCTGTTTTTGACGGTAAGAAGATTAC<br>TGTTACTGGTACTCTCTGGAACGGTAACAAGATTATTGACGAGCGCCTCATTAACCCTGACGGTCCGCTCCTCTTTCGCGTTACTATTAACGGTGTTACTGGTTGGCG<br>CCTCTGTGAGCGCATTCTCGCTTAAAAGCTTGGCGTAATCATGGTCATAGCTGTTTCCTGTGTGAAATTGTTATCCGCTCACAATTCCACACAACATACGAGCCGG |
| Nanoluc_21+Ser_CCG<br>*Nannoluc gene | GGCGATTAAGTTGGGTAACGCCAGGGTTTTCCCAGTCACGACGTTGTAAAACGACGGCCAGTGAATTCTAATACGACTCACTATAGGGGAATTGTGAGCGGATAAC<br>AATTCCCCTCTAGAAATAATTTTGTTTAACTTTAAGAAGGAGATATACATATGGTTTTTACTCTCGAGGACTTTGTTGGTGACTGGCGCCAGACTGCTGGTTACAACC<br>TCGACCAGGTTCTCGAGCAGGGTGGTGTTTCTCGGCTCTTTCAGAACCTCGGTGTTTCTGTTACTCCTATTCAGCGCATTGTTCTCCGGGGTGAGAACGGTCTCAAGA<br>TTGACATTCACGTTATTATTCCTTACGAGGGTCTCTCTGGTGACCAGATGGGTCAGATTGAGAAGATTTTTAAGGTTGTTTACCCTGTTGACGACCACCACTTTAAGG<br>TTATTCTCCACTACGGTACTCTCGTTATTGACGGTGTTACTCCTAACATGATTGACTACTTTGGTCGCCCTTACGAGGGTATTGCTGTTTTTGACGGTAAGAAGATTAC<br>TGTTACTGGTACTCTCTGGAACGGTAACAAGATTATTGACGAGCGCCTCATTAACCCTGACGGTCGGCTCCTCTTTCGCGTTACTATTAACGGTGTTACTGGTTGGCG<br>CCTCTGTGAGCGCATTCTCGCTTAAAAGCTTGGCGTAATCATGGTCATAGCTGTTTCCTGTGTGAAATTGTTATCCGCTCACAATTCCACACAACATACGAGCCGG |
| Nanoluc_21+Ser_CGU<br>*Nannoluc gene | GGCGATTAAGTTGGGTAACGCCAGGGTTTTCCCAGTCACGACGTTGTAAAACGACGGCCAGTGAATTCTAATACGACTCACTATAGGGGAATTGTGAGCGGATAAC<br>AATTCCCCTCTAGAAATAATTTTGTTTAACTTTAAGAAGGAGATATACATATGGTTTTTACTCTCGAGGACTTTGTTGGTGACTGGCGCCAGACTGCTGGTTACAACC<br>TCGACCAGGTTCTCGAGCAGGGTGGTGTTTCTACGCTCTTTCAGAACCTCGGTGTTTCTGTTACTCCTATTCAGCGCATTGTTCTCACGGGTGAGAACGGTCTCAAGA<br>TTGACATTCACGTTATTATTCCTTACGAGGGTCTCTCTGGTGACCAGATGGGTCAGATTGAGAAGATTTTTAAGGTTGTTTACCCTGTTGACGACCACCACTTTAAGG<br>TTATTCTCCACTACGGTACTCTCGTTATTGACGGTGTTACTCCTAACATGATTGACTACTTTGGTCGCCCTTACGAGGGTATTGCTGTTTTTGACGGTAAGAAGATTAC<br>TGTTACTGGTACTCTCTGGAACGGTAACAAGATTATTGACGAGCGCCTCATTAACCCTGACGGTACGCTCCTCTTTCGCGTTACTATTAACGGTGTTACTGGTTGGCG<br>CCTCTGTGAGCGCATTCTCGCTTAAAAGCTTGGCGTAATCATGGTCATAGCTGTTTCCTGTGTGAAATTGTTATCCGCTCACAATTCCACACAACATACGAGCCGG |
| Nanoluc_21+Ser_CAC<br>*Nannoluc gene | GGCGATTAAGTTGGGTAACGCCAGGGTTTTCCCAGTCACGACGTTGTAAAACGACGGCCAGTGAATTCTAATACGACTCACTATAGGGGAATTGTGAGCGGATAAC<br>AATTCCCCTCTAGAAATAATTTTGTTTAACTTTAAGAAGGAGATATACATATGGTTTTTACTCTCGAGGACTTTGTTGGTGACTGGCGCCAGACTGCTGGTTACAACC<br>TCGACCAGGTTCTCGAGCAGGGTGGTGTTTCTGTGCTCTTTCAGAACCTCGGTGTTTCTGTTACTCCTATTCAGCGCATTGTTCTCGTGGGTGAGAACGGTCTCAAGA<br>TTGACATTCACGTTATTATTCCTTACGAGGGTCTCTCTGGTGACCAGATGGGTCAGATTGAGAAGATTTTTAAGGTTGTTTACCCTGTTGACGACCACCACTTTAAGG<br>TTATTCTCCACTACGGTACTCTCGTTATTGACGGTGTTACTCCTAACATGATTGACTACTTTGGTCGCCCTTACGAGGGTATTGCTGTTTTTGACGGTAAGAAGATTAC<br>TGTTACTGGTACTCTCTGGAACGGTAACAAGATTATTGACGAGCGCCTCATTAACCCTGACGGTGTGCTCCTCTTTCGCGTTACTATTAACGGTGTTACTGGTTGGCG<br>CCTCTGTGAGCGCATTCTCGCTTAAAAGCTTGGCGTAATCATGGTCATAGCTGTTTCCTGTGTGAAATTGTTATCCGCTCACAATTCCACACAACATACGAGCCGG |
| Nanoluc_21+Ser_CGC<br>*Nannoluc gene | GGCGATTAAGTTGGGTAACGCCAGGGTTTTCCCAGTCACGACGTTGTAAAACGACGGCCAGTGAATTCTAATACGACTCACTATAGGGGAATTGTGAGCGGATAAC<br>AATTCCCCTCTAGAAATAATTTTGTTTAACTTTAAGAAGGAGATATACATATGGTTTTTACTCTCGAGGACTTTGTTGGTGACTGGCGCCAGACTGCTGGTTACAACC<br>TCGACCAGGTTCTCGAGCAGGGTGGTGTTTCTGCGCTCTTTCAGAACCTCGGTGTTTCTGTTACTCCTATTCAGCGCATTGTTCTCGCGGGTGAGAACGGTCTCAAGA<br>TTGACATTCACGTTATTATTCCTTACGAGGGTCTCTCTGGTGACCAGATGGGTCAGATTGAGAAGATTTTTAAGGTTGTTTACCCTGTTGACGACCACCACTTTAAGG<br>TTATTCTCCACTACGGTACTCTCGTTATTGACGGTGTTACTCCTAACATGATTGACTACTTTGGTCGCCCTTACGAGGGTATTGCTGTTTTTGACGGTAAGAAGATTAC<br>TGTTACTGGTACTCTCTGGAACGGTAACAAGATTATTGACGAGCGCCTCATTAACCCTGACGGTGCGCTCCTCTTTCGCGTTACTATTAACGGTGTTACTGGTTGGCG<br>CCTCTGTGAGCGCATTCTCGCTTAAAAGCTTGGCGTAATCATGGTCATAGCTGTTTCCTGTGTGAAATTGTTATCCGCTCACAATTCCACACAACATACGAGCCGG |
| Nanoluc_21+Leu_CGA<br>*Nannoluc gene | GGCGATTAAGTTGGGTAACGCCAGGGTTTTCCCAGTCACGACGTTGTAAAACGACGGCCAGTGAATTCTAATACGACTCACTATAGGGGAATTGTGAGCGGATAAC<br>AATTCCCCTCTAGAAATAATTTTGTTTAACTTTAAGAAGGAGATATACATATGGTTTTTACTCTCGAGGACTTTGTTGGTGACTGGCGCCAGACTGCTGGTTACAACC<br>TCGACCAGGTTCTCGAGCAGGGTGGTGTTTCTTCTTCGTTTCAGAACCTCGGTGTTTCTGTTACTCCTATTCAGCGCATTGTTCTCTCTGGTGAGAACGGTCTCAAGAT<br>TGACATTCACGTTATTATTCCTTACGAGGGTTCGTCTGGTGACCAGATGGGTCAGATTGAGAAGATTTTTAAGGTTGTTTACCCTGTTGACGACCACCACTTTAAGGT<br>TATTCTCCACTACGGTACTCTCGTTATTGACGGTGTTACTCCTAACATGATTGACTACTTTGGTCGCCCTTACGAGGGTATTGCTGTTTTTGACGGTAAGAAGATTACT<br>GTTACTGGTACTCTCTGGAACGGTAACAAGATTATTGACGAGCGCTCGATTAACCCTGACGGTTCTCTCCTCTTTCGCGTTACTATTAACGGTGTTACTGGTTGGCGC<br>CTCTGTGAGCGCATTTCGGCTTAAAAGCTTGGCGTAATCATGGTCATAGCTGTTTCCTGTGTGAAATTGTTATCCGCTCACAATTCCACACAACATACGAGCCGG |
| Nanoluc_21+Leu_CGG<br>*Nannoluc gene | GGCGATTAAGTTGGGTAACGCCAGGGTTTTCCCAGTCACGACGTTGTAAAACGACGGCCAGTGAATTCTAATACGACTCACTATAGGGGAATTGTGAGCGGATAAC<br>AATTCCCCTCTAGAAATAATTTTGTTTAACTTTAAGAAGGAGATATACATATGGTTTTTACTCTCGAGGACTTTGTTGGTGACTGGCGCCAGACTGCTGGTTACAACC<br>TCGACCAGGTTCTCGAGCAGGGTGGTGTTTCTTCTCCGTTTCAGAACCTCGGTGTTTCTGTTACTCCTATTCAGCGCATTGTTCTCTCTGGTGAGAACGGTCTCAAGAT<br>TGACATTCACGTTATTATTCCTTACGAGGGTCCGTCTGGTGACCAGATGGGTCAGATTGAGAAGATTTTTAAGGTTGTTTACCCTGTTGACGACCACCACTTTAAGGT<br>TATTCTCCACTACGGTACTCTCGTTATTGACGGTGTTACTCCTAACATGATTGACTACTTTGGTCGCCCTTACGAGGGTATTGCTGTTTTTGACGGTAAGAAGATTACT<br>GTTACTGGTACTCTCTGGAACGGTAACAAGATTATTGACGAGCGCCCGATTAACCCTGACGGTTCTCTCCTCTTTCGCGTTACTATTAACGGTGTTACTGGTTGGCGC<br>CTCTGTGAGCGCATTCCGGCTTAAAAGCTTGGCGTAATCATGGTCATAGCTGTTTCCTGTGTGAAATTGTTATCCGCTCACAATTCCACACAACATACGAGCCGG |
| Nanoluc_21+Leu_CCG<br>*Nannoluc gene | GGCGATTAAGTTGGGTAACGCCAGGGTTTTCCCAGTCACGACGTTGTAAAACGACGGCCAGTGAATTCTAATACGACTCACTATAGGGGAATTGTGAGCGGATAAC<br>AATTCCCCTCTAGAAATAATTTTGTTTAACTTTAAGAAGGAGATATACATATGGTTTTTACTCTCGAGGACTTTGTTGGTGACTGGCGCCAGACTGCTGGTTACAACC<br>TCGACCAGGTTCTCGAGCAGGGTGGTGTTTCTTCTCGGTTTCAGAACCTCGGTGTTTCTGTTACTCCTATTCAGCGCATTGTTCTCTCTGGTGAGAACGGTCTCAAGAT<br>TGACATTCACGTTATTATTCCTTACGAGGGTCGGTCTGGTGACCAGATGGGTCAGATTGAGAAGATTTTTAAGGTTGTTTACCCTGTTGACGACCACCACTTTAAGGT<br>TATTCTCCACTACGGTACTCTCGTTATTGACGGTGTTACTCCTAACATGATTGACTACTTTGGTCGCCCTTACGAGGGTATTGCTGTTTTTGACGGTAAGAAGATTACT<br>GTTACTGGTACTCTCTGGAACGGTAACAAGATTATTGACGAGCGCCGGATTAACCCTGACGGTTCTCTCCTCTTTCGCGTTACTATTAACGGTGTTACTGGTTGGCGC<br>CTCTGTGAGCGCATTCGGGCTTAAAAGCTTGGCGTAATCATGGTCATAGCTGTTTCCTGTGTGAAATTGTTATCCGCTCACAATTCCACACAACATACGAGCCGG |
| Nanoluc_21+Leu_GCU<br>*Nannoluc gene | GGCGATTAAGTTGGGTAACGCCAGGGTTTTCCCAGTCACGACGTTGTAAAACGACGGCCAGTGAATTCTAATACGACTCACTATAGGGGAATTGTGAGCGGATAAC<br>AATTCCCCTCTAGAAATAATTTTGTTTAACTTTAAGAAGGAGATATACATATGGTTTTTACTCTCGAGGACTTTGTTGGTGACTGGCGCCAGACTGCTGGTTACAACC<br>TCGACCAGGTTCTCGAGCAGGGTGGTGTTTCTTCTAGCTTTCAGAACCTCGGTGTTTCTGTTACTCCTATTCAGCGCATTGTTCTCTCTGGTGAGAACGGTCTCAAGAT<br>TGACATTCACGTTATTATTCCTTACGAGGGTAGCTCTGGTGACCAGATGGGTCAGATTGAGAAGATTTTTAAGGTTGTTTACCCTGTTGACGACCACCACTTTAAGGT<br>TATTCTCCACTACGGTACTCTCGTTATTGACGGTGTTACTCCTAACATGATTGACTACTTTGGTCGCCCTTACGAGGGTATTGCTGTTTTTGACGGTAAGAAGATTACT<br>GTTACTGGTACTCTCTGGAACGGTAACAAGATTATTGACGAGCGCAGCATTAACCCTGACGGTTCTCTCCTCTTTCGCGTTACTATTAACGGTGTTACTGGTTGGCGC<br>CTCTGTGAGCGCATTAGCGCTTAAAAGCTTGGCGTAATCATGGTCATAGCTGTTTCCTGTGTGAAATTGTTATCCGCTCACAATTCCACACAACATACGAGCCGG |
| Nanoluc_21+Leu_CGU<br>*Nannoluc gene | GGCGATTAAGTTGGGTAACGCCAGGGTTTTCCCAGTCACGACGTTGTAAAACGACGGCCAGTGAATTCTAATACGACTCACTATAGGGGAATTGTGAGCGGATAAC<br>AATTCCCCTCTAGAAATAATTTTGTTTAACTTTAAGAAGGAGATATACATATGGTTTTTACTCTCGAGGACTTTGTTGGTGACTGGCGCCAGACTGCTGGTTACAACC<br>TCGACCAGGTTCTCGAGCAGGGTGGTGTTTCTTCTACGTTTCAGAACCTCGGTGTTTCTGTTACTCCTATTCAGCGCATTGTTCTCTCTGGTGAGAACGGTCTCAAGAT<br>TGACATTCACGTTATTATTCCTTACGAGGGTACGTCTGGTGACCAGATGGGTCAGATTGAGAAGATTTTTAAGGTTGTTTACCCTGTTGACGACCACCACTTTAAGGT<br>TATTCTCCACTACGGTACTCTCGTTATTGACGGTGTTACTCCTAACATGATTGACTACTTTGGTCGCCCTTACGAGGGTATTGCTGTTTTTGACGGTAAGAAGATTACT<br>GTTACTGGTACTCTCTGGAACGGTAACAAGATTATTGACGAGCGCACGATTAACCCTGACGGTTCTCTCCTCTTTCGCGTTACTATTAACGGTGTTACTGGTTGGCGC<br>CTCTGTGAGCGCATTACGGCTTAAAAGCTTGGCGTAATCATGGTCATAGCTGTTTCCTGTGTGAAATTGTTATCCGCTCACAATTCCACACAACATACGAGCCGG |
| Nanoluc_21+Leu_CAC<br>*Nannoluc gene | GGCGATTAAGTTGGGTAACGCCAGGGTTTTCCCAGTCACGACGTTGTAAAACGACGGCCAGTGAATTCTAATACGACTCACTATAGGGGAATTGTGAGCGGATAAC<br>AATTCCCCTCTAGAAATAATTTTGTTTAACTTTAAGAAGGAGATATACATATGGTTTTTACTCTCGAGGACTTTGTTGGTGACTGGCGCCAGACTGCTGGTTACAACC<br>TCGACCAGGTTCTCGAGCAGGGTGGTGTTTCTTCTGTGTTTCAGAACCTCGGTGTTTCTGTTACTCCTATTCAGCGCATTGTTCTCTCTGGTGAGAACGGTCTCAAGAT<br>TGACATTCACGTTATTATTCCTTACGAGGGTGTGTCTGGTGACCAGATGGGTCAGATTGAGAAGATTTTTAAGGTTGTTTACCCTGTTGACGACCACCACTTTAAGGT<br>TATTCTCCACTACGGTACTCTCGTTATTGACGGTGTTACTCCTAACATGATTGACTACTTTGGTCGCCCTTACGAGGGTATTGCTGTTTTTGACGGTAAGAAGATTACT<br>GTTACTGGTACTCTCTGGAACGGTAACAAGATTATTGACGAGCGCGTGATTAACCCTGACGGTTCTCTCCTCTTTCGCGTTACTATTAACGGTGTTACTGGTTGGCGC<br>CTCTGTGAGCGCATTGTGGCTTAAAAGCTTGGCGTAATCATGGTCATAGCTGTTTCCTGTGTGAAATTGTTATCCGCTCACAATTCCACACAACATACGAGCCGG |
| Nanoluc_21+Leu_CGC<br>*Nannoluc gene | GGCGATTAAGTTGGGTAACGCCAGGGTTTTCCCAGTCACGACGTTGTAAAACGACGGCCAGTGAATTCTAATACGACTCACTATAGGGGAATTGTGAGCGGATAAC<br>AATTCCCCTCTAGAAATAATTTTGTTTAACTTTAAGAAGGAGATATACATATGGTTTTTACTCTCGAGGACTTTGTTGGTGACTGGCGCCAGACTGCTGGTTACAACC<br>TCGACCAGGTTCTCGAGCAGGGTGGTGTTTCTTCTGCGTTTCAGAACCTCGGTGTTTCTGTTACTCCTATTCAGCGCATTGTTCTCTCTGGTGAGAACGGTCTCAAGAT<br>TGACATTCACGTTATTATTCCTTACGAGGGTGCGTCTGGTGACCAGATGGGTCAGATTGAGAAGATTTTTAAGGTTGTTTACCCTGTTGACGACCACCACTTTAAGGT |

| | |
|---|---|
| | TATTCTCCACTACGGTACTCTCGTTATTGACGGTGTTACTCCTAACATGATTGACTACTTTGGTCGCCCTTACGAGGGTATTGCTGTTTTTGACGGTAAGAAGATTACT<br>GTTACTGGTACTCTCTGGAACGGTAACAAGATTATTGACGAGCGCGCGATTAACCCTGACGGTTCTCTCCTCTTTCGCGTTACTATTAACGGTGTTACTGGTTGGCGC<br>CTCTGTGAGCGCATTGCGGCTTAAAAGCTTGGCGTAATCATGGTCATAGCTGTTTCCTGTGTGAAATTGTTATCCGCTCACAATTCCACACAACATACGAGCCGG |
| GAL_21C<br>*GAL gene | GGCGATTAAGTTGGGTAACGCCAGGGTTTTCCCAGTCACGACGTTGTAAAACGACGGCCAGTGAATTCTAATACGACTCACTATAGGGGAATTGTGAGCGGATAAC<br>AATTCCCCTCTAGAAATAATTTTGTTTAACTTTAAGAAGGAGATATACATATGACTATGATTACTGACTCTCTCGCTGTTGTTCTCCAGCGCCGCGACTGGGAGAACC<br>CTGGTGTTACTCAGCTCAACCGCCTCGCTGCTCACCCTCCTTTTGCTTCTTGGCGCAACTCTGAGGAGGCTCGCACTGACCGCCCTTCTCAGCAGCTCCGCTCTCTCAA<br>CGGTGAGTGGCGCTTTGCTTGGTTTCCTGCTCCTGAGGCTGTTCCTGAGTCTTGGCTCGAGTGTGACCTCCCTGAGGCTGACACTGTTGTTGTTCCTTCTAACTGGCAG<br>ATGCACGGTTACGACGCTCCTATTTACACTAACGTTACTTACCCTATTACTGTTAACCCTCCTTTTGTTCCTACTGAGAACCCTACTGGTTGTTACTCTCTCACTTTTAA<br>CGTTGACGAGTCTTGGCTCCAGGAGGGTCAGACTCGCATTATTTTTGACGGTGTTAACTCTGCTTTTCACCTCTGGTGTAACGGTCGCTGGGTTGGTTACGGTCAGGA<br>CTCTCGCCTCCCTTCTGAGTTTGACCTCTCTGCTTTTCTCCGCGCTGGTGAGAACCGCCTCGCTGTTATGGTTCTCCGCTGGTCTGACGGTTCTTACCTCGAGGACCAG<br>GACATGTGGCGCATGTCTGGTATTTTTCGCGACGTTTCTCTCCTCCACAAGCCTACTACTCAGATTTCTGACTTTCACGTTGCTACTCGCTTTAACGACGACTTTTCTC<br>GCGCTGTTCTCGAGGCTGAGGTTCAGATGTGTGGTGAGCTCCGCGACTACCTCCGCGTTACTGTTTCTCTCTGGCAGGGTGAGACTCAGGTTGCTTCTGGTACTGCTC<br>CTTTTGGTGGTGAGATTATTGACGAGCGCGGTGGTTACGCTGACCGCGTTACTCTCCGCCTCAACGTTGAGAACCCTAAGCTCTGGTCTGCTGAGATTCCTAACCTCT<br>ACCGCGCTGTTGTTGAGCTCCACACTGCTGACGGTACTCTCATTGAGGCTGAGGCTTGTGACGTTGGTTTTCGCGAGGTTCGCATTGAGAACGGTCTCCTCCTCCTCA<br>ACGGTAAGCCTCTCCTCATTCGCGGTGTTAACCGCCACGAGCACCACCCTCTCCACGGTCAGGTTATGGACGAGCAGACTATGGTTCAGGACATTCTCCTCATGAAG<br>CAGAACAACTTTAACGCTGTTCGCTGTTCTCACTACCCTAACCACCCTCTCTGGTACACTCTCTGTGACCGCTACGGTCTCTACGTTGTTGACGAGGCTAACATTGAG<br>ACTCACGGTATGGTTCCTATGAACCGCCTCACTGACGACCCTCGCTGGCTCCCTGCTATGTCTGAGCGCGTTACTCGCATGGTTCAGCGCGACCGCAACCACCCTTCT<br>GTTATTATTTGGTCTCTCGGTAACGAGTCTGGTCACGGTGCTAACCACGACGCTCTCTACCGCTGGATTAAGTCTGTTGACCCTTCTCGCCCTGTTCAGTACGAGGGT<br>GGTGGTGCTGACACTACTGCTACTGACATTATTTGTCCTATGTACGCTCGCGTTGACGAGGACCAGCCTTTTCCTGCTGTTCCTAAGTGGTCTATTAAGAAGTGGCTC<br>TCTCTCCCTGGTGAGACTCGCCCTCTCATTCTCTGTGAGTACGCTCACGCTATGGGTAACTCTCTCGGTGGTTTTGCTAAGTACTGGCAGGCTTTTCGCCAGTACCCTC<br>GCCTCCAGGGTGGTTTTGTTTGGGACTGGGTTGACCAGTCTCTCATTAAGTACGACGAGAACGGTAACCCTTGGTCTGCTTACGGTGGTGACTTTGGTGACACTCCTA<br>ACGACCGCCAGTTTTGTATGAACGGTCTCGTTTTTGCTGACCGCACTCCTCACCCTGCTCTCACTGAGGCTAAGCACCAGCAGCAGTTTTTCAGTTTCGCCTCTCTGG<br>TCAGACTATTGAGGTTACTTCTGAGTACCTCTTTCGCCACTCTGACAACGAGCTCCTCCACTGGATGGTTGCTCTCGACGGTAAGCCTCTCGCTTCTGGTGAGGTTCC<br>TCTCGACGTTGCTCCTCAGGGTAAGCAGCTCATTGAGCTCCCTGAGCTCCCTCAGCCTGAGTCTGCTGGTCAGCTCTGGCTCACTGTTCGCGTTGTTCAGCCTAACGC<br>TACTGCTTGGTCTGAGGCTGGTCACATTTCTGCTTGGCAGCAGTGGCGCCTCGCTGAGAACCTCTCTGTTACTCTCCCTGCTGCTTCTCACGCTATTCCTCACCTCACT<br>ACTTCTGAGATGGACTTTTGTATTGAGCTCGGTAACAAGCGCTGGCAGTTTAACCGCCAGTCTGGTTTTCTCTCTCAGATGTGGATTGGTGACAAGAAGCAGCTCCTC<br>ACTCCTCTCCGCGACCAGTTTACTCGCGCTCCTCTCGACAACGACATTGGTGTTTCTGAGGCTACTCGCATTGACCCTAACGCTTGGGTTGAGCGCTGGAAGGCTGCT<br>GGTCACTACCAGGCTGAGGCTGCTCTCCTCCAGTGTACTGCTGACACTCTCGCTGACGCTGTTCTCATTACTACTGCTCACGCTTGGCAGCACCAGGGTAAGACTCTC<br>TTTATTTCTCGCAAGACTTACCGCATTGACGGTTCTGGTCAGATGGCTATTACTGTTGACGTTGAGGTTGCTTCTGACACTCCTCACCCTGCTCGCATTGGTCTCAACT<br>GTCAGCTCGCTCAGGTTGCTGAGCGCGTTAACTGGCTCGGTCTCGGTCCTCAGGAGAACTACCCTGACCGCCTCACTGCTGCTTGTTTTGACCGCTGGGACCTCCCTC<br>TCTCTGACATGTACACTCCTTACGTTTTTCCTTCTGAGAACGGTCTCCGCTGTGGTACTCGCGAGCTCAACTACGGTCCTCACCAGTGGCGCGGTGACTTTCAGTTTA<br>ACATTTCTCGCTACTCTCAGCAGCAGCTCATGGAGACTTCTCACCGCCACCTCCTCCACGCTGAGGAGGGTACTTGGCTCAACATTGACGGTTTTCACATGGGTATTG<br>GTGGTGACGACTCTTGGTCTCCTTCTGTTTCTGCTGAGTTTCAGCTCTCTGCTGGTCGCTACCACTACCAGCTCGTTTGGTGTCAGAAGTAAAAAAGCTTGGCGTAAT<br>CATGGTCATAGCTGTTTCCTGTGTGAAATTGTTATCCGCTCACAATTCCACACAACATACGAGCCGG |
| Luciferase_21C<br>*Luciferase gene | GGCGATTAAGTTGGGTAACGCCAGGGTTTTGTTTAACTTTAAGAAGGAGATATACATATGGAGGACGCTTCCCAGTCACGACGTTGTAAAACGACGGCCAGTGAA<br>TTCTAATACGACTCACTATAGGGGAATTGTGAGCGGATAACAATTCCCCTCTAGAAATAATTTTGTTTAACTTTAAGAAGGAGATATACATATGGAGGACGCTAAGA<br>ACATTAAGAAGGGTCCTGCTCCTTTTTACCCTCTCGAGGACGGTACTGCTGGTGAGCAGCTCCACAAGGCTATGAAGCGCTACGCTCTCGTTCCTGGTACTATTGCTT<br>TTACTGACGCTCACATTGAGGTTAACATTACTTACGCTGAGTACTTTGAGATGTCTGTTCGCCTCGCTGAGGCTATGAAGCGCTACGGTCTCAACACTAACCACCGCA<br>TTGTTGTTTGTTCTGAGAACTCTCTCCAGTTTTTATGCCTGTTCTCGGTGCTCTCTTTATTGGTGTTGCTGTTGCTCCTGCTAACGACATTTACAACGAGCGCGAGCTC<br>CTCAACTCTATGAACATTTCTCAGCCTACTGTTGTTTTTGTTTCTAAGAAGGGTCTCCAGAAGATTCTCAACGTTCAGAAGAAGCTCCCTATTATTCAGAAGATTATT<br>ATTATGGACTCTAAGACTGACTACCAGGGTTTTCAGTCTATGTACACTTTTGTTACTTCTCACCTCCCTCCTGGTTTTAACGAGTACGACTTTGTTCCTGAGTCTTTTG<br>ACCGCGACAAGACTATTGCTCTCATTATGAACTCTTCTGGTTCTACTGGTCTCCCTAAGGGTGTTGCTCTCCCTCACCGCACTGCTTGTGTTCGCTTTTCTCACGCTCG<br>CGACCCTATTTTTGGTAACCAGATTATTCCTGACACTGCTATTCTCTCTGTTGTTCCTTTTCACCACGGTTTTGGTATGTTTACTACTCTCGGTTACCTCATTTGTGGTT<br>TTCGCGTTGTTCTCATGTACCGCTTTGAGGAGGAGCTCTTTCTCCGCTCTCTCCAGGACTACAAGATTCAGTCTGCTCTCCTCGTTCCTACTCTCTTTTCTTTTTTTGCT<br>AAGTCTACTCTCATTGACAAGTACGACCTCTCTAACCTCCACGAGATTGCTTCTGGTGGTGCTCCTCTCTCTAAGGAGGTTGGTGAGGCTGTTGCTAAGCGCTTTCAC<br>CTCCCTGGTATTCGCCAGGGTTACGGTCTCACTGAGACTACTTCTGCTATTCTCATTACTCCTGAGGGTGACGACAAGCCTGGTGCTGTTGGTAAGGTTGTTCCTTTTT<br>TTGAGGCTAAGGTTGTTGACCTCGACACTGGTAAGACTCTCGGTGTTAACCAGCGCGGTGAGCTCTGTGTTCGCGGTCCTATGATTATGTCTGGTTACGTTAACAACC<br>CTGAGGCTACTAACGCTCTCATTGACAAGGACGGTTGGCTCCACTCTGGTGACATTGCTTACTGGGACGAGGACGAGCACTTTTTTATTGTTGACCGCCTCAAGTCTC<br>TCATTAAGTACAAGGGTTACCAGGTTGCTCCTGCTGAGCTCGAGTCTATTCTCCTCCAGCACCCTAACATTTTTGACGCTGGTGTTGCTGGTCTCCCTGACGACGACG<br>CTGGTGAGCTCCCTGCTGCTGTTGTTGTTCTCGAGCACGGTAAGACTATGACTGAGAAGGAGATTGTTGACTACGTTGCTTCTCAGGTTACTACTGCTAAGAAGCTCC<br>GCGGTGGTGTTGTTTTTGTTGACGAGGTTCCTAAGGGTCTCACTGGTAAGCTCGACGCTCGCAAGATTCGCGAGATTCTCATTAAGGCTAAGAAGGGTGGTAAGTCT<br>AAGCTCTAAAAAAGCTTGGCGTAATCATGGTCATAGCTGTTTCCTGTGTGAAATTGTTATCCGCTCACAATTCCACACAACATACGAGCCGG |
| mStayGold_21C<br>*mStayGold gene | GGCGATTAAGTTGGGTAACGCCAGGGTTTTCCCAGTCACGACGTTGTAAAACGACGGCCAGTGAATTCTAATACGACTCACTATAGGGGAATTGTGAGCGGATAAC<br>AATTCCCCTCTAGAAATAATTTTGTTTAACTTTAAGAAGGAGATATACATATGGTTTCTACTGGTGAGGAGCTCTTTACTGGTGTTGTTCCTTTTAAGTTTCAGCTCAA<br>GGGTACTATTAACGGTAAGTCTTTTACTGTTGAGGGTGAGGGTGAGGGTAACTCTCACGAGGGTTCTCACAAGGGTAAGTACGTTTGTACTTCTGGTAAGCTCCCTA<br>TGTCTTGGGCTGCTCTCGGTACTTCTTTTGGTTACGGTATGAAGTACTACACTAAGTACCCTTCTGGTCTCAAGAACTGGTTTCACGAGGTTATGCCTGAGGGTTTTAC<br>TTACGACCGCCACATTCAGTACAAGGGTGACGGTTCTATTCACGCTAAGCACCAGCACTTTATGAAGAACGGTACTTACCACAACATTGTTGAGTTTACTGGTCAGG<br>ACTTTAAGGAGAACTCTCCTGTTCTCACTGGTGACATGGACGTTTCTCTCCCTAACGAGGTTCAGCACATTCCTATTGACGACGGTGTTGAGTGTACTGTTACTCTCC<br>AGTACCCTCTCCTCTCTGACGAGTCTAAGTGTGTTGAGGCTTACCAGAACACTATTATTAAGCCTCTCCACAACCAGCCTGCTCCTGACGTTCCTTTTCACTGGATTC<br>GCAAGCAGTACACTCAGTCTAAGGACGACACTGAGGAGCGCGACCACATTATTCAGTCTGAGACTCTCGAGGCTCACCTCTAAAAAAGCTTGGCGTAATCATGGTC<br>ATAGCTGTTTCCTGTGTGAAATTGTTATCCGCTCACAATTCCACACAACATACGAGCCGG |

### Table S2. Primers used in this study

| Primer No. | Sequence |
|---|---|
| 1 | GGCGATTAAGTTGGGTAACGCCAG |
| 2 | CCGGCTCGTATGTTGTGTGG |
| 3 | GGTTCTGGTGGTAACTCTGGTTCTTCTGGTGGTTCTTCTGGTGTTTCTGGTTGGC |
| 4 | GGCCGCAAGCCTATTAAGAAATCTTCTTAAAGAGGCGCCAACCAGAAACACCAGAAG |
| 5 | GAACCAGAGTTACCACCAGAACCCTTCTGACACCAAACGAGCTGG |
| 6 | GAACCAGAGTTACCACCAGAACCGAGCTTAGACTTACCACCCTTCTTAGC |
| 7 | GAACCAGAGTTACCACCAGAACCGAGGTGAGCCTCGAGAGTCTC |
| 8 | GGCCGCAAGCCTATTAAGAAATC |

| | |
|---|---|
| 9 | CCGCGTAATACGACTCACTATAGGGGCTATAGCTCAGCTG |
| 10 | TGGTGGAGCTAAGCGGGATCG |
| 11 | ATGCAAGCGCTCTCCCAGC |
| 12 | GGAGAGCGCTTGCATCGCATGCAAGAGGTCAG |
| 13 | CCGCGTAATACGACTCACTATAGCGCCCGTAGCTCAG |
| 14 | TGGCGCGCCCGACAGGATTCG |
| 15 | AGGGCAGCGCTCTATCCAGCTGAG |
| 16 | ATAGAGCGCTGCCCTGCGGAGGCAGAGG |
| 17 | ATAGAGCGCTGCCCTCCTGAGGCAGAGGTC |
| 18 | CCGCGTAATACGACTCACTATAGGAGCGGTAGTTCAGTCG |
| 19 | TGGCGGAACGGACGGGACTCG |
| 20 | CCGCGTAATACGACTCACTATAGGCGCGTTAACAAAGCG |
| 21 | TGGAGGCGCGTTCCGGAGTCG |
| 22 | CCGCGTAATACGACTCACTATAGCGGGAATAGCTCAGTTG |
| 23 | TGGAGCGGGAAACGAGAC |
| 24 | AAGGTCGTGCTCTACCAACTGAGCTATTC |
| 25 | GTAGAGCACGACCTTCCCAAGGTCGGGGTC |
| 26 | CCGCGTAATACGACTCACTATAGTCCCCTTCGTCTAGAGG |
| 27 | TGGCGTCCCCTAGGGGATTCG |
| 28 | CCGCGTAATACGACTCACTATAGGTGGCTATAGCTCAGTTG |
| 29 | TGGGGTGGCTAATGGGATTCG |
| 30 | CCGCGTAATACGACTCACTATAGCGAAGGTGGCGGAATTG |
| 31 | TGGTGCGAGGGGGGGGA |
| 32 | AAGCTAGCGCGTCTACCAATTCCGC |
| 33 | TAGACGCGCTAGCTTCAAGTGTTAGTGTCCTTAC |
| 34 | TAGACGCGCTAGCTTGAGGTGTTAGTGTCCTTAC |
| 35 | CCGCGTAATACGACTCACTATAGGGTCGTTAGCTCAGTTGG |
| 36 | TGGTGGGTCGTGCAGGATT |
| 37 | CCGCGTAATACGACTCACTATAGGCTACGTAGCTCAGTTG |
| 38 | TGGTGGCTACGACGGGATTCG |
| 39 | CCGCGTAATACGACTCACTATAGCCCGGATAGCTCAGTC |
| 40 | TGGTGCCCGGACTCGGAA |
| 41 | CCGCGTAATACGACTCACTATAGGTGAGGTGTCCGAGTG |
| 42 | TGGCGGTGAGGGGGGGATTCG |
| 43 | AGGCGTGCTCCTTCAGCCACTCG |
| 44 | TGAAGGAGCACGCCTCGAAAGTGTGTATACGGCAAC |
| 45 | TGAAGGAGCACGCCTGCTAAGTGTGTATACGGCAAC |
| 46 | CCGCGTAATACGACTCACTATAGCTGATATGGCTCAGTTGG |
| 47 | TGGTGCTGATACCCAGAGTCG |
| 48 | AAGGGTGCGCTCTACCAACTGAGC |
| 49 | GTAGAGCGCACCCTTCGTAAGGGTGAGGTCCCCAG |
| 50 | CCGCGTAATACGACTCACTATAGGTGGGGTTCCCGAG |
| 51 | TGGTGGTGGGGGAAGGATTCG |
| 52 | CCGCGTAATACGACTCACTATAGCGTCCGTAGCTCAGTTG |
| 53 | TGGTGCGTCCGAGTGGACTCG |
| 54 | AAGGTGGTGCTCTAACCAACTGAGCTAC |
| 55 | TTAGAGCACCACCTTCACATGGTGGGGGTCGG |
| 56 | GAGATTAATACGACTCACTATAGGGCACGTAGCGCAGCCTGG |

| | |
|---|---|
| 57 | TGGTCGGCACGAGAGGATTT |
| 58 | ATGACGGTGCGCTACCAGGCTGC |
| 59 | GTAGCGCACCGTCATCGGGTGTCGGGGG |
| 60 | GGAGAGCGCTTGCATCAAATGCAAGAGGTCAG |
| 61 | GGAGAGCGCTTGCATCGAATGCAAGAGGTCAG |
| 62 | GGAGAGCGCTTGCATCAGATGCAAGAGGTCAG |
| 63 | GGAGAGCGCTTGCATCGGATGCAAGAGGTCAG |
| 64 | GGAGAGCGCTTGCATCCGATGCAAGAGGTCAG |
| 65 | GGAGAGCGCTTGCATGCTATGCAAGAGGTCAG |
| 66 | GGAGAGCGCTTGCATCGTATGCAAGAGGTCAG |
| 67 | GGAGAGCGCTTGCATCACATGCAAGAGGTCAG |
| 68 | GGAGAGCGCTTGCATCCCATGCAAGAGGTCAG |
| 69 | TGAAGGAGCACGCCTCAAAAGTGTGTATACGGCAAC |
| 70 | TGAAGGAGCACGCCTCAGAAGTGTGTATACGGCAAC |
| 71 | TGAAGGAGCACGCCTCGGAAGTGTGTATACGGCAAC |
| 72 | TGAAGGAGCACGCCTCCGAAGTGTGTATACGGCAAC |
| 73 | TGAAGGAGCACGCCTCGTAAGTGTGTATACGGCAAC |
| 74 | TGAAGGAGCACGCCTCACAAGTGTGTATACGGCAAC |
| 75 | TGAAGGAGCACGCCTCGCAAGTGTGTATACGGCAAC |
| 76 | TGAAGGAGCACGCCTCCCAAGTGTGTATACGGCAAC |
| 77 | TAGACGCGCTAGCTTCGAGTGTTAGTGTCCTTAC |
| 78 | TAGACGCGCTAGCTTCGGGTGTTAGTGTCCTTAC |
| 79 | TAGACGCGCTAGCTTCCGGTGTTAGTGTCCTTAC |
| 80 | TAGACGCGCTAGCTTGCTGTGTTAGTGTCCTTAC |
| 81 | TAGACGCGCTAGCTTCGTGTGTTAGTGTCCTTAC |
| 82 | TAGACGCGCTAGCTTCACGTGTTAGTGTCCTTAC |
| 83 | TAGACGCGCTAGCTTCGCGTGTTAGTGTCCTTAC |
| 84 | TAGACGCGCTAGCTTCCCGTGTTAGTGTCCTTAC |
| 85 | GGAGAGCGCTTGCATUGCATGCAAGAGGTCAG |
| 86 | ATAGAGCGCTGCCCTTCGGAGGCAGAGG |
| 87 | ATAGAGCGCTGCCCTTCTGAGGCAGAGGTC |
| 88 | GAGATTAATACGACTCACTATAGGGGGTATCGCCAAGCGGTAAG |
| 89 | TGGCGGGGGTACGAGGATTCG |
| 90 | AATCCGGTGCCTTACCGCTTG |
| 91 | GTAAGGCACCGGATTTTGATTCCGGCATTCC |
| 92 | AGGGCGGTGTCCTGGGCCTC |
| 93 | CCAGGACACCGCCCTTTCACGGCGGTAAC |
| 94 | GTAGAGCACGACCTTTCCAAGGTCGGGGTC |
| 95 | GAGATTAATACGACTCACTATAGGGCTTGTAGCTCAGGTGGTTAG |
| 96 | TGGTAGGCCTGAGTGGACTTG |
| 97 | AGGGGTGCGCTCTAACCACC |
| 98 | TTAGAGCGCACCCCTTATAAGGGTGAGGTCG |
| 99 | TAGACGCGCTAGCTTTAAGTGTTAGTGTCCTTACG |
| 100 | TAGACGCGCTAGCTTTAGGTGTTAGTGTCCTTACG |
| 101 | AGTCAACTGCTCTACCAACTGAGC |
| 102 | GTAGAGCAGTTGACTTTTAATCAATTGGTCGC |
| 103 | GTAGCGCACCGTCATTGGGTGTCGGGGG |
| 104 | TGAAGGAGCACGCCTTGAAAGTGTGTATACGGCAAC |
| 105 | GTAGAGCGCACCCTTTGTAAGGGTGAGGTCCCCAG |
| 106 | TTAGAGCACCACCTTTACATGGTGGGGGTCGG |
| 107 | ACACTCTTTCCCTACACGACGCTCTTCCGATCTCGTAACCTGCTTCTTGGCGCAACTCTG |
| 108 | ACACTCTTTCCCTACACGACGCTCTTCCGATCTGAATTGGGGCTTCTTGGCGCAACTCTG |

| | |
|---|---|
| 109 | ACACTCTTTCCCTACACGACGCTCTTCCGATCTAACCACTCGCTTCTTGGCGCAACTCTG |
| 110 | ACACTCTTTCCCTACACGACGCTCTTCCGATCTCAGGGTATGCTTCTTGGCGCAACTCTG |
| 111 | ACACTCTTTCCCTACACGACGCTCTTCCGATCTTAGGAGCAGCTTCTTGGCGCAACTCTG |
| 112 | GTGACTGGAGTTCAGACGTGTGCTCTTCCGATCTGAGACTTCGGTCCTCGAGGTAAGAACCGTC |
| 113 | ACACTCTTTCCCTACACGACGCTCTTCCGATCTATCACTGCTTACAACGAGCGCGAGCTC |
| 114 | ACACTCTTTCCCTACACGACGCTCTTCCGATCTGAAGGCAATTACAACGAGCGCGAGCTC |
| 115 | ACACTCTTTCCCTACACGACGCTCTTCCGATCTCTCGAGAATTACAACGAGCGCGAGCTC |
| 116 | ACACTCTTTCCCTACACGACGCTCTTCCGATCTCGTCAACATTACAACGAGCGCGAGCTC |
| 117 | ACACTCTTTCCCTACACGACGCTCTTCCGATCTCAAGAGTCTTACAACGAGCGCGAGCTC |
| 118 | GTGACTGGAGTTCAGACGTGTGCTCTTCCGATCTAGGGTGAGCGGAGAAAGAGCTCCTCC |
| 119 | ACACTCTTTCCCTACACGACGCTCTTCCGATCTGGACTAGTGTTGAGGGTGAGGGTGAGG |
| 120 | ACACTCTTTCCCTACACGACGCTCTTCCGATCTCAGAGGAAGTTGAGGGTGAGGGTGAGG |
| 121 | ACACTCTTTCCCTACACGACGCTCTTCCGATCTCGCTTGATGTTGAGGGTGAGGGTGAGG |
| 122 | ACACTCTTTCCCTACACGACGCTCTTCCGATCTGGAAATCGGTTGAGGGTGAGGGTGAGG |
| 123 | ACACTCTTTCCCTACACGACGCTCTTCCGATCTAGTGCTCTGTTGAGGGTGAGGGTGAGG |
| 124 | GTGACTGGAGTTCAGACGTGTGCTCTTCCGATCTTCGATCTGGCGAATCCAGTGAAAAGGAACGTC |

### Table S3. tRNAs used in this study

| tRNA(or pre-tRNA) | Primer set for IVT template | Primer set for mutation | Sequence |
|---|---|---|---|
| tRNA$^{Ala}$GGC | 9, 10 | - | GGGGCUAUAGCUCAGCUGGGAGAGCGCUUGCAUGGCAUGCAAGAGGUCAGCGGUUCGAUCCCGCUUAGCUCCACCA |
| tRNA$^{Ala}$CGC | 9, 10 | 11,12 | GGGGCUAUAGCUCAGCUGGGAGAGCGCUUGCAUCGCAUGCAAGAGGUCAGCGGUUCGAUCCCGCUUAGCUCCACCA |
| tRNA$^{Arg}$CCG | 13,14 | - | GCGCCCGUAGCUCAGCUGGAUAGAGCGCUGCCCUCCGGAGGCAGAGGUCUCAGGUUCGAAUCCUGUCGGGCGCGCCA |
| tRNA$^{Arg}$GCG | 13,14 | 15,16 | GCGCCCGUAGCUCAGCUGGAUAGAGCGCUGCCCUGCGGAGGCAGAGGUCUCAGGUUCGAAUCCUGUCGGGCGCGCCA |
| tRNA$^{Arg}$CCU | 13,14 | 15,17 | GCGCCCGUAGCUCAGCUGGAUAGAGCGCUGCCCUCCUGAGGCAGAGGUCUCAGGUUCGAAUCCUGUCGGGCGCGCCA |
| tRNA$^{Asn}$GUC | - | - | UCCUCUGUAGUUCAGUCGGUAGAACGGCGGACUGUUAAUCCGUAUGUCACUGGUUCGAGUCCAGUCAGAGGAGCCA |
| tRNA$^{Asp}$GUC | 18,19 | - | GGAGCGGUAGUUCAGUCGGUUAGAAUACCUGCCUGUCACGCAGGGGGUCGCGGGUUCGAGUCCCGUCCGUUCCGCCA |
| tRNA$^{Cys}$GCA | 20,21 | - | GGCGCGUUAACAAAGCGGUUAUGUAGCGGAUUGCAAAUCCGUCUAGUCCGGUUCGACUCCGGAACGCGCCUCCA |
| tRNA$^{Gln}$CUG | - | - | UGGGGUAUCGCCAAGCGGUAAGGCACCGGAUUCUGAUUCCGGCAUUCCGAGGUUCGAAUCCUCGUACCCCAGCCA |
| tRNA$^{Gly}$GCC | 22,23 | - | GCGGGAAUAGCUCAGUUGGUAGAGCACGACCUUGCCAAGGUCGGGGUCGCGAGUUCGAGUCUCGUUUCCCGCUCCA |
| tRNA$^{Gly}$CCC | 22,23 | 24,25 | GCGGGAAUAGCUCAGUUGGUAGAGCACGACCUUCCCAAGGUCGGGGUCGCGAGUUCGAGUCUCGUUUCCCGCUCCA |
| tRNA$^{Glu}$CUC | 26,27 | - | GUCCCCUUCGUCUAGAGGCCCAGGACACCGCCCUCUCACGGCGGUAACAGGGGUUCGAAUCCCCUAGGGGACGCCA |
| tRNA$^{His}$GUG | 28,29 | - | GGUGGCUAUAGCUCAGUUGGUAGAGCCCUGGAUUGUGAUUCCAGUUGUCGUGGGUUCGAAUCCCAUUAGCCACCCCA |
| tRNA$^{Ile}$GAU | - | - | AGGCUUGUAGCUCAGGUGGUUAGAGCGCACCCCUGAUAAGGGUGAGGUCGGUGGUUCAAGUCCACUCAGGCCUACCA |
| tRNA$^{Leu}$CAG | 30,31 | - | GCGAAGGUGGCGGAAUUGGUAGACGCGCUAGCUUCAGGUGUUAGUGUCCUUACGGACGUGGGGGUUCAAGUCCCCCCCCUCGCACCA |
| tRNA$^{Leu}$CAA | 30,31 | 32,33 | GCGAAGGUGGCGGAAUUGGUAGACGCGCUAGCUUCAAGUGUUAGUGUCCUUACGGACGUGGGGGUUCAAGUCCCCCCCCUCGCACCA |
| tRNA$^{Leu}$GAG | 30,31 | 32,34 | GCGAAGGUGGCGGAAUUGGUAGACGCGCUAGCUUGAGGUGUUAGUGUCCUUACGGACGUGGGGGUUCAAGUCCCCCCCCUCGCACCA |
| tRNA$^{Lys}$CUU | 35,36 | - | GGGUCGUUAGCUCAGUUGGUAGAGCAGUUGACUCUUAAUCAAUUGGUCGCAGGUUCGAAUCCUGCACGACCCACCA |
| tRNA$^{fMet}$CAU | - | - | CGCGGGGUGGAGCAGCCUGGUAGCUCGUCGGGCUCAUAACCCGAAGAUCGUCGGUUCAAAUCCGGCCCCCGCAACCA |
| tRNA$^{mMet}$CAU | 37,38 | - | GGCUACGUAGCUCAGUUGGUUAGAGCACAUCACUCAUAAUGAUGGGGUCACAGGUUCGAAUCCCGUCGUAGCCACCA |
| tRNA$^{Phe}$GAA | 39,40 | - | GCCCGGAUAGCUCAGUCGGUAGAGCAGGGGAUUGAAAAUCCCCGUGUCCUUGGUUCGAUUCCGAGUCCGGGCACCA |
| tRNA$^{Ser}$GGA | 41,42 | - | GGUGAGGUGUCCGAGUGGCUGAAGGAGCACGCCUGGAAAGUGUGUAUACGGCAACGUAUCGGGGGUUCGAAUCCCCCCCUCACCGCCA |
| tRNA$^{Ser}$CGA | 41,42 | 43,44 | GGUGAGGUGUCCGAGUGGCUGAAGGAGCACGCCUCGAAAGUGUGUAUACGGCAACGUAUCGGGGGUUCGAAUCCCCCCCUCACCGCCA |
| tRNA$^{Ser}$GCU | 41,42 | 43,45 | GGUGAGGUGUCCGAGUGGCUGAAGGAGCACGCCUGCUAAGUGUGUAUACGGCAACGUAUCGGGGGUUCGAAUCCCCCCCUCACCGCCA |
| tRNA$^{Thr}$GGU | 46,47 | - | GCUGAUAUGGCUCAGUUGGUAGAGCGCACCCUUGGUAAGGGUGAGGUCCCCAGUUCGACUCUGGGUAUCAGCACCA |
| tRNA$^{Thr}$CGU | 46,47 | 48,49 | GCUGAUAUGGCUCAGUUGGUAGAGCGCACCCUUCGUAAGGGUGAGGUCCCCAGUUCGACUCUGGGUAUCAGCACCA |
| tRNA$^{Trp}$CCA | - | - | AGGGGCGUAGUUCAAUUGGUAGAGCACCGGUCUCCAAAACCGGGUGUUGGGAGUUCGAGUCUCUCCGCCCCUGCCA |
| tRNA$^{Tyr}$GUA | 50,51 | - | GGUGGGGUUCCCGAGCGGCCAAAGGGAGCAGACUGUAAAUCUGCCGUCACAGACUUCGAAGGUUCGAAUCCUUCCCCCACCACCA |
| tRNA$^{Val}$GAC | 52,53 | - | GCGUCCGUAGCUCAGUUGGUUAGAGCACCACCUUGACAUGGUGGGGGUCGGUGGUUCGAGUCCACUCGGACGCACCA |
| tRNA$^{Val}$CAC | 52,53 | 54,55 | GCGUCCGUAGCUCAGUUGGUUAGAGCACCACCUUCACAUGGUGGGGGUCGGUGGUUCGAGUCCACUCGGACGCACCA |
| tRNA$^{Pro}$GGG | - | - | CGGCACGUAGCGCAGCCUGGUAGCGCACCGUCAUGGGGUGUCGGGGGUCGGAGGUUCAAAUCCUCUCGUGCCGACCA |
| tRNA$^{Pro}$CGG | 56,57 | 58,59 | GGGCACGUAGCGCAGCCUGGUAGCGCACCGUCAUCGGGUGUCGGGGGUCGGAGGUUCAAAUCCUCUCGUGCCGACCA |
| tRNA$^{Ala}$CAA *anticodon | 9, 10 | 11, 60 | GGGGCUAUAGCUCAGCUGGGAGAGCGCUUGCAUCAAAUGCAAGAGGUCAGCGGUUCGAUCCCGCUUAGCUCCACCA |

| | | | |
|---|---|---|---|
| tRNA$^{Ala}$CGA | 9, 10 | 11, 61 | GGGGCUAUAGCUCAGCUGGGAGAGCGCUUGCAUCGAAUGCAAGAGGUCAGCGGUUCGAUCCCGCUUAGCUCCACCA |
| tRNA$^{Ala}$CAG | 9, 10 | 11, 62 | GGGGCUAUAGCUCAGCUGGGAGAGCGCUUGCAUCAGAUGCAAGAGGUCAGCGGUUCGAUCCCGCUUAGCUCCACCA |
| tRNA$^{Ala}$CGG | 9, 10 | 11, 63 | GGGGCUAUAGCUCAGCUGGGAGAGCGCUUGCAUCGGAUGCAAGAGGUCAGCGGUUCGAUCCCGCUUAGCUCCACCA |
| tRNA$^{Ala}$CCG | 9, 10 | 11, 64 | GGGGCUAUAGCUCAGCUGGGAGAGCGCUUGCAUCCGAUGCAAGAGGUCAGCGGUUCGAUCCCGCUUAGCUCCACCA |
| tRNA$^{Ala}$GCU | 9, 10 | 11, 65 | GGGGCUAUAGCUCAGCUGGGAGAGCGCUUGCAUGCUAUGCAAGAGGUCAGCGGUUCGAUCCCGCUUAGCUCCACCA |
| tRNA$^{Ala}$CGU | 9, 10 | 11, 66 | GGGGCUAUAGCUCAGCUGGGAGAGCGCUUGCAUCGUAUGCAAGAGGUCAGCGGUUCGAUCCCGCUUAGCUCCACCA |
| tRNA$^{Ala}$CAC | 9, 10 | 11, 67 | GGGGCUAUAGCUCAGCUGGGAGAGCGCUUGCAUCACAUGCAAGAGGUCAGCGGUUCGAUCCCGCUUAGCUCCACCA |
| tRNA$^{Ala}$CCC | 9, 10 | 11, 68 | GGGGCUAUAGCUCAGCUGGGAGAGCGCUUGCAUCCCAUGCAAGAGGUCAGCGGUUCGAUCCCGCUUAGCUCCACCA |
| tRNA$^{Ser}$CAA<br>*anticodon | 41, 42 | 43, 69 | GGUGAGGUGUCCGAGUGGCUGAAGGAGCACGCCUCAAAAGUGUGUAUACGGCAACGUAUCGGGGGUUCGAAUCCCCCCCUCACCGCCA |
| tRNA$^{Ser}$CAG | 41, 42 | 43, 70 | GGUGAGGUGUCCGAGUGGCUGAAGGAGCACGCCUCAGAAGUGUGUAUACGGCAACGUAUCGGGGGUUCGAAUCCCCCCCUCACCGCCA |
| tRNA$^{Ser}$CGG | 41, 42 | 43, 71 | GGUGAGGUGUCCGAGUGGCUGAAGGAGCACGCCUCGGAAGUGUGUAUACGGCAACGUAUCGGGGGUUCGAAUCCCCCCCUCACCGCCA |
| tRNA$^{Ser}$CCG | 41, 42 | 43, 72 | GGUGAGGUGUCCGAGUGGCUGAAGGAGCACGCCUCCGAAGUGUGUAUACGGCAACGUAUCGGGGGUUCGAAUCCCCCCCUCACCGCCA |
| tRNA$^{Ser}$CGU | 41, 42 | 43, 73 | GGUGAGGUGUCCGAGUGGCUGAAGGAGCACGCCUCGUAAGUGUGUAUACGGCAACGUAUCGGGGGUUCGAAUCCCCCCCUCACCGCCA |
| tRNA$^{Ser}$CAC | 41, 42 | 43, 74 | GGUGAGGUGUCCGAGUGGCUGAAGGAGCACGCCUCACAAGUGUGUAUACGGCAACGUAUCGGGGGUUCGAAUCCCCCCCUCACCGCCA |
| tRNA$^{Ser}$CGC | 41, 42 | 43, 75 | GGUGAGGUGUCCGAGUGGCUGAAGGAGCACGCCUCGCAAGUGUGUAUACGGCAACGUAUCGGGGGUUCGAAUCCCCCCCUCACCGCCA |
| tRNA$^{Ser}$CCC | 41, 42 | 43, 76 | GGUGAGGUGUCCGAGUGGCUGAAGGAGCACGCCUCCCAAGUGUGUAUACGGCAACGUAUCGGGGGUUCGAAUCCCCCCCUCACCGCCA |
| tRNA$^{Leu}$CGA<br>*anticodon | 30, 31 | 32, 77 | GCGAAGGUGGCGGAAUUGGUAGACGCGCUAGCUUCGAGUGUUAGUGUCCUUACGGACGUGGGGGUUCAAGUCCCCCCCUCGCACCA |
| tRNA$^{Leu}$CGG | 30, 31 | 32, 78 | GCGAAGGUGGCGGAAUUGGUAGACGCGCUAGCUUCGGGUGUUAGUGUCCUUACGGACGUGGGGGUUCAAGUCCCCCCCUCGCACCA |
| tRNA$^{Leu}$CCG | 30, 31 | 32, 79 | GCGAAGGUGGCGGAAUUGGUAGACGCGCUAGCUUCCGGUGUUAGUGUCCUUACGGACGUGGGGGUUCAAGUCCCCCCCUCGCACCA |
| tRNA$^{Leu}$GCU | 30, 31 | 32, 80 | GCGAAGGUGGCGGAAUUGGUAGACGCGCUAGCUUGCUGUGUUAGUGUCCUUACGGACGUGGGGGUUCAAGUCCCCCCCUCGCACCA |
| tRNA$^{Leu}$CGU | 30, 31 | 32, 81 | GCGAAGGUGGCGGAAUUGGUAGACGCGCUAGCUUCGUGUGUUAGUGUCCUUACGGACGUGGGGGUUCAAGUCCCCCCCUCGCACCA |
| tRNA$^{Leu}$CAC | 30, 31 | 32, 82 | GCGAAGGUGGCGGAAUUGGUAGACGCGCUAGCUUCACGUGUUAGUGUCCUUACGGACGUGGGGGUUCAAGUCCCCCCCUCGCACCA |
| tRNA$^{Leu}$CGC | 30, 31 | 32, 83 | GCGAAGGUGGCGGAAUUGGUAGACGCGCUAGCUUCGCGUGUUAGUGUCCUUACGGACGUGGGGGUUCAAGUCCCCCCCUCGCACCA |
| tRNA$^{Leu}$CCC | 30, 31 | 32, 84 | GCGAAGGUGGCGGAAUUGGUAGACGCGCUAGCUUCCCGUGUUAGUGUCCUUACGGACGUGGGGGUUCAAGUCCCCCCCUCGCACCA |
| tRNA$^{Ala}$UGC | 9, 10 | 11, 85 | GGGGCUAUAGCUCAGCUGGGAGAGCGCUUGCAUUGCAUGCAAGAGGUCAGCGGUUCGAUCCCGCUUAGCUCCACCA |
| tRNA$^{Arg}$UCG | 13,14 | 15, 86 | GCGCCCGTAGCTCAGCTGGATAGAGCGCTGCCCTUCGGAGGCAGAGGTCTCAGGTTCGAATCCTGTCGGGCGCGCCA |
| tRNA$^{Arg}$UCU | 13,14 | 15, 87 | GCGCCCGTAGCTCAGCTGGATAGAGCGCTGCCCTUCUGAGGCAGAGGTCTCAGGTTCGAATCCTGTCGGGCGCGCCA |
| tRNA$^{Gln}$UUG | 88, 89 | 90, 91 | TGGGGTATCGCCAAGCGGTAAGGCACCGGATTTTGATTCCGGCATTCCGAGGTTCGAATCCTCGTACCCCAGCCA |
| tRNA$^{Glu}$UUC | 26,27 | 92, 93 | GTCCCCTTCGTCTAGAGGCCCAGGACACCGCCCTTTCACGGCGGTAACAGGGGTTCGAATCCCCTAGGGGACGCCA |
| tRNA$^{Gly}$UCC | 22,23 | 24, 94 | GCGGGAATAGCTCAGTTGGTAGAGCACGACCTTUCCAAGGTCGGGGTCGCGAGTTCGAGTCTCGTTTCCCGCTCCA |
| tRNA$^{Ile}$UAU | 95, 96 | 97, 98 | AGGCTTGTAGCTCAGGTGGTTAGAGCGCACCCCTTATAAGGGTGAGGTCGGTGGTTCAAGTCCACTCAGGCCTACCA |
| tRNA$^{Leu}$UAA | 30,31 | 32, 99 | GCGAAGGUGGCGGAAUUGGUAGACGCGCUAGCUUUAAGUGUUAGUGUCCUUACGGACGUGGGGGUUCAAGUCCCCCCCUCGCACCA |
| tRNA$^{Leu}$UAG | 30,31 | 32, 100 | GCGAAGGUGGCGGAAUUGGUAGACGCGCUAGCUUUAGGUGUUAGUGUCCUUACGGACGUGGGGGUUCAAGUCCCCCCCUCGCACCA |
| tRNA$^{Lys}$UUU | 35,36 | 101, 102 | GGGTCGTTAGCTCAGTTGGTAGAGCAGTTGACTTTTAATCAATTGGTCGCAGGTTCGAATCCTGCACGACCCACCA |
| tRNA$^{Pro}$UGG | 56,57 | 58, 103 | CGGCACGTAGCGCAGCCTGGTAGCGCACCGTCATUGGGTGTCGGGGGTCGGAGGTTCAAATCCTCTCGTGCCGACCA |
| tRNA$^{Ser}$UGA | 41, 42 | 43, 104 | GGUGAGGUGUCCGAGUGGCUGAAGGAGCACGCCUUGAAAGUGUGUAUACGGCAACGUAUCGGGGGUUCGAAUCCCCCCCUCACCGCCA |

| | | | |
|---|---|---|---|
| $tRNA^{Thr}UGU$ | 46, 47 | 48, 105 | GCTGATATGGCTCAGTTGGTAGAGCGCACCCTTUGUAAGGGTGAGGTCCCCAGTTCGACTCTGG GTATCAGCACCA |
| $tRNA^{Val}UAC$ | 52, 53 | 54, 106 | GCGTCCGTAGCTCAGTTGGTTAGAGCACCACCTTUACATGGTGGGGGTCGGTGGTTCGAGTCCAC TCGGACGCACCA |

**Table S4. Weights for nucleotide substitution probabilities**

| | First base | Second base | Third base |
|---|---|---|---|
| For transitions | 1 | 0.5 | 1 |
| For transversions | 0.5 | 0.1 | 1 |

**Table S5. Physicochemical property values of amino acids used for cost evaluation**

| Amino acid | Polar Requirement (PR) | Molecular Volume (MV) | Hydropathy Index (HI) |
|---|---|---|---|
| Ala | 7 | 31 | 1.8 |
| Arg | 9.1 | 124 | -4.5 |
| Asp | 13 | 54 | -3.5 |
| Asn | 10 | 56 | -3.5 |
| Cys | 4.8 | 55 | 2.5 |
| Glu | 12.5 | 83 | -3.5 |
| Gln | 8.6 | 85 | -3.5 |
| Gly | 7.9 | 3 | -0.4 |
| His | 8.4 | 96 | -3.2 |
| Ile | 4.9 | 111 | 4.5 |
| Leu | 4.9 | 111 | 3.8 |
| Lys | 10.1 | 119 | -3.9 |
| Met | 5.3 | 105 | 1.9 |
| Phe | 5 | 132 | 2.8 |
| Pro | 6.6 | 32.5 | -1.6 |
| Ser | 7.5 | 32 | -0.8 |
| Thr | 6.6 | 61 | -0.7 |
| Trp | 5.2 | 170 | -0.9 |
| Tyr | 5.4 | 136 | -1.3 |
| Val | 5.6 | 84 | 4.2 |

**Table S6. Standard composition of the tRNA-free PURE system (tfPURE)**

| Component | Concentration | Component | Concentration |
|---|---|---|---|
| Initiation Factor 1 | 25 µM | Tryptophanyl-tRNA Synthetase | 28 nM |
| Initiation Factor 2 | 1.0 µM | Tyrosyl-tRNA Synthetase | 0.15 µM |
| Initiation Factor 3 | 4.9 µM | Valyl-tRNA Synthetase | 17 nM |
| Elongation Factor G | 1.1 µM | Methionyl-tRNA Formyltransferase | 0.59 µM |
| Elongation Factor Tu | 80 µM | Myokinase | 1.4 µM |
| Elongation Factor Ts | 3.3 µM | Creatine kinase | 0.25 µM |
| Release Factor 1 | 49 nM | Nucleoside diphosphate kinase | 16 nM |
| Release Factor 2 | 48 nM | Pyrophosphatase | 41 nM |
| Release Factor 3 | 0.17 µM | Trigger Factor | 1.0 µM |
| Ribosome Recycling Factor | 3.9 µM | E. coli DEAH type RNA helicase A | 63 nM |
| Alanyl-tRNA Synthetase | 0.73 µM | Ribosome | 1.0 µM |
| Arginyl-tRNA Synthetase | 31 nM | Tyrosine | 0.30 mM |
| Asparaginyl-tRNA Synthetase | 0.42 µM | Cysteine | 0.30 mM |
| Asparagyl-tRNA Synthetase | 0.12 µM | 18 other amino acids | 0.36 mM |
| Cysteinyl-tRNA Synthetase | 24 nM | ATP | 0.38 mM |
| Glutaminyl-tRNA Synthetase | 60 nM | GTP | 0.25 mM |
| Glutamyl-tRNA Synthetase | 0.23 µM | CTP | 0.13 mM |
| Glycyl-tRNA Synthetase | 86 nM | UTP | 0.13 mM |
| Histidyl-tRNA Synthetase | 85 nM | N-2-hydroxyethylpiperazine-N'-2-ethanesulfonic acid (pH7.6) | 0.10 M |
| Isoleucyl-tRNA Synthetase | 0.37 µM | Glutamic acid potassium salt | 70 mM |
| Leucyl-tRNA Synthetase | 41 nM | Spermidine | 0.375 mM |
| Lysyl-tRNA Synthetase | 0.12 µM | Creatine phosphate | 25 mM |
| Methionyl-tRNA Synthetase | 0.11 µM | Dithiothreitol | 6 mM |
| Phenylalanyl-tRNA Synthetase | 0.13 µM | 10-formyl-5,6,7,8-tetrahydro folic acid | 10 µg/mL |
| Prolyl-tRNA Synthetase | 0.17 µM | Yeast inorganic pyrophosphatase (NEB) | 0.2 units/mL |
| Seryl-tRNA Synthetase | 78 nM | RNase Plus Inhibitor (Promega) | 0.1 U/µL |
| Threonyl-tRNA Synthetase | 84 nM | T7 RNAP (Takara) | 1.7 U/µL |

### Table S7. tRNA composition in each genetic code

| tRNA (ng/ μL) | MGC | near-SGC(RV) | SGC | Code1 | Code2 | Code3 | Code4 | Code5 | Code6 | Code7 | Code8 | Code9 | Code10 |
|---|---|---|---|---|---|---|---|---|---|---|---|---|---|
| $tRNA^{Ala}$GGC | 12 | 12 | 12 | 12 | 12 | 12 | 12 | 12 | 12 | 12 | 12 | 12 | 12 |
| $tRNA^{Ala}$CGC | - | 12 | 12 | 12 | - | 12 | - | 12 | - | - | - | - | 12 |
| $tRNA^{Arg}$CCG | - | 12 | 12 | - | - | - | - | - | - | - | - | - | - |
| $tRNA^{Arg}$GCG | 12 | 12 | 12 | 12 | 12 | 12 | 12 | 12 | 12 | 12 | 12 | 12 | 12 |
| $tRNA^{Arg}$CCU | - | 100 | 100 | 12 | 12 | 12 | 12 | 12 | 12 | 12 | 12 | 12 | 12 |
| $tRNA^{Asp}$GUC | 12 | 12 | 12 | 12 | 12 | 12 | 12 | 12 | 12 | 12 | 12 | 12 | 12 |
| $tRNA^{Cys}$GCA | 12 | 12 | 12 | 12 | 12 | 12 | 12 | 12 | 12 | 12 | 12 | 12 | 12 |
| $tRNA^{Gly}$GCG | 12 | 12 | 12 | 12 | 12 | 12 | 12 | 12 | 12 | 12 | 12 | 12 | 12 |
| $tRNA^{Gly}$CCC | - | 12 | 12 | 12 | 12 | 12 | 12 | 12 | 12 | 12 | 12 | 12 | 12 |
| $tRNA^{Glu}$CUC | 100 | 100 | 100 | 100 | 100 | 100 | 100 | 100 | 100 | 100 | 100 | 100 | 100 |
| $tRNA^{His}$GUG | 12 | 12 | 12 | 12 | 12 | 12 | 12 | 12 | 12 | 12 | 12 | 12 | 12 |
| $tRNA^{Leu}$CAG | - | 12 | 12 | - | - | - | - | - | - | 12 | - | 12 | - |
| $tRNA^{Leu}$CAA | - | 12 | 12 | - | - | 12 | - | 12 | - | 12 | - | 12 | - |
| $tRNA^{Leu}$GAG | 12 | 12 | 12 | 12 | 12 | 12 | 12 | 12 | 12 | 12 | 12 | 12 | 12 |
| $tRNA^{Lys}$CUU | 12 | 12 | 12 | 12 | 12 | 12 | 12 | 12 | 12 | 12 | 12 | 12 | 12 |
| $tRNA^{mMet}$CAU | 12 | 12 | 12 | 12 | 12 | 12 | 12 | 12 | 12 | 12 | 12 | 12 | 12 |
| $tRNA^{Phe}$GAA | 12 | 12 | 12 | 12 | 12 | 12 | 12 | 12 | 12 | 12 | 12 | 12 | 12 |
| $tRNA^{Ser}$GGA | 12 | 12 | 12 | 12 | 12 | 12 | 12 | 12 | 12 | 12 | 12 | 12 | 12 |
| $tRNA^{Ser}$CGA | - | 12 | 12 | - | - | - | - | - | - | - | - | 12 | 12 |
| $tRNA^{Ser}$GCU | - | 12 | 12 | - | - | - | 12 | - | - | - | - | - | 12 |
| $tRNA^{Thr}$GGU | 12 | 12 | 12 | 12 | 12 | 12 | 12 | 12 | 12 | 12 | 12 | 12 | 12 |
| $tRNA^{Thr}$CGU | - | 12 | 12 | - | - | - | - | - | - | - | - | - | - |
| $tRNA^{Tyr}$GUA | 12 | 12 | 12 | 12 | 12 | 12 | 12 | 12 | 12 | 12 | 12 | 12 | 12 |
| $tRNA^{Val}$GAC | 12 | 12 | 12 | 12 | 12 | 12 | 12 | 12 | 12 | 12 | 12 | 12 | 12 |
| $tRNA^{Val}$CAC | - | 100 | 100 | - | - | - | - | - | - | - | - | - | - |
| $tRNA^{Pro}$CGG | - | 100 | 100 | - | - | - | - | - | - | - | - | - | - |
| $tRNA^{Pro}$GGG | 100 | 100 | 100 | 100 | 100 | 100 | 100 | 100 | 100 | 100 | 100 | 100 | 100 |
| $tRNA^{Ile}$GAU | 100 | 100 | 100 | 100 | 100 | 100 | 100 | 100 | 100 | 100 | 100 | 100 | 100 |
| $tRNA^{Asn}$GUU | 100 | 100 | 100 | 100 | 100 | 100 | 100 | 100 | 100 | 100 | 100 | 100 | 100 |
| $tRNA^{Gln}$CUG | 12 | 12 | 12 | 12 | 12 | 12 | 12 | 12 | 12 | 12 | 12 | 12 | 12 |
| $tRNA^{Trp}$CCA | 12 | 12 | 12 | 12 | 12 | 12 | 12 | 12 | 12 | 12 | 12 | 12 | 12 |
| $tRNA^{fMet}$CAU | 12 | 12 | 12 | 12 | 12 | 12 | 12 | 12 | 12 | 12 | 12 | 12 | 12 |
| $tRNA^{Ala}$UGC | - | - | 12 | - | - | - | - | - | - | - | - | - | - |
| $tRNA^{Arg}$UCG | - | - | 12 | - | - | - | - | - | - | - | - | - | - |
| $tRNA^{Arg}$UCU | - | - | 12 | - | - | - | - | - | - | - | - | - | - |
| $tRNA^{Gln}$UUG | - | - | 12 | - | - | - | - | - | - | - | - | - | - |
| $tRNA^{Glu}$UUC | - | - | 100 | - | - | - | - | - | - | - | - | - | - |
| $tRNA^{Gly}$UCC | - | - | 12 | - | - | - | - | - | - | - | - | - | - |
| $tRNA^{Ile}$UAU | - | - | 100 | - | - | - | - | - | - | - | - | - | - |

| | | | | | | | | | | | | | |
|---|---|---|---|---|---|---|---|---|---|---|---|---|---|
| tRNA$^{Leu}$UAA | - | - | 12 | - | - | - | - | - | - | - | - | - | - |
| tRNA$^{Leu}$UAG | - | - | 12 | - | - | - | - | - | - | - | - | - | - |
| tRNA$^{Lys}$UUU | - | - | 12 | - | - | - | - | - | - | - | - | - | - |
| tRNA$^{Pro}$UGG | - | - | 100 | - | - | - | - | - | - | - | - | - | - |
| tRNA$^{Ser}$UGA | - | - | 12 | - | - | - | - | - | - | - | - | - | - |
| tRNA$^{Thr}$UGU | - | - | 12 | - | - | - | - | - | - | - | - | - | - |
| tRNA$^{Val}$UAC | - | - | 12 | - | - | - | - | - | - | - | - | - | - |
| tRNA$^{Ala}$CAA | - | - | - | 40 | - | - | 40 | - | - | - | 40 | - | 40 |
| tRNA$^{Ala}$CGA | - | - | - | - | 60 | - | 60 | - | - | 60 | - | - | - |
| tRNA$^{Ala}$CAG | - | - | - | 12 | - | 12 | - | 12 | - | - | 12 | - | 12 |
| tRNA$^{Ala}$CGG | - | - | - | 12 | - | - | - | - | - | 12 | - | - | - |
| tRNA$^{Ala}$CCG | - | - | - | - | - | - | 12 | - | - | - | 12 | - | - |
| tRNA$^{Ala}$GCU | - | - | - | - | - | - | - | 12 | - | 12 | 12 | - | - |
| tRNA$^{Ala}$CGU | - | - | - | 60 | - | - | - | - | - | - | - | - | - |
| tRNA$^{Ala}$CAC | - | - | - | - | - | - | 12 | - | - | - | 12 | - | 12 |
| tRNA$^{Ser}$CAA | - | - | - | - | 12 | - | - | - | 12 | - | - | - | - |
| tRNA$^{Ser}$CAG | - | - | - | - | 12 | - | 12 | - | 12 | - | - | - | - |
| tRNA$^{Ser}$CGG | - | - | - | - | 12 | 12 | - | 12 | - | - | - | 12 | 12 |
| tRNA$^{Ser}$CCG | - | - | - | - | - | - | - | 12 | - | 12 | - | - | 12 |
| tRNA$^{Ser}$CGU | - | - | - | - | 80 | - | 80 | 80 | - | 80 | - | 80 | 80 |
| tRNA$^{Ser}$CAC | - | - | - | - | - | - | - | 12 | 12 | - | - | - | - |
| tRNA$^{Ser}$CGC | - | - | - | - | - | - | - | - | - | 12 | - | 12 | - |
| tRNA$^{Leu}$CGA | - | - | - | 80 | - | 80 | - | 80 | 80 | - | 80 | - | - |
| tRNA$^{Leu}$CGG | - | - | - | - | - | - | 12 | - | 12 | - | 12 | 12 | - |
| tRNA$^{Leu}$CCG | - | - | - | 12 | 12 | 12 | - | - | 12 | - | - | 12 | - |
| tRNA$^{Leu}$GCU | - | - | - | 80 | 80 | 80 | - | - | 80 | - | - | - | - |
| tRNA$^{Leu}$CGU | - | - | - | - | - | 12 | - | - | 12 | - | 12 | - | - |
| tRNA$^{Leu}$CAC | - | - | - | 12 | 12 | 12 | - | - | - | 12 | - | 12 | - |
| tRNA$^{Leu}$CGC | - | - | - | - | 12 | - | 12 | - | 12 | - | 12 | - | - |

**Table S8. Major statistical comparisons of NanoLuc translation efficiency in Fig. 1D**

| Comparison | p value (Tukey's HSD test) |
|---|---|
| MGC vs near-SGC | $2.45\times10^{-6}$ |
| near-SGC vs near-SGC (RV) | $1.04\times10^{-6}$ |
| MGC vs near-SGC (RV) | 0.9503 |
| SGC vs near-SGC (RV) | $2.44\times10^{-15}$ |

**Table S9. Welch's t-test for NanoLuc translation using variant tRNAs in Fig. 2**

| Ala | | | | | | | | | |
|---|---|---|---|---|---|---|---|---|---|
| Codon | UUG | UCG | CUG | CCG | CGG | ACG | AGC | GUG | GGG |
| p value | $1.12×10^{-3}$ | $1.98×10^{-3}$ | $1.15×10^{-3}$ | $1.36×10^{-3}$ | $1.49×10^{-3}$ | $1.44×10^{-3}$ | $1.94×10^{-3}$ | $1.50×10^{-3}$ | $1.80×10^{-3}$ |
| Ser | | | | | | | | | |
| Codon | UUG | CUG | CCG | CGG | ACG | GUG | GCG | GGG | |
| p value | $1.98×10^{-3}$ | $2.26×10^{-3}$ | $1.90×10^{-4}$ | $3.37×10^{-2}$ | $5.14×10^{-4}$ | $1.11×10^{-2}$ | $9.08×10^{-4}$ | 0.778 | |
| Leu | | | | | | | | | |
| Codon | UCG | CCG | CGG | AGC | ACG | GUG | GCG | GGG | |
| p value | $3.17×10^{-4}$ | $1.06×10^{-5}$ | $3.58×10^{-4}$ | $6.19×10^{-5}$ | $9.88×10^{-4}$ | $1.11×10^{-3}$ | $3.21×10^{-4}$ | $2.07×10^{-4}$ | |

**Table S10. Spearman correlation analysis for Fig. 5**

| Reporter | Cost metric | Spearman's ρ | P value |
|---|---|---|---|
| GAL | Cost_PR | 0.109 | 0.75 |
| GAL | Cost_MV | 0.100 | 0.770 |
| GAL | Cost_HI | 0.00 | 1.00 |
| Luc | Cost_PR | -0.036 | 0.915 |
| Luc | Cost_MV | 0.245 | 0.467 |
| Luc | Cost_HI | -0.055 | 0.873 |
| mSG | Cost_PR | -0.191 | 0.574 |
| mSG | Cost_MV | -0.018 | 0.958 |
| mSG | Cost_HI | -0.227 | 0.502 |
